# A dual-dimensional deconvolution environment for ZT Scan DIA in metabolomics

**DOI:** 10.1101/2025.08.20.671307

**Authors:** Yuki Matsuzawa, Kanako Tokiyoshi, Bujinlkham Buyantogtokh, Takaki Oka, Rana Yamamoto, Lu Deng, Gordana Ivosev, David Cox, Paul RS Baker, Anjali Chelur, Nic Bloomfield, Manami Takeuchi, Ushio Takeda, Mikiko Takahashi, Mayu Hasegawa, Junki Miyamoto, Jason Causon, Takeshi Harayama, Hiroshi Tsugawa

## Abstract

We present a scanning data-independent acquisition (DIA) strategy, ZT Scan DIA, combined with dual-dimensional tandem mass spectrometry spectral filtering and deconvolution along both the quadrupole and retention time axes to reconstruct compound-specific MS2 spectra from complex mixtures. This approach is particularly effective for hydrophilic metabolomics data, where spectral similarity-based annotation is widely used, increasing annotation rates by 119–193% compared with conventional data-dependent acquisition (DDA) and window-based DIA methods. In lipidomics, deconvolution improved annotation precision by removing contaminant product ions and enabled separate quantification of co-eluting isomers using MS2 chromatograms, although common diagnostic ions could also be erroneously removed. Nevertheless, optimization of analysis parameters minimized this negative effect. Furthermore, we developed a practical data processing pipeline in which raw ZT Scan DIA-MS2 chromatograms are directly used for isomer separation and MS2-based quantification, covering 1,393 and 3,020 molecules for human plasma and mouse liver tissues, respectively. All data processing steps, including direct import of vendor raw data, are supported in MS-DIAL.

## Introduction

Mass spectrometry (MS) is the gold standard for comprehensive profiling of small molecules. These molecules are collectively referred to as the metabolome, lipidome, or exposome, depending on the biological context^1–3^. The challenge of small-molecule profiling lies in their vast diversity of chemical structures and concentrations within living organisms. As of July 6, 2025, the Human Metabolome Database (HMDB) listed 248,138 compounds in human blood, with concentrations ranging from 1 pM for aldosterone to 32 mM for hydrogen carbonate—a dynamic range spanning over 10⁷-fold^4^. Capturing all of these molecules simultaneously remains unrealistic owing to the limited dynamic range of MS (10⁴–10⁵). Consequently, optimized extraction protocols and analytical conditions are employed and generate >200 terabytes of MS data, which is housed in public repositories^5–7^. Although large-scale machine learning applications increasingly benefit from rich and structured knowledge bases, as exemplified by recent advances in language models, the demand for highly reusable and scalable omics datasets to drive transformative progress in biology has increased^8, 9^.

Liquid chromatography (LC) coupled with electrospray ionization-based tandem MS (MS/MS) is the most widely used platform for untargeted metabolomics and lipidomics. Because molecular annotation is a prerequisite for biological interpretation, properties, such as retention time (RT), precursor *m/z*, isotope pattern, and product ion (MS2) spectra, were computationally leveraged to assign molecular formulae, classes, and structures^1^. Among these, MS2 spectra are the most informative descriptors for characterizing molecular substructures and are typically acquired using either data-dependent acquisition (DDA) or data-independent acquisition (DIA)^10^. DDA provides high-purity MS2 spectra using narrow precursor isolation windows (<1 Da), which facilitate more reliable metabolite annotation based on spectral similarity metrics, such as dot product or spectral entropy, against reference libraries. In contrast, DIA acquires comprehensive MS2 data for all detectable precursor ions across the RT and *m/z* dimensions, thereby enabling the MS2 spectral detection of low-abundance molecules and semi-quantification of structural isomers based on unique fragment ion chromatographic profiles. Another advantage of DIA-based metabolomics is the high completeness of MS2 data, which facilitates retrospective reanalysis after spectral library updates and can enhance the value of repository-scale search frameworks such as MASST and ReDU environments^9, 11, 12^. However, a major challenge of DIA-based untargeted metabolomics is the mathematical deconvolution of highly multiplexed MS2 spectra, in which contaminant ions decrease spectral similarity scores and increase false-negative rates^13–15^. Moreover, the accurate characterization of co-eluting structural isomers remains a pressing issue in lipidomics, requiring the optimal utilization of diagnostic fragment ions that define lipid classes and acyl chain compositions^16^. Suitably, in this study, we present a novel untargeted analysis framework based on ZT Scan DIA, where dual-dimensional MS2 spectral filtering and deconvolution along both the quadrupole and retention time axes were implemented in a mass spectrometry data analysis platform MS-DIAL to accelerate metabolomics and lipidomics studies.

## Results

The latest ZT Scan DIA platforms (ZT Scan DIA 2.0 and later) enable acquisition of MS2 spectral data over a broader *m/z* range than the first-generation ZT Scan DIA method, which is essential for untargeted metabolomics and lipidomics. ZT Scan DIA is a DIA method that employs a sliding scanning technique to continuously isolate precursor ions for MS2 data acquisition, in contrast to window-based scanning approaches, such as SWATH-DIA (**Supplementary Fig. 1**). The unique data structure generated by ZT Scan DIA enables a new concept in spectra interpretation by leveraging the MS2 signal traces across the quadrupole (Q1) data bins (**Fig. 1a**). MS2 spectra acquired with isolation windows ranging from 5–20 Da are distributed into Q1 bins centered on the precursor isolation range, where the Q1 bin resolution is automatically determined by the width of the isolation window ranging from 1.0 Da for narrow windows (5 Da) to 4.0 Da for wider windows (20 Da). During each scan cycle, the MS2 signals from a given analyte were recorded across multiple overlapping Q1 bins corresponding to the isolation window.

**Fig. 1.**
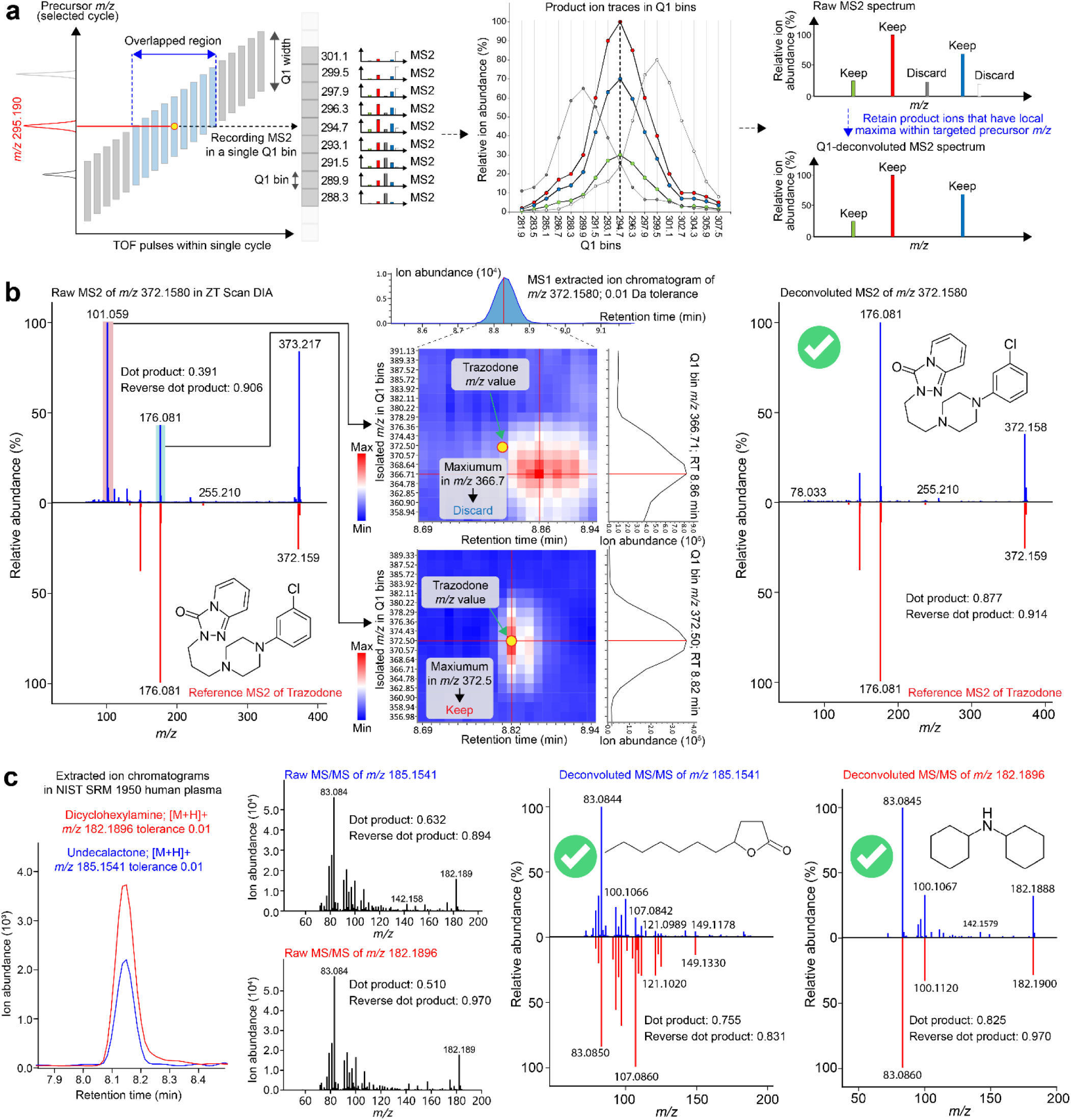
Concept and application of Q1-axis-based MS2 spectral filtering. (a) Schematic overview of Q1 axis-based data processing (Q1Dec), a spectrum filtering strategy that purifies product ion (MS2) spectra using ion intensity traces across Q1 bins. When a precursor isolation window of 7.5 Da is scanned across a precursor range, fragment ions of *m/z* 295.190 precursor ion are recorded across multiple Q1 bins spanning from *m/z* 287.69–302.69. Each Q1 bin has a width of 1.5 Da. Q1Dec retains product ions only if their intensity maxima fall within the Q1 bin of the target precursor or its immediate neighbors; all others are excluded. (b) Example of Q1Dec applied to trazodone. The raw MS2 spectrum (blue) includes contaminant signals, whereas the Q1Dec-processed spectrum aligns with the reference spectrum (red). A true fragment ion at *m/z* 176.081 shows its maximum intensity in the Q1 bin at *m/z* 372.50, consistent with the precursor *m/z* of protonated trazodone (theoretical *m/z* 372.1586). In contrast, the ion at *m/z* 101.059 is excluded as a contaminant, as its signal peaks in a mismatched Q1 bin (*m/z* 366.71). Heatmap axes represent scan time (x-axis) and Q1 bins (y-axis); red indicates higher ion intensity. (c) Application of Q1Dec to two co-eluted exposome compounds (γ-undecalactone and dicyclohexylamine) that share retention time but differ in terms of precursor *m/z* values. Raw MS2 spectra contain overlapping signals and background ions. Q1Dec resolves both compounds based on diagnostic fragments, enabling confident annotation with dot product and reverse dot product scores exceeding 0.755 and 0.831 for γ-undecalactone and 0.825 and 0.970 for dicyclohexylamine, respectively.

To reconstruct the compound-specific MS2 spectra, we developed a filtering algorithm termed Q1 axis-based deconvolution (Q1Dec). Q1Dec extracts MS2 ion chromatograms across Q1 bins and retains fragment ions whose intensity maxima fall within a user-defined Q1-bin tolerance around the precursor *m/z*. All other signals were discarded as contaminants. Following systematic evaluation of annotation performance at tolerances of ±0, ±1, ±2, and ±3 Q1 bins (described later), we set the default tolerance to ±1 bin for metabolomics and ±2 bins for lipidomics while allowing the Q1-bin tolerance to remain user-configurable. An illustrative example of compound detected thanks to Q1Dec is trazodone, which is a serotonin reuptake inhibitor used as an antidepressant. Notably, this compound is not currently listed among HMDB entries for human blood, although it has previously been reported by a targeted LC-MS/MS approach^17^. This example illustrates that accurate spectral deconvolution in untargeted workflows can reveal molecules that are not typically anticipated or included in targeted analyses (**Fig. 1b**). In this case, the ion at *m/z* 101.0585 was excluded because its signal peaked in the Q1 bin centered at *m/z* 366.71, which does not correspond to the precursor *m/z* of protonated trazodone (theoretical *m/z* 372.1586; Q1 bin centered at *m/z* 372.50). In contrast, the fragment ion at *m/z* 176.0808 was retained as its intensity maximum aligned with the correct Q1 bin. The Q1Dec-processed MS2 spectrum achieved dot product and reverse dot product similarity scores of 0.877 and 0.914, respectively. These values are substantially higher than those of the raw spectrum (0.391 and 0.906) and exceeded the commonly used 0.6–0.8 thresholds for confident annotation in untargeted metabolomics^18, 19^.

On the other hand, we observed that several low-abundance product ions remained after Q1Dec filtering, even though they would be expected to be removed based on the reference MS2 patterns. We therefore investigated the effect of product ion abundance on Q1Dec filtering performance (**Supplementary Fig. 2**; see **Methods** for the evaluation methodology). By comparing raw, Q1Dec-filtered, and reference MS2 spectra, product ions were classified into true positives (TP), true negatives (TN), false positives (FP), and false negatives (FN), according to whether they were retained or removed and whether they were expected to be retained or removed. Among these categories, TP and FN were used as direct readouts of Q1Dec performance because they represent clear cases of successful retention and erroneous removal of true fragment ions, respectively. TN and FP cases were not discussed in detail because they are strongly influenced by co-eluting molecules within the same precursor range and therefore require more careful interpretation. TP cases, in which product ions that should be retained were correctly retained, tended to show higher intensities than FN cases, in which true fragment ions were incorrectly removed. This suggests that, in low-intensity spectral regions, random spike-like signals or signal fluctuations have a stronger influence, increasing the risk that an apparent local maximum is detected in an incorrect Q1 bin. In addition, FN events may occur for low-intensity fragment ions due to the higher influence of co-eluting compounds, as discussed in the later lipidomics section.

We further demonstrated the utility of Q1Dec by resolving the co-eluted exposome compounds γ-undecalactone and dicyclohexylamine, which share the same RT in hydrophilic metabolomics data from NIST SRM 1950 human plasma (**Fig. 1c**). These compounds have precursor *m/z* values of 185.154 and 182.190, respectively, differing by only ∼3 Da, whereas the precursor isolation window in the ZT Scan DIA method was set to 7.2 Da. A diagnostic ion at *m/z* 83.084 was observed in both compounds and was retained in the Q1Dec MS2 spectra as the local maxima for this fragment in the Q1 bins, corresponding to *m/z* 185.1541 and 182.1896, respectively. The annotation of dicyclohexylamine was successful because of the substantial improvement in spectral similarity, indicated by an increased dot product score from 0.510 (raw MS2) to 0.825 (Q1Dec MS2) when matched against the reference MS2 spectrum. In contrast, the product ion at *m/z* 100.107 derived from dicyclohexylamine was not excluded from the Q1Dec MS2 spectrum of γ-undecalactone owing to a low signal-to-noise ratio, which caused its local maximum to appear within the Q1 bin at *m/z* 185. Nevertheless, the Q1Dec spectrum was sufficiently cleaned for confident annotation of γ-undecalactone, as reflected in an improved dot product score of 0.755 compared to 0.632 using the raw MS2 spectrum.

We further evaluated the use of RT-based deconvolution with Q1 filtering (RTDec, originally termed MS2Dec) to refine the Q1Dec-processed spectra (**Fig. 2**)^13^. For example, we applied Q1Dec followed by RTDec to the MS2 spectrum of 6-gingerol (**Fig. 2a**). Q1Dec effectively removed the major contaminant ions at *m/z* 288.289. However, certain nonspecific fragment ions, such as *m/z* 133.099, which exhibited a local maximum in the Q1 bin of *m/z* 295.77, remained in the spectrum after Q1Dec alone. RTDec addresses this limitation by evaluating the similarity of the elution profiles. This approach compares the chromatographic trace of each ion against a model peak generated from a unique mass fragment trace specific to the target precursor. This layer of LC deconvolution successfully excludes ions arising from the solvent background and co-eluting species with distinct RTs, resulting in a more purified MS2 spectrum. The dot product score of the final annotated spectrum substantially improved from 0.274, which was obtained using Q1Dec alone, to 0.605.

**Fig. 2.**
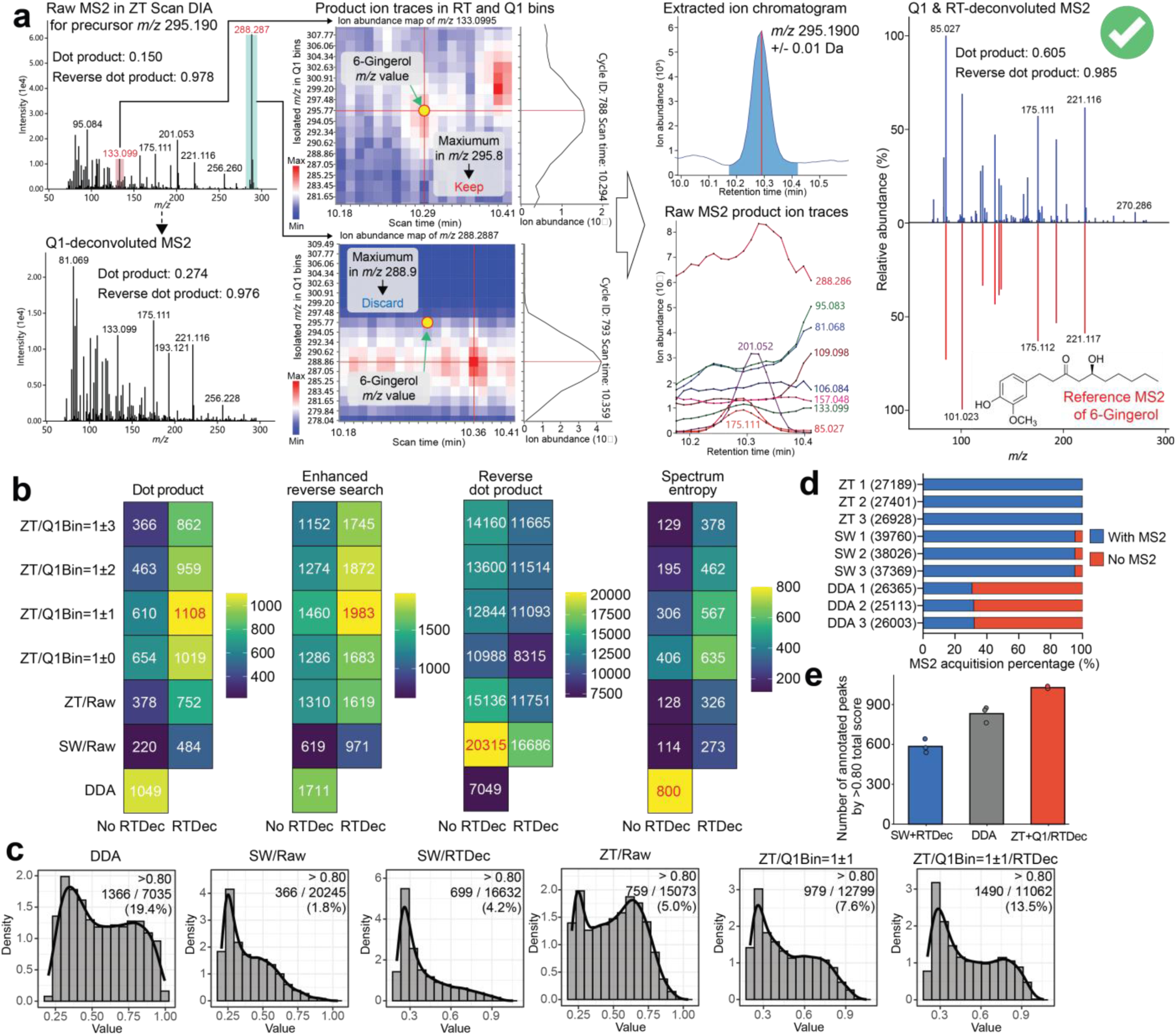
Validation of dual-dimensional MS2 processing using hydrophilic metabolomics data. (a) Example of Q1 axis-based spectral filtering (Q1Dec) and retention time-based deconvolution (RTDec) applied to the MS2 spectrum of 6-gingerol. A contaminant ion at *m/z* 288.287 was excluded owing to a mismatch between its Q1 bin maximum and the precursor *m/z* of 6-gingerol, whereas the fragment ion at *m/z* 133.099 was retained based on alignment with the correct Q1 bin at *m/z* 295.77. Subsequent RTDec applied to Q1Dec-processed MS2 chromatograms further purified the spectrum, thus increasing the dot product score from 0.150 (raw) and 0.274 (Q1Dec) to 0.605. (b) Number of peaks with similarity scores exceeding 0.80 for dot product, enhanced dot product, reverse dot product, and spectral entropy similarity in spectral-library matching. DDA, SW, and ZT denote data-dependent acquisition, SWATH-DIA, and ZT Scan DIA, respectively. For Q1Dec, different Q1-bin tolerances (±0, ±1, ±2, and ±3 bins) were evaluated, and RTDec was tested with and without application. (c) Distributions of MS-DIAL total scores for peaks with reverse dot product scores above 0.8. The x-axis represents the total score and the y-axis represents density. Results are shown for six configurations: DDA, original SWATH-DIA (SW/Raw), SWATH-DIA + RTDec (SW/RTDec), original ZT Scan DIA (ZT/Raw), ZT Scan DIA + Q1Dec (±1 bin) (ZT/Q1Bin=±1), and ZT Scan DIA + Q1Dec (±1 bin) + RTDec (ZT/Q1Bin=±1/RTDec). (d) Tandem mass spectrometry (MS/MS) assignment rates to precursor ions detected. Owing to spectral deconvolution processes, no MS2 data were acquired for several precursor ions, even for DIA methods. (e) Comparison of annotated peak numbers in untargeted metabolomics using DDA, SWATH-DIA + RTDec, and ZT Scan DIA + Q1Dec (±1 bin) + RTDec. The number of annotated features with MS-DIAL total scores exceeding 0.80 was counted for each method.

To systematically evaluate how dual-dimensional deconvolution affects spectral library search based metabolite annotation, we compared MS2 spectra generated by different acquisition and processing strategies using polar metabolomics data from NIST SRM 1950 human plasma acquired under identical LC conditions with DDA, SWATH-DIA (SW), and ZT Scan DIA (ZT) (three technical replicates each). For the DIA-based datasets, we examined the effects of RTDec and Q1Dec as well as different Q1-bin tolerances (±0, ±1, ±2, and ±3 bins). The resulting MS2 spectra were searched against MS2 reference libraries including MassBank, MoNA, and NIST20, and evaluated using dot product, reverse dot product, enhanced reverse search^20^, and spectral entropy similarity^21^ scores (**Fig. 2b**). We did not include MS2DeepScore^22^ or DreaMS^23^ in the benchmarking because their implementations would require deep-learning infrastructure that is not currently straightforward to integrate into the MS-DIAL runtime environment. For the conventional dot product score and the enhanced dot product score, which has recently been reported to outperform spectral entropy and machine-learning-based approaches such as MS2DeepScore, the ZT spectra processed with Q1-bin tolerance ±1 and RTDec produced the largest number of peaks with similarity scores exceeding 0.8. In contrast, for the reverse dot product metric, the SW spectra without RTDec produced the highest number of peaks with scores above 0.8. This result likely reflects the longer accumulation time of SW in addition to the fact that the MS2 spectra remain unfiltered. For the spectral entropy similarity score, DDA produced the largest number of peaks with scores above 0.8, followed by ZT processed with Q1-bin tolerance ±0 and RTDec, and then ZT processed with Q1-bin tolerance ±1 and RTDec. This trend is reasonable because the spectral entropy formulation was originally optimized using DDA spectra. We next evaluated the data using the total MS-DIAL score, which integrates dot product, reverse dot product, and spectral peak match percentage. Under this metric, the ZT spectra processed with Q1-bin tolerance ±1 and RTDec produced the largest number of peaks with scores above 0.8, with DDA ranking second (**Fig. 2c**). These results indicate that, for metabolite annotation based on spectral similarity searching, processing ZT data with Q1-bin tolerance ±1 and RTDec yields the highest number of annotated features. However, the number of false positives may also increase under these conditions, and a systematic evaluation using comprehensive standard mixtures would be required for a definitive assessment. At minimum, our results indicate that annotation reproducibility within individual samples is highest when ZT Scan DIA data are processed with Q1-bin tolerance ±1 and RTDec (**Fig. 2d and 2e**). In DDA, the MS2 acquisition coverage per sample was approximately 30–35%, whereas ZT processed with Q1-bin tolerance ±1 and RTDec achieved MS2 coverage exceeding 99.5% (**Fig. 2d**). The coverage does not reach 100% in DIA because some precursor features produce extremely weak MS2 signals or lose all fragment ions during the deconvolution process. In addition, the difference in the total number of detected peaks across acquisition methods partly reflects differences in MS1 accumulation time and cycle time. This higher MS2 coverage fundamentally changes the annotation behavior in the acquisition strategies. In DIA-based workflows, the number of annotated compounds rapidly reaches a plateau because MS2 information is available for nearly all precursor features within a single sample. In contrast, DDA-based workflows gradually accumulate annotations as the number of analyzed samples increases due to stochastic precursor selection^24^. This explains why DDA still produces competitive results in the present dataset (N = 3). However, when considering annotation yield from a single sample and the reproducibility of those annotations, ZT Scan DIA processed with Q1-bin tolerance ±1 and RTDec consistently outperforms DDA, highlighting the practical advantage of the ZT + Q1/RTDec strategy for spectral library searching based metabolite annotation (**Fig. 2e**).

Furthermore, we evaluated the advantages and disadvantages of spectral deconvolutions for ZT Scan DIA in untargeted lipidomics. The product ion at *m/z* 184.07 in positive ion mode is a well-known diagnostic fragment of the phosphocholine head group found in phosphatidylcholine (PC) and sphingomyelin (SM) species. Due to the large diversity of PC and SM molecular species that cover a broad range of retention times and *m/z*, this fragment ion broadly contaminates MS2 data of other lipid species. While the spectral filtering and/or deconvolution processes have the potential to reduce this contamination, there was the risk that this fragment ion is erroneously removed from PC/SM fragment ions by these processes. In analyzing the ZT Scan DIA preliminary data, we realized this issue by generating an ion heatmap of *m/z* 184.0726 across the RT range, in which PC 16:0_18:1, the most abundant phospholipid in blood and many mammalian cells and tissues^25–27^, was eluted in the ZT Scan DIA data from NIST SRM 1950 human plasma (**Supplementary Fig. 3**). The heatmap revealed that this diagnostic ion is broadly distributed across Q1 bins from *m/z* 750.57–770.78, with a local maximum centered at *m/z* 760.54, corresponding to the precursor ion *m/z* 760.5841 of PC 16:0_18:1 in the MS1 dimension. Importantly, the same *m/z* 184.07 fragment was also observed in the DDA spectra for the neighboring precursors *m/z* 768.5594 and 770.6047. The existence of *m/z* 184.0725 in the product ion spectrum from the precursor *m/z* 768.5594 was considered to be a contaminant fragment ion, as this peak corresponded to phosphatidylethanolamine (PE) 18:0_20:4. The contaminant fragment ion of *m/z* 184.07 may mislead the annotation of this precursor *m/z* 768.5594 toward PC 35:4 (theoretical *m/z* 768.5537; [M+H]^+^). In contrast, Q1Dec applied to ZT Scan DIA data successfully removed the *m/z* 184.0725 signal from the MS2 spectrum of *m/z* 768.5594, meaning that the Q1Dec filtering could reduce false positive annotations in the complex lipid mixture. However, the mis-elimination of *m/z* 184.0726 signal from the MS2 spectrum of *m/z* 770.6047 was also observed. According to the product ions of the corresponding peak in the negative ion mode, as well as the elution order of lipids inferred from recent lipidomics studies^28, 29^, this peak should have been annotated as PC O-36:3, indicating the risk that the Q1Dec filtering could incorrectly remove some of the diagnostic ions and increase false negatives in untargeted lipidomics. These preliminary observations revealed both advantages and disadvantages of Q1Dec filtering in lipidomics.

In order to establish the parameters that achieve the best balance between the advantages and disadvantages of ZT Scan DIA in lipidomics, we performed systematic evaluations of Q1Dec filtering and RTDec deconvolution in ZT Scan DIA analyses with five different Q1 window settings (**Fig. 3a**). True positives (TPs) and false positives (FPs) were defined using data dependent acquisition (DDA) MS2 data from NIST SRM 1950 plasma, whereas peaks that were not annotated by the MS-DIAL lipidomics workflow were classified as “not evaluated” (NE). FPs were assigned when lipid annotations violated retention time rules in reversed-phase LC-based lipidomics, where increasing carbon number leads to longer retention times, whereas increasing unsaturation results in shorter retention times. In addition, peaks that were annotated due to coincidental matches to so-called “bar-code” product ions, defined here as noise signals consisting of ions with similar intensities appearing at random *m/z* values in SCIEX instruments, were also labeled as false positives. To construct the DDA-based reference dataset, alignment peak features from three technical replicates (n = 3) were used. The resulting lists of TPs, FPs, and NEs derived from the DDA data were used as “reference annotation set” in this evaluation. For each ZT Scan DIA dataset (n = 1), processed with or without Q1Dec and RTDec under different Q1 window widths (5, 9, 13.5, 16.2, and 19.9 Da), detected peaks were classified into one of six categories: TP, FP, NE, true negative (TN), false negative (FN), or putative true positive (pTP) (**Fig. 3a**).

**Figure 3.**
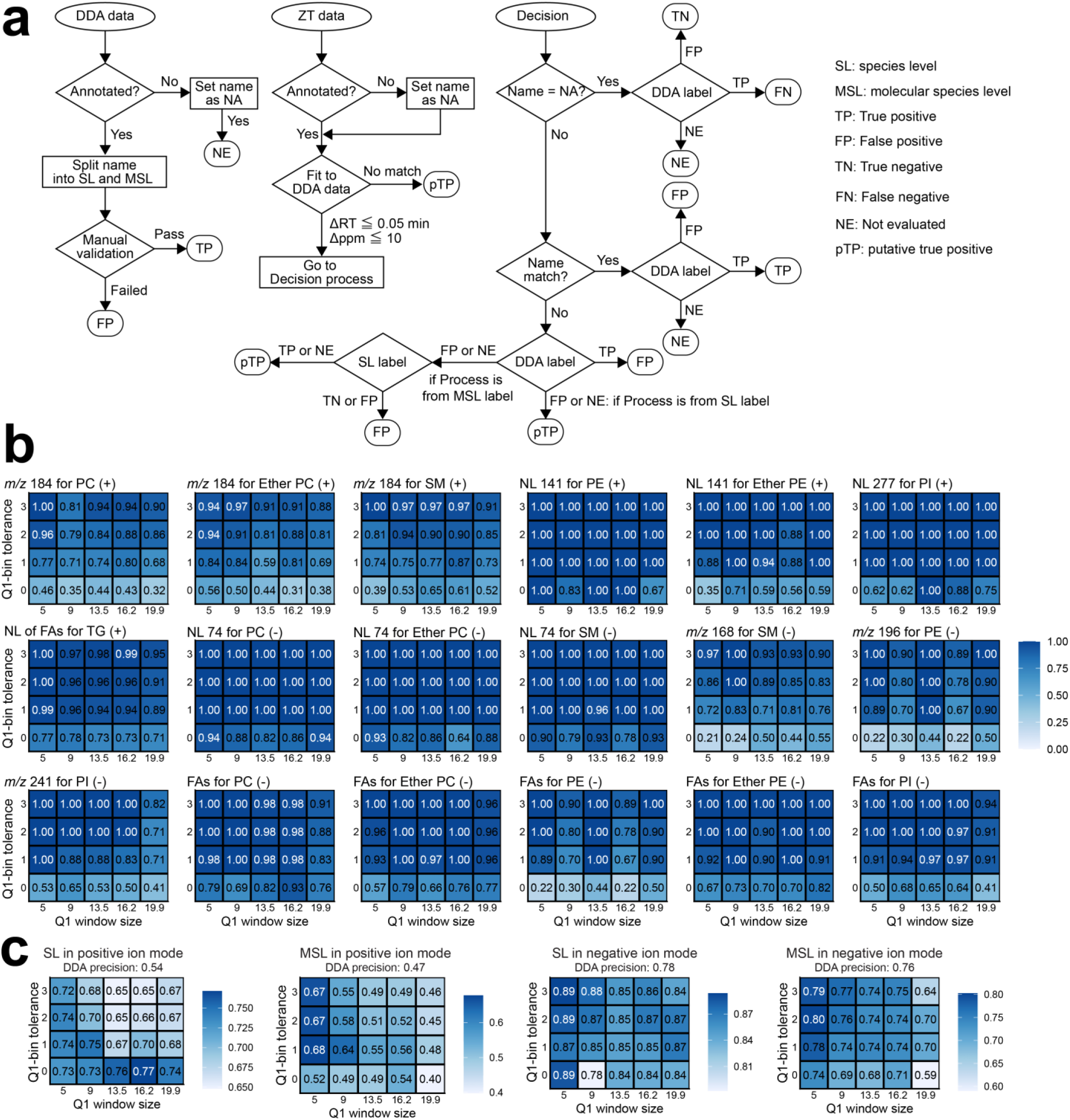
Systematic evaluation of Q1Dec effects on lipid annotation accuracy and diagnostic ion retention in ZT Scan DIA. (a) Workflow for classification of lipid annotation outcomes using DDA-derived reference annotation dataset and ZT Scan DIA results. DDA data were used to define true positives (TP), false positives (FP), and not evaluated (NE) peaks based on annotation results and retention time rules. ZT Scan DIA peaks were matched to DDA features within *m/z* (≤ 10 ppm) and retention time (≤ 0.05 min) tolerances, and classified into TP, FP, NE, true negative (TN), false negative (FN), or putative true positive (pTP) based on annotation consistency and label agreement at species level (SL) and molecular species level (MSL). Decision rules for classification are illustrated in the flowchart. (b) Recall of diagnostic fragment ions in ZT Scan DIA data acquired with different Q1 window sizes and processed under variable Q1-bin tolerance conditions. Recall was calculated as the fraction of ZT Scan DIA peaks matching DDA-derived true positives (Δ*m/z*≤ 10 ppm, ΔRT ≤ 0.05 min) in which the corresponding diagnostic ion was detected. The *m/z* tolerance for product ion matching was set to 0.01 Da. (c) Precision of peak annotation, as calculated by TP/(TP+FP), at the species level (SL) and molecular species level (MSL) in positive and negative ion modes. Precision was evaluated across different Q1 window sizes and Q1-bin tolerance conditions. The abbreviations of NL, PI, TG, FAs denote neutral loss, phosphatidylinositol, triacylglycerol, and fatty acyl related ions, respectively.

We first evaluated how Q1 window width and Q1-bin tolerance affect Q1Dec filtering for the retention of important diagnostic ions that are used for the annotations of major lipid species (**Fig. 3b**). For this analysis, we examined whether these diagnostic ions were present in the product ion spectra of ZT Scan DIA peaks that matched true positive (TP) peaks from the DDA dataset (reference annotation set) within *m/z* (< 0.01 Da) and retention time (< 0.05 min) tolerances. Since MS1 accumulation time was 200 ms in DDA, whereas it was 100 ms in ZT Scan DIA (i.e., half that of DDA), several peaks detected in DDA were not detected in ZT Scan DIA. Such peaks were excluded from this evaluation, since the objective was to evaluate deconvolution algorithms on shared peaks. The results indicated that tolerances of 0 or ±1 can cause severe mis-exclusion of diagnostic ions (**Fig. 3b**). Based on these results, setting the Q1-bin tolerance to ±2 or ±3 appeared to minimize the adverse effect of Q1Dec filtering, even for *m/z* 184, which showed the highest exclusion rate. However, increasing the Q1-bin tolerance also reduced the efficiency of DIA MS2 spectrum clean-up. Indeed, comparison of TPs and FPs defined using DDA data showed that a Q1-bin tolerance of ±2 provided higher precision, i.e., a lower false discovery rate, than wider tolerance settings (**Fig. 3c**). We therefore set Q1-bin tolerance of ±2 as a default for lipidomics analyses. Importantly, Q1Dec filtering improved annotation precision compared with DDA in both positive and negative ion modes, primarily by reducing false positives through removal of contaminant product ions.

When comparing Q1 windows, the 5 Da Q1 window yielded the lowest mis-exclusion rate. Notably, the *m/z* 184 fragment derived from PC O-36:3, which was removed in 7.2 Da Q1 window condition with ±1 Q1-bin tolerance in preliminary experiments (**Supplementary Fig. 3**), was not removed under the 5 Da Q1 window condition with ±2 Q1-bin tolerance. Under the Q1 window of 7.2 Da, the *m/z* 184 fragment derived from the M+3 isotope of PC 16:0_18:1 interfered with the Q1-bin trace of PC O-36:3. In contrast, under the 5 Da Q1 window condition, the isotope signal of PC 16:0_18:1 did not extend into the precursor *m/z* region of PC O-36:3. This observation indicates that reducing the Q1 window width can decrease the burden of deconvolution and help reduce false negative annotations. Of note, when using the Q1-bin tolerance of ±2 and a 5 Da Q1 window, the issue of mis-exclusion of fragment ions was almost exclusive to the *m/z* 184 diagnostic ion in positive mode and a few fatty acid fragments for ether PC in negative ion mode, showing that Q1Dec performance is maximized by optimizing parameters (**Fig. 3b**).

To visualize how mis-exclusion of the *m/z* 184 fragment occurs, we then overlaid the peak spots (defined by retention time and *m/z*) of curated PC, ether PC, and SM species detected in positive ion mode onto the *m/z* 184 ion chart obtained from the ZT Scan DIA data, with mis-excluded species shown as different symbols (**Fig. 4a**). This analysis showed that the mis-exclusion of the *m/z* 184 diagnostic ion was most likely to occur for low-abundance lipid peaks when a high-abundance co-eluting peak producing the same *m/z* 184 fragment was present in a similar retention time and precursor *m/z* region. In particular, for SM species, the corresponding PC species producing *m/z* 184 were often detected approximately 3 Da away, which falls just outside the ±2 Q1-bin tolerance used in this analysis. Thus, the reduced mis-exclusion of *m/z* 184 for SM at a ±3 Q1-bin tolerance (**Fig. 3b**) likely reflects the ability of this more permissive setting to include interference from nearby ions producing the same fragment, rather than an improvement in Q1Dec specificity. This analysis showed that fragment ion mis-exclusion was an explainable phenomenon, rather than a random failure of ZT Scan DIA and our deconvolution algorithms.

**Figure 4.**
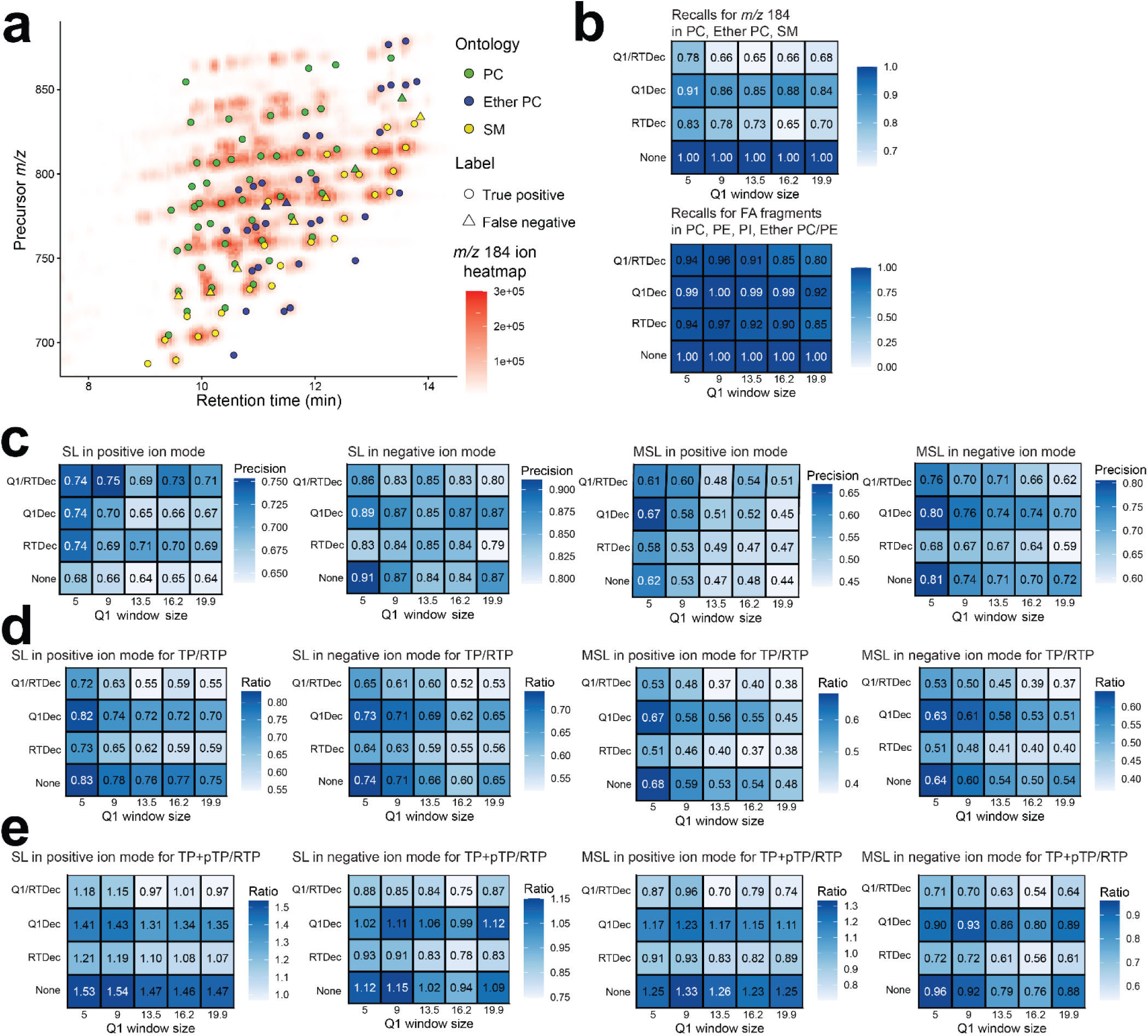
Effects of Q1Dec and RTDec on lipid annotation accuracy and diagnostic ion retention in ZT Scan DIA. (a) Overlay of PC, ether PC, and SM peak spots (defined by retention time + *m/z*) and the signal distribution of phosphocholine fragment ion (*m/z* 184) obtained by ZT Scan DIA with a 5 Da Q1 window size. Lipids that were correctly annotated after Q1Dec (TP for true positive) are shown as circles, while those not annotated due to the mis-exclusion of the fragment ion by Q1Dec are shown as triangles (FN for false negative). Note that most triangles are in proximity of a circle, showing that the common fragment ion was attributed to only one of them, due to the design of Q1Dec. (b) Recall of diagnostic fragment ions in ZT Scan DIA data processed under different conditions (no deconvolution, None; only RTDec used, RTDec; only Q1Dec used, Q1Dec; both Q1Dec and RTDec used, Q1/RTDec). Upper panel: recall of the phosphocholine fragment ion (*m/z* 184) from PC, ether PC, and SM in positive ion mode. Lower panel: recall of fatty acyl related fragment ions of PC, PE, PI, ether PC, and ether PE in negative ion mode. Recall was calculated as the fraction of ZT Scan DIA peaks matching DDA-derived true positives (*m/z* ≤ 10 ppm, ΔRT ≤ 0.05 min) in which the corresponding diagnostic ion was detected. (c) Precision TP/(TP+FP) at the species level (SL) and molecular species level (MSL) in positive and negative ion modes. Precision was evaluated across different processing conditions (None, RTDec, Q1Dec, and Q1Dec + RTDec). (d) Recall of reference annotation set’s true positives (TP/RTP) at SL and MSL levels in positive and negative ion modes, indicating the fraction of DDA-derived true positives recovered in ZT Scan DIA data. (e) Extended recall (TP+pTP)/RTP, incorporating putative true positives (pTP), at SL and MSL levels in positive and negative ion modes. This metric reflects both confirmed and plausibly correct annotations detected in ZT Scan DIA data.

With the Q1-bin tolerance fixed at ±2, we next evaluated whether RTDec affected the overall mis-exclusion rates of diagnostic ions, including the *m/z* 184 fragment derived from PC, ether PC, and SM in positive ion mode; fatty acyl related ions derived from PC, ether PC, PI, PE, and ether PE in negative ion mode; and other key diagnostic ions used for annotating major lipid classes (**Fig. 4b**; see **Supplementary Fig. 4** for individual results). In general, mis-exclusions increased when RTDec was applied, either with or without Q1Dec, and this effect became more pronounced as the Q1 window size increased. The *m/z* 184 fragment was affected most strongly, likely because its product ion chromatographic trace is distorted by the large number of co-isolated lipid molecules that generate this fragment, especially under wider isolation windows. RTDec also incorrectly excluded several fatty acyl related ions, indicating that this deconvolution step has a generally negative effect in lipidomics. Overall, these results indicate that mis-exclusion of important diagnostic ions can be minimized by acquiring ZT Scan DIA data with a 5 Da Q1 window and applying Q1Dec with a Q1-bin tolerance of ±2, without RTDec. This optimized condition reduced mis-exclusion to only a few SM species (**Fig. 3b** and **Fig. 4b**).

For each condition of deconvolution, we next calculated TP/(TP+FP), which reflects the rate of correctly assigned peaks among all annotated peaks (**Fig. 4c**). As described above, Q1Dec filtering consistently reduced false positives, thereby improving annotation precision. In contrast, RTDec increased the number of false negatives (FNs), resulting in decreased overall performance. We then calculated TP/RTP and (TP+pTP)/RTP, where RTP represents true positives in the reference annotation set derived from DDA data (N=3) (**Fig. 4d** and **4e**). TP/RTP (which has a maximum of 1 by definition) represents the coverage of lipids annotated by ZT Scan DIA among those annotated by DDA, while (TP+pTP)/RTP reflects the potential increase in annotation numbers by including peaks that were only annotated by ZT Scan DIA. Compared with the no-deconvolution condition, Q1Dec filtering led to a moderate decrease in both metrics, reflecting the slight increase in false negatives. This trend is consistent with the mis-exclusion of *m/z* 184 and fatty acyl related ions described above. Collectively, these results highlight several key findings. First, RTDec introduced an unacceptable increase in false negatives and provided limited benefit for lipidomics, where shared diagnostic ions are frequently observed in similar RT and *m/z* regions. Second, Q1Dec filtering slightly increased false negatives but effectively improved annotation accuracy. Third, Q1-bin tolerances of ±2 or ±3 bins with 5 Da Q1 window yielded the best overall performance. Finally, a single ZT Scan DIA dataset processed with Q1Dec yielded a similar or greater number of annotations than three replicate DDA datasets, although direct comparison is difficult because the number of detected peaks and product ion sensitivities differ among acquisition methods. Of note, because this systematic evaluation was conducted using a ZenoTOF 7600+ system rather than the ZenoTOF 8600 platform for which ZT Scan DIA was originally designed, important diagnostic fragment ions were sometimes missing owing to reduced accumulation time (thus explaining the relatively low TP/RTP values); therefore, the effects of Q1Dec and RTDec should be interpreted relative to the no-deconvolution condition.

In fact, mis-excluded ions can often be complemented by other diagnostic ions within the same spectrum or by MS2 spectral information acquired in a different ion mode. At the same time, to mitigate the risk of losing critical diagnostic ions, we implemented a visualization feature in MS-DIAL that simultaneously displays raw and deconvoluted MS2 spectra. This allows users to readily identify potential mis-exclusion events during data curation.

While Q1Dec filtering provides a clear advantage in lipidomics by reducing false positives, fully extracting the value of ZT Scan DIA data requires not only improving deconvolution accuracy but also enabling the use of MS2 information that remains as close as possible to the raw data. In particular, establishing a data analysis environment for MS2-based isomer separation and relative quantification is essential. We therefore investigated how MS2 fragment ions can be used to resolve co-eluting lipid structural isomers and expand lipid feature detection and quantification in ZT Scan DIA data. As an initial evaluation, we examined the correlation between the precursor ion peak heights and total ion counts of the corresponding MS2 spectra (**Fig. 5a**). Although certain abundant product ions, particularly PCs and triacylglycerols (TG), exhibited saturation relative to their MS1 precursor intensities, the overall correlation coefficient exceeded 0.89, indicating that MS2 fragment ions can also serve as quantitative measures. To facilitate the comprehensive use of MS2 data in metabolic profiling, we implemented new functions in MS-DIAL that allow quantification based on isomer-specific MS2 fragment ions when MS1 features are assigned multiple co-eluting isomers. For this, MS1 features that are defined by RT and precursor *m/z* were extended to multiple features that are defined by RT, precursor *m/z*, and isoform-specific MS2 fragments, whenever the peak was detected in all biological samples and if the lipid was annotated at the molecular species level (see **Methods** and **Supplementary Fig. 5**). This allowed the discrimination of co-eluting lipid isomers and their separate quantification based on MS2 fragment ion intensity. Applying this approach to aligned peak features from NIST SRM 1950 plasma and mouse liver tissue, we observed annotated feature count increases of 153 and 121%, respectively, compared with that of conventional two-dimensional feature detection (**Fig. 5b and 5c**). The total number of non-redundant annotated features reached 1,393 in the plasma and 3,020 in the liver. This surpassed the annotation coverage reported in our previous DDA-based study using a TripleTOF 6600 system, which characterized 541 and 746 lipid features in the same or similar biological samples, respectively^26^. Although direct comparisons of annotation results are limited by differences in acquisition mode and instrumentation, our results suggest that ZT Scan DIA, with three-dimensional feature detection, yields broader lipidome coverage than that of conventional untargeted lipidomic methods using standard extraction protocols^30^.

**Figure 5.**
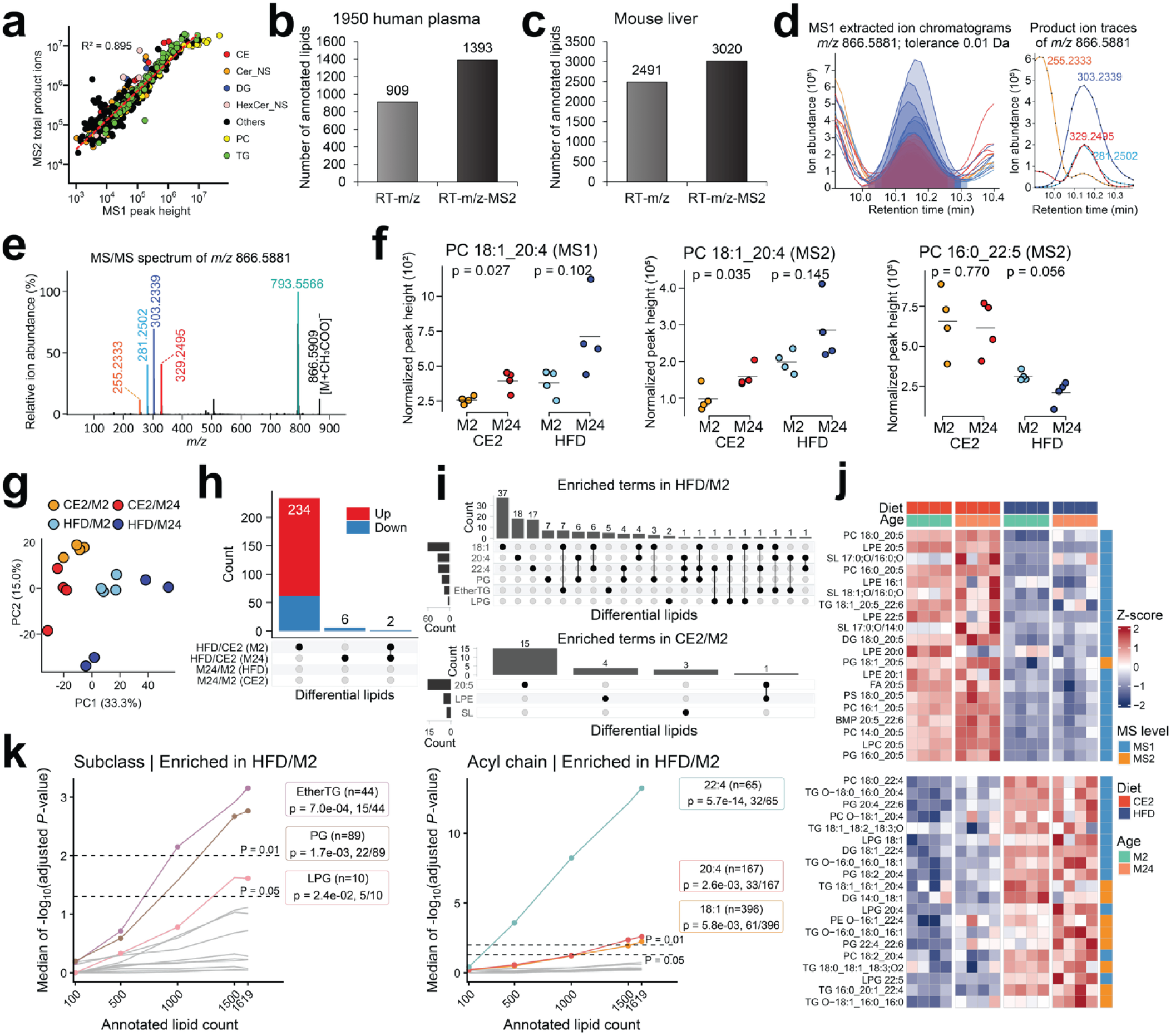
MS2 chromatogram–based lipidome profiling in ZT Scan DIA data processing. (a) Correlation between MS1- and MS2-based quantification values. Pearson correlation coefficients were calculated between MS1 precursor peak heights and the summed intensities of MS2 product ions. (b) Comparison of lipid feature counts using RT–*m/z* versus RT–*m/z*–MS2 dimensional criteria. Features were extracted from aligned data of NIST SRM 1950 plasma (N = 3). Positive and negative ion mode results were combined, and duplicate annotations were removed to count unique lipid features. (c) Same analysis as in (b), applied to mouse liver tissue data (four experimental conditions; N = 4). (d) Representative example of MS2 chromatogram–based separation of lipid structural isomers. Extracted ion chromatograms (EICs) of *m/z* 866.5881, initially annotated as PC 18:1_20:4, are shown across four mouse conditions (N = 4). MS2 chromatograms of diagnostic fragment ions (*m/z* 281.2502 and 303.2339 for PC 18:1_20:4; *m/z* 255.2333 and 329.2495 for PC 16:0_22:5) were used to resolve co-eluting isomers. (e) MS2 spectrum at the chromatographic peak apex corresponding to the feature shown in (d). (f) Comparison of normalized MS2-based quantification across biological groups. Product ion intensities specific to each structural isomer were summed. Statistical significance was evaluated using two-tailed Student’s *t*-test. (g) Principal component analysis (PCA) score plots generated from log-transformed, autoscaled lipid abundance data. (h) UpSet plot showing differentially expressed lipids (DELs), defined by adjusted *P* < 0.05 (two-sided Student’s *t*-test with Benjamini–Hochberg correction) and fold change ≥ 1.5 or ≤ 0.5 relative to control. (i) UpSet plot of enriched lipid terms identified by lipid molecule enrichment analysis. Statistical significance was defined as in (h). (j) Heatmap of DELs, highlighting representative lipids selected based on enrichment analysis results. Columns are grouped by dietary condition (CE2 or HFD) and age (M2 or M24), and rows indicate the MS level used for quantification (MS1 or MS2). (k) Relationship between the number of annotated lipids and statistical significance in enrichment analysis. The x-axis represents the number of lipid variables included in the analysis. For each subsampling condition, the proportion of significantly altered lipids was kept constant, and adjusted *P* values were evaluated. Numbers below each data point indicate the ratio of significant lipids to the total number of lipids within each category (significant lipids / total lipids in the category).

The advantage of MS2-based peak quantification was further exemplified by the separation of PC 18:1_20:4 and PC 16:0_22:5 isomers in the lipidomes of young (two-month-old) and aged (24-month-old) mice fed either a high-fat diet (HFD) or a control diet (CE2) for one month (**Fig. 5d**, **Fig. 5e, and Fig. 5f**). Both compounds were annotated at the molecular species level using MS-DIAL, with PC 18:1_20:4 selected as the representative annotation in the aligned peak set across the liver samples. By extracting unique fragment ion traces specific to each isomer, we uncovered distinct abundance patterns: PC 18:1_20:4 levels were elevated in aged mice across both diets, whereas PC 16:0_22:5 levels were reduced under HFD conditions and further decreased in aged mice fed the HFD. The accumulation of polyunsaturated fatty acid (PUFA)-containing phospholipids, such as PC 18:1_20:4, is a characteristic lipidomic signature of aging^25^. On the other hand, diet- and age-associated reductions in PC 16:0_22:5 levels have rarely been documented, while it is reported that n-3 PUFAs decline under HFD conditions, and 22:5 species in PC have been previously characterized as n-3 PUFA in the mouse liver using an oxygen attachment dissociation MS/MS based double bond position–resolved lipidomics technique^31–33^. Although the underlying mechanisms remain unclear, such isomer-specific trends may reflect changes in the activities of enzymes, such as fatty acid desaturases and acyltransferases. Notably, the use of MS2 chromatograms enabled the detection of distinct changes in PC 16:0_22:5 that were not captured by MS1-based quantification, primarily reflecting the behavior of PC 18:1_20:4.

Using a comprehensive lipidomics dataset of mouse liver tissues, we performed principal component analysis (PCA) on the auto-scaled data of log-transformed lipid abundances. This analysis revealed distinct score distributions among the four groups (CE2, young; CE2, aged; HFD, young; and HFD, aged) along the first (PC1) and second (PC2) principal component axes (**Fig. 5g**). We further compared the young and aged groups under the control diet condition by orthogonal partial least squares discriminant analysis (OPLS-DA) and examined lipids with variable importance in projection (VIP), which captured age-related differences (**Supplementary Fig. 6**). This analysis revealed an age-associated increase in bis(monoacylglycero)phosphate (BMP) and several glycerolipids containing PUFAs, consistent with a previous report^25^. Differentially expressed lipids (DEL) were also characterized by using Student’s t-tests with Benjamini–Hochberg correction (adjusted P < 0.05). This revealed that only a small number of DELs could be attributed to aging, due to the large inter-individual variability in lipid expression levels at 24-month-old mice. The most notable lipid signature captured in our lipidomics experiment was that HFD feeding in young mice induced substantial changes in the hepatic lipidome (234 lipid species), whereas HFD feeding in aged mice had almost no statistically significant effect on the liver lipid profiles (**Fig. 5h**).

Using the lipid names present in the liver lipidomics dataset and those identified as DELs in the CE2/M2 vs. HFD/M2 comparison, enrichment analysis of lipid classes and fatty acid properties revealed that, under HFD feeding, acyl chains such as monounsaturated fatty acids (18:1) and 20:4 (likely arachidonic acid), as well as lipid classes including phosphatidylglycerol (PG) and ether-linked TG (Ether TG) were extracted as the significantly enriched terms (**Fig. 5i**). The profile of the most abundant molecules within these enriched terms further showed that HFD feeding led to an increase in lipids containing saturated fatty acids (16:0, 18:0), monounsaturated fatty acids, and 20:4 (**Fig. 5j**). These trends closely reflected the fatty acid composition of the HFD used in this study (D12492; Research Diets, Inc.). In contrast, CE2 feeding uniquely enriched acyl chains such as 20:5 (likely eicosapentaenoic acid) and 22:6 (likely docosahexaenoic acid), along with lipid classes including PC, phosphatidylserine (PS), and PG. The higher abundance of lipids containing 20:5 under CE2 feeding compared with HFD is likely attributable to the absence of eicosapentaenoic acid in the HFD chow. Furthermore, we discovered that sulfonolipids, which are bacterial-derived lipids^34^, were substantially decreased under HFD feeding. Recent studies have reported that sulfonolipid levels increase with aging in mice^25^, a trend also observed in our dataset, and that sulfonolipids are decreased in the feces of patients with inflammatory bowel disease (IBD)^35^. In addition, representative sulfonolipids such as sulfobacin A and B are known to interact with Toll-like receptor 4. Therefore, the observation that hepatic sulfonolipid abundance changes depending on dietary conditions is biologically intriguing. Although more experiments are required to investigate the mechanisms, the link between the gut microbiome and these lipids raises the possibility that differences in gut microbiota composition between CE2- and HFD-fed mice contribute to these changes. While the molecular mechanisms underlying the absence of lipidomic changes in aged mice fed a HFD remain unclear, aged mice are known to exhibit elevated inflammatory markers across various tissues^36^, which may contribute to reduced hepatic lipid uptake during high-fat feeding. Importantly, although enriched lipid classes and pathways largely recapitulate known biological trends^25, 37^, the increased number of annotated lipid species reduces sparsity in enrichment testing. To test how peak annotation numbers affect the robustness of enrichment analysis, we randomly sub-sampled our datasets and performed the same enrichment analysis (**Fig. 5k**). These results suggest that increasing the number of annotated lipids enhances the sensitivity of lipid ontology analysis, leading to lower adjusted *P* values and thereby supporting more reliable biological interpretation^38^. Detailed mechanistic studies are beyond the scope of this work and will be reported elsewhere.

## Discussion

In this study, we developed the Q1 axis–based deconvolution (Q1Dec) algorithm, implemented direct import and reader of ZT Scan DIA data (WIFF2/WIFF Scan format) into MS-DIAL, and established a workflow for extracting quantitative information from MS2 data. These functionalities have been integrated and released as part of the MS-DIAL 5.6 series. Moreover, we systematically evaluated the effects of Q1Dec and RTDec using both hydrophilic metabolomics and lipidomics datasets. In hydrophilic metabolomics, where annotations are primarily based on spectral similarity scores, application of Q1Dec and RTDec deconvolutions improved multiple similarity metrics including dot product, reverse dot product, enhanced dot product, spectral entropy similarity, and MS-DIAL total score, and resulted in higher annotation coverage and reproducibility compared with DDA-based approaches. In contrast, in lipidomics datasets, we observed a key limitation of deconvolution: commonly shared diagnostic fragment ions, such as *m/z* 184 in positive ion mode, were occasionally removed, leading to an increase in false negatives. Nevertheless, by optimizing Q1 isolation windows and Q1Dec parameters, we could limit this issue to only a few lipids, and found that the mis-exclusion was a predictable phenomenon caused by the proximity of lipids that share the fragment ion. At the same time, deconvolution reduced noise, contaminant ions, and co-eluting signals, thereby decreasing the number of false positives and improving annotation precision (TP/(TP+FP), equivalent to lowering the false discovery rate).

The applicability of Q1Dec filtering critically depends on the degree of chromatographic separation. In workflows where a large number of lipid species co-elute, such as shotgun lipidomics, fragment-to-precursor assignment becomes inherently ambiguous, and Q1Dec is therefore unlikely to provide reliable deconvolution. HILIC-based lipidomics represents an intermediate case. Although lipid classes are separated, multiple molecular species within each class often elute within a narrow retention time window. Under such conditions, the disadvantages of Q1Dec are expected to be more pronounced than in reversed-phase LC; however, subtle retention time differences among molecular species may still provide partial separation. The extent to which Q1Dec can be beneficial or detrimental in HILIC lipidomics remains to be quantitatively evaluated and will require dedicated benchmarking studies. In the present study, we therefore focused on reversed-phase LC–based lipidomics, where the balance between chromatographic separation and DIA multiplexing allowed a quantitative assessment of Q1Dec performance. Based on the results, we concluded that it provides a practical advantage by reducing false positives at the cost of increased false negatives. In practical applications, the higher signal intensity and broader peak coverage achieved with the latest instruments will likely increase the number of detectable DIA signals, including unwanted ions, thereby increasing the value of Q1Dec filtering for reducing spectral complexity and the burden of manual curation. To further support the curation process, we implemented a visualization feature in MS-DIAL that simultaneously displays raw and deconvoluted MS2 spectra. This allows users to readily identify potential mis-exclusion events during data curation. Of note, the main focus of this study was the development and evaluation of deconvolution algorithms, graphical user interface, visualization tools, and quantification tools adapted to ZT Scan DIA, which is independent of the hardware development. In other words, even without further software development, the current limitations may be further mitigated by future hardware improvements that provide higher signal intensity, improved stability, and narrower Q1 isolation windows^39^.

Furthermore, we emphasize that the ability to perform MS2-based quantification using product ions is one of the major advantages of DIA. Lipidomics, where the correspondence between molecules and diagnostic ions can be systematically defined, is particularly well suited to leverage this strength. Although concepts and software for DIA-based MS2 quantification have been proposed previously^40–44^, to our knowledge MS-DIAL is the first platform that enables this functionality at scale using ZT Scan DIA data while directly accessing WIFF files. In this context, the strength of ZT Scan DIA over traditional DIA lies in the possibility to use MS2 spectra deconvoluted by Q1Dec to identify which fragments are free from interferences of other molecules eluting nearby, thereby improving the reliability of quantification. Furthermore, at present, few software environments support direct processing of ZT Scan DIA data, particularly with vendor raw file access and Q1-axis based spectral filtering. Although a command-line based tool Masster (not yet formally published) has reported support for ZT Scan data processing very recently, most widely used metabolomics platforms, including mzmine^45^, XCMS^46^, OpenMS^47^, and asari^48^, do not currently provide direct SCIEX WIFF2/WIFF Scan file import or Q1-axis based deconvolution functionality. In this context, the practical advantage of MS-DIAL lies not only in the implementation of Q1Dec filtering, but also in its graphical user interface, interactive visualization of raw and deconvoluted MS2 spectra, and direct support for large-scale vendor raw data processing. These features facilitate manual data curation and improve usability for routine metabolomics and lipidomics workflows.

In conclusion, our mathematical deconvolution algorithms applied to the ZT Scan DIA data enable the reconstruction of high-purity, compound-specific MS2 spectra that often surpass those obtained from conventional DDA in terms of spectral accuracy. These algorithms were fully integrated into the graphical user interface of MS-DIAL, which now features a dynamic MS2 loader that substantially reduces memory consumption. This allows users to handle ZT Scan DIA files exceeding 20 gigabytes in size (**Supplementary Fig. 7**). This platform improves annotation coverages in both untargeted metabolomics and lipidomics, enables the deconvolution of lipid structural isomers through MS2-based chromatographic features, and supports retrospective analysis because all molecular information is embedded in the raw data.

## Methods

### Dynamic MS2 loader environment

The ability to directly import vendor-specific MS raw data files without file conversion, such as mzML, is one of the notable features of MS-DIAL. WIFF (.wiff), WIFF2 (.wiff2), and WiffScan (.wiff.scan) files are typically generated from recent SCIEX machines, where MS spectral data are stored in a WiffScan file and spectral retrieval can be performed from WIFF or WIFF2 files via software development kits. Both the WIFF and WIFF2 data on direct imports are supported by MS-DIAL. The dynamic MS2 loader functions were developed for the WIFF2 format file, where the ZT Scan DIA metadata information was fully supported.

For data processing that included the processes of peak picking, MS2 peak purification, peak annotation, and peak alignment, the previous MS-DIAL program stored all MS1 and MS2 data in random-access memory because the data size of MS2 was similar to or typically lower than that of MS1. However, the situation changed in ZT Scan DIA, where the MS2 data size (10–20 GB) became 100–200 times larger than that of the MS1 (10–200 MB). This is also true for other high-end machines, such as the Orbitrap Astral of Thermo Fisher Scientific. Furthermore, the current MS-DIAL program only requires MS1 data for the peak picking and peak alignment processes, and not all MS2 data are used for spectral deconvolution and annotation. These facts motivated us to develop a dynamic MS2 loader system that calls MS2 data on demand. Thus, the source codes for MS2 data retrieval were completely rewritten in this study to execute on-demand MS2 extraction, whereas the spectra were stored in the cache once the data were retrieved. This allowed one to save memory space and reduce the required time for ZT Scan DIA data analysis. Although this environment is currently available in SCIEX WIFF2 format, the dynamic MS2 loader will become available for all MS vendor data in the future. Importantly, the system is available not only for ZT Scan DIA, but also for other DIA methods, such as SWATH-DIA, in addition to the DDA method.

### Q1Dec: Q1-axis information-based filtering

After peak picking based on retention time and MS1 *m/z* dimensions (RT–*m/z* features), Q1Dec filtering was applied to each RT–*m/z* feature. The MS2 spectral data corresponding to the closest precursor *m/z* value (Q1 bin; typically acquired with a step size of ∼1.0–1.5 Da) to the target RT–*m/z* feature were retrieved. In addition, MS2 spectra from neighboring Q1 bins within a user-defined tolerance around the central Q1 bin were also collected.

For each product ion, its intensity trace across the selected Q1 bins was evaluated, and the ion was retained if its local maximum was located within the specified Q1-bin tolerance; otherwise, it was excluded as a contaminant. The Q1-bin tolerance is a user-configurable parameter (e.g., ±1, ±2, or ±3 bins), allowing flexible control of filtering stringency depending on the analytical context.

### RTDec: RT-axis information-based MS2 deconvolution

The MS2 spectrum constructed using the Q1Dec function was used as the starting spectrum for the RTDec process. Furthermore, users could set two thresholds before starting the deconvolution process: the MS2 absolute peak height cutoff (default zero) and the relative MS2 peak height cutoff (default zero). These were set to 100 and 0.0%, respectively, for hydrophilic metabolomics data, whereas the thresholds of the MS2 absolute peak height were determined using the MS2 peak heights of barcode (noise) ions. This resulted in threshold values of 50, 100, and 160 for DDA, SWATH-DIA, and ZT Scan DIA, respectively, for untargeted lipidomics data. The MS2 relative abundance cut-off was set at 0.0%. These thresholds are useful for reducing the processing time for deconvolution via ZT Scan DIA and clean-up of noise ions whose abundance typically increases when a short accumulation time (higher scanning speed for example) is used, as performed in ZT Scan DIA. Filtering was applied to the Q1Dec deconvoluted MS2 spectrum.

By using product ion values in the starter MS2 spectrum with a user-defined binning value (0.05 Da in this study), the MS2 chromatograms are extracted from the Q1 bin matched with the precursor *m/z* of interest. The RT range was set as the peak maximum plus or minus twice the peak width. The computational workflow, including MS2 peak quality estimation, MS2 model chromatogram construction, and the least squares method to provide the deconvoluted MS2 chromatograms, were the same as those of the MS2Dec algorithm described in a previous study.

### MS2 feature extractions based on MS-DIAL lipid in silico MS2 templates or MS2 correlation matrix

DIA techniques provide an environment for the use of MS2 chromatogram data for peak annotation and quantification of co-eluted structural isomers. In this study, we developed several functions to utilize MS2 spectra for MS-DIAL (**Supplementary Fig. 5 and Supplementary Fig. 8**). In lipidomics, the rt-*m/z* peak features satisfying the following three prerequisites are expanded into the MS2 dimension for further feature extraction. First, the peak must be detected for all samples. The gap-filled peaks in the peak alignment process are beyond the scope of this study. Second, the lipids of the second or later candidates must be annotated at the molecular species level, where up to three co-eluted isomers (candidates) are considered in this study. Finally, the MS2 data should be available for all samples, although this requisite is always true for DIA. The function to provide the peak features in the rt-*m/z*-MS2 dimensions was applied for the peak alignment results. Furthermore, a function to determine the unique MS2 *m/z* value (unique mass) for isomer separation and quantification was developed. For this function, the *m/z* value, which is not used for annotation in other isomer candidates, is automatically determined, while users define their own unique mass in the MS-DIAL graphical user interface. If the annotation is performed in experimental MS2 spectral libraries such as MassBank, MoNA, and NIST, the product ion that is contained as a minor peak with less than 1% relative intensity in the other candidates is also considered a candidate for a unique mass. Only one unique mass value is currently used in the MS-DIAL program.

In contrast to the above environments, where annotations are essential and unique mass determination is based on the reference MS2 spectra, MS-DIAL offers a different method (MS2 correlation matrix) to extract unique metabolite features that contain both known and unknown peaks (**Supplementary Fig.7**). Dot-product similarity scores were calculated for all sample pairs in an aligned peak, where users could set their own thresholds to define the number of unique components. In this function, the unique mass is defined from the experimental MS2 spectra, where the intensity values are summed within a 0.5-Da bin, and the product ion peak with more than 70% relative peak intensity is recognized as a unique mass candidate as the default setting. If unique mass values were shared among some components, the components were treated as a component group. This was similar to the protein grouping algorithm performed in the conventional shotgun proteomics data analysis pipeline. The feature and its application will be detailed elsewhere.

### Benchmark data acquisitions to compare DDA, SWATH-DIA, and ZT Scan DIA technologies

Human plasma samples (NIST SRM 1950) were purchased from the National Institute of Standards and Technology (NIST). The Bligh and Dyer technique was used for lipid extraction. Briefly, 10 μL of blood sample was mixed with 1,000 μL of an ice-cold MeOH/CHCl_3_/H_2_O (10:4:4, v/v/v) solvent. Solvents were vortexed for 10 s, centrifuged at 15,000 × g for 10 min at 4°C, and the lipid layer of the supernatant was transferred to a clean tube. Supernatant was dried using a speedvac and resuspended into a final concentration of 1 µL plasma per 10 µL of injection solvent. Hydrophilic metabolite extraction was performed via methanol protein precipitation with four volumes of ice-cold methanol. The solvent was vortexed for 10 s, and the solution was centrifuged at 15,000 × g for 10 min. Supernatant was dried using a speedvac and resuspended in water to a final concentration of 1 µL extract equivalent to 0.1 µL plasma. Three technical replicates were used for each experiment.

LC-MS/MS analysis was performed using an Agilent Integrated System for LC comprising a G7167B autosampler, G7120A binary pump, G7116B column management system, and ZenoTOF 8600 system for MS (SCIEX, Toronto, Canada). For hydrophilic metabolomics, the metabolite separation was performed using the Kinetex F5 column (150 × 2.1 mm, 2.6 μm, Phenomenex Inc., CA, US) and the following mobile phases: (A) 0.1% formic acid in H_2_O and (B) 0.1% formic acid in acetonitrile (ACN). The column was maintained at 30°C and a flow rate of 0.2 mL/min. A sample volume of 3 μL was used for the injection. The separation was performed using the following gradient: 0 min 0.0% (B), 2.0 min 0.0% (B), 14.0 min 95% (B), 16.0 min 95% (B), 16.1 min 0% (B), and 23.0 min 0% (B). The common MS conditions were as follows: ion source gas 1, 40 psi; ion source gas 2, 60 psi; curtain gas, 40 psi; charged aerosol detector (CAD) gas, 7; source temperature, 500°C; spray voltage, 4000 V; *m/z* range of time-of-flight (TOF)-MS, 70‒1000 Da; *m/z* range of TOF MS/MS, 50‒1000 Da; QJet declustering potential, 20 V; collision energy, 30 eV; Q1 resolution, unit. The following conditions were used for ZT Scan DIA: Q1 width, 7.2 Da; Q1 speed, 1390 Da/s; Q1 precursor range, 60–1000; MS1 accumulation time, 50 ms; and cycle time, 0.784 s. For SWATH DIA, the precursor isolation window was set to 50 with a 1-Da overlap, and the Q1 precursor range was set to 80–950 to match the cycle time, as used in the ZT Scan DIA. The MS1 accumulation time was set to 100 ms, and the cycle time was 0.764 s. For DDA, the conditions of the top 50 precursors per survey were used. The MS1 accumulation time was set to 100 ms, and the cycle time was 0.668 s.

For lipidomics, the lipid separation was carried out with the column of Luna C18 (150 × 2.0 mm, 2.1 μm, Phenomenex Inc., CA, US) and the mobile phases of (A) ACN/isopropyl alcohol (IPA)/H_2_O (3:2:5, v/v/v) and (B) ACN/IPA (1:9, v/v) where both contained 10 mM ammonium acetate. The column was maintained at 55°C and a flow rate of 0.4 mL/min. A sample volume of 5 μL was used for the injection. The separation was performed using the following gradient: 0 min 10.0% (B), 2.7 min 45.0% (B), 2.8 min 53% (B), 9 min 65% (B), 9.1 min 89% (B), 11.0 min 92% (B), 11.1 min 100% (B), 11.9 min 100% (B), 12.0 min 10% (B), and 15.0 min 10% (B). The MS conditions for DDA and SWATH-DIA were as follows: ion source gas 1, 40 psi; ion source gas 2, 60 psi; curtain gas, 40 psi; CAD gas, 7; source temperature, 350°C; spray voltage, 5500 V; *m/z* range of TOF-MS, 70‒1000 Da; *m/z* range of TOF MS/MS, 50‒1000 Da; QJet declustering potential, 20 V; collision energy, 30 eV; Q1 resolution, unit; MS1 accumulation time, 100 ms; MS2 accumulation time, 5 ms.

The MS conditions for ZT Scan DIA were as follows: ion source gas 1, 50 psi; ion source gas 2, 70 psi; curtain gas, 40 psi; CAD gas, 7; source temperature, 250°C; spray voltage, 5500 V; QJet declustering potential, 20 V; collision energy, 42 eV; collision energy spread 15 eV; time bins to sum (MS1), 6; time bins to sum (MS2), 6; MS1 accumulation time, 50 ms; *m/z* range of TOF-MS, 200–1250; *m/z* range of TOF-MS/MS, 100–1250; Q1 resolution, unit; zeno pulsing, enabled; zeno threshold, 2000000 cps; collision energy, 37 eV; collision energy spread, 0 eV; Q1 width, 6.8 Da; Q1 scan speed, 1015 Da/s; total MS/MS scan time, 1040 ms.

### MS-DIAL parameters

MS-DIAL was built as version “5.6” where dynamic MS2 loading is available when a WIFF2 file format is imported. Importantly, we developed the “rawdatahander.dll” package independently to handle raw MS data. The package uses a vendor-specific application programming interface to convert the profile spectrum into a centroid spectrum. This means that the current rawdatahandler.dll always parses the spectrum as a centroid. Given that the MS-DIAL program uses the rawdatahander package, MS-DIAL always retrieves centroid mass spectra as long as users import the vendor’s raw data. Therefore, “centroid” was selected as the ms1 and ms2 data type in the MS-DIAL parameter settings. In the analysis file settings window, acquisition type was set to “DDA,” “SWATH,” or “ZTScan” according to the data sets. The following parameters were used: max number of isotope recognitions, 5; number of threads, 6; minimum peak amplitude, 500; MS/MS absolute abundance cut-off, 50, 100, and 160 for DDA, SWATH-DIA, and ZT Scan DIA for untargeted lipidomics, respectively; 100 for hydrophilic metabolomics data; MS/MS relative abundance cut-off, 0; and exclusion after precursor *m/z*, 0.5. The Q1-bin tolerance, referred to as the “retrieved Q1-bin region” in MS-DIAL, was evaluated at ±0, ±1, ±2, and ±3 bins. These tolerances correspond to parameter settings of ±1, ±2, ±3, and ±4 bin regions, respectively, in the MS-DIAL software. For hydrophilic metabolomics, the following annotation parameters were used: simple dot product, 600; weighted dot product, 600; reverse dot product, 800; minimum matched peak number, three; and minimum matched peak percentage, 0%. In lipidomics, the annotation parameters were set as follows: dot product, 150; weighted dot product, 150; reverse dot product, 300 (default in MS-DIAL); minimum matched peak number, 1; and minimum matched peak percentage, 0%. Default parameter values were used for the other parameters. All data were analyzed using a Windows OS personal computer with the following system: processor, AMD EPYC 7452 32-Core processor with 2.35 GHz; installed RAM, 256 GB, graphics card, NVIDIA Quadro P4000 (8 GB), and storage, 13.2 TB.

### Evaluation of annotation rates and spectral similarity values when compared with those of the experimental MS2 spectral libraries

Hydrophilic metabolomics data from NIST SRM 1950 human plasma (three technical replicates) were analyzed using three acquisition methods: data-dependent acquisition (DDA), SWATH-DIA, and ZT Scan DIA. An in-house MS2 spectral library containing 1,492,414 spectra from 259,277 compounds in positive ion mode was used. This library was compiled from multiple sources, including RIKEN Plasma, KI-GIAR ZIC-HILIC, BMDMS-NP, CASMI 2012, CASMI 2016, EMBL-MCF, EMBL-MCF 2.0 HRMS, Fiehn HILIC, GNPS, HCD Natural Product Library, MassBank, MetaboBase, METLIN, NIST14, NIST20, Pathogen Box, PEP-NP, QiaoLab PGN, ReSpect, and Vaniya–Fiehn Natural Products libraries. The options “use retention time for scoring” and “use retention time for filtering” were disabled. All other parameter settings were identical to those described above for hydrophilic metabolomics analysis. For ZT Scan DIA data, four deconvolution conditions were evaluated: Q1Dec + RTDec, Q1Dec only, RTDec only, and no deconvolution. In addition, Q1-bin tolerances of ±0, ±1, ±2, and ±3 bins were tested. SWATH-DIA data were analyzed with and without RTDec. ZT Scan DIA and SWATH-DIA are denoted as ZT and SW, respectively. Spectral similarity was evaluated using weighted dot product, reverse dot product, enhanced dot product, and spectral entropy similarity scores, calculated with default hyperparameters as described in the original publications^20, 21^. The MS-DIAL total score was also calculated as a weighted sum of the dot product, reverse dot product, and matched peak percentage, with weighting factors of 3, 2, and 1, respectively. The matched peak percentage was defined as the ratio of the number of product ions matched to the reference spectrum to the total number of product ions in the reference spectrum.

### Investigating the effect of product ion abundance in the Q1Dec filtering

The outcomes of Q1Dec filtering for observed product ions can be classified into four categories: (1) contaminant product ions that should be removed are correctly removed (true negative: TN), (2) true product ions that should be retained are correctly retained (true positive: TP), (3) contaminant product ions that should be removed are incorrectly retained (false positive: FP), and (4) true product ions that should be retained are incorrectly removed (false negative: FN). A practical reference set for “ions that should be removed” and “ions that should be retained” was constructed using the hydrophilic metabolomics dataset from human plasma. Peaks annotated after Q1Dec processing with reverse dot product ≥ 0.9 were extracted from the library search results, and the experimental MS2 spectra were compared with the corresponding reference MS2 spectra using an *m/z* binning value of 0.05. Product ions present in the reference spectrum were defined as “ions that should be retained,” whereas ions absent from the reference spectrum but present in the raw spectrum before deconvolution were defined as “ions that should be removed.”. Based on this definition, the product ion intensity distributions for TP, TN, FP, and FN were investigated (**Supplementary Fig. 2**).

### Evaluation of annotation rates in lipidomics

Lipid annotation was performed using a rule-based decision-tree algorithm optimized for each lipid subclass. The MS-DIAL version 5.6 series was used, in which a total of 160 lipid subclasses were managed. False hits were manually excluded based on the RT behavior of lipids in reversed-phase chromatography, according to a previous study^49^. The lipids annotated as the adduct forms of [M-H_2_O+H]^+^, [M+H]^+^, or [M+NH_4_]^+^ for the positive ion mode and [M-H]^−^ and [M+CH_3_COO]**^−^**for the negative ion mode were used to compare the annotation coverage among the analytical techniques.

### Experimental design to showcase three-dimensional feature extractions in ZT Scan DIA

The animal experiments were conducted in accordance with the ethical protocol approved by the Tokyo University of Agriculture and Technology (R5-50). C57BL/6J male mice were purchased from SLC (Shizuoka, Japan). Mice at 11 weeks or two years of age were used for the experiments and sacrificed for tissue collection. All mice were initially maintained on CE-2 chow (CLEA Japan, Tokyo, Japan), and a subset of animals was switched to an HFD (D12492; Research Diets, New Jersey, USA) four weeks before sacrifice. The livers were harvested, and tissues were immediately frozen after dissection and stored at −80°C until lipid extraction. NIST SRM 1950 plasma was also analyzed as a benchmark.

Lipids were extracted on ice using a biphasic solvent system comprising ice-cold methanol (MeOH; FUJIFILM Wako Pure Chemical, Osaka, Japan), MTBE (Sigma-Aldrich, Tokyo, Japan), and ultrapure water (UP-0090a-0U1; Organo, Tokyo, Japan). Unless otherwise stated, all solvents were kept on ice throughout the procedure. An internal standard (IS) mixture containing EquiSPLASH LIPIDOMIX (Avanti Polar Lipids, USA) and FA 16:0-d₃/FA 18:0-d₃ (SRL, Canada) was used for normalization. The detailed composition of the IS mixture is provided in **Supplementary Data 1** and **Supplementary Data 2**. For plasma samples from NIST SRM 1950 (20 µL, n = 3), 225 µL of MeOH containing internal standards was added. After vortexing for 10 s, 750 µL of MTBE was added. The mixture was vortexed again, sonicated for 10 min, shaken at 4°C for 30 min, and centrifuged at 16,000 × g for 3 min at 4°C. The upper organic phase (800 µL) was collected, mixed with 154 µL of water, vortexed, and centrifuged again. A total of 560 µL of the upper layer was collected, divided equally into four tubes, dried under vacuum, and stored at −80 °C. Frozen mouse liver tissues were homogenized at 2500 rpm for 15 s using a multibead shocker equipped with a metal cone (YASUI KIKAI, Japan). One milliliter of MeOH was added to the homogenate. An aliquot corresponding to 5.0 mg of tissue was transferred and diluted to a final volume of 225 µL with MeOH containing internal standards. Lipid extraction was performed as previously described. All dried lipid extracts were reconstituted in 40 µL of MeOH, vortexed for 30 s, centrifuged at 16,000 × g for 3 min at 4°C, and transferred to LC-MS vials fitted with micro-inserts (Agilent Technologies, USA).

The LC-MS/MS analysis was performed using a Nexera XR UHPLC system (SIL-40C XR autosampler, LC-40D XR binary pump, CTO-40C column oven, and SCL-40 system controller; Shimadzu, Kyoto, Japan) coupled to a ZenoTOF 8600 mass spectrometer (SCIEX, Toronto, Canada). Lipids were separated on an Acquity UPLC Peptide BEH C18 column (50 × 2.1 mm, 1.7 μm; Waters, Milford, MA, USA) maintained at 45°C with a flow rate of 0.3 mL/min. The mobile phases consisted of (A) ACN/methanol/water (1:1:3, v/v/v) with 5 mM ammonium acetate and 10 nM ethylenediaminetetraacetic acid and (B) ACN/isopropanol (1:9, v/v) with the same additives (FUJIFILM Wako Pure Chemical, Osaka, Japan). A sample volume of 2 μL was used for the injection. The separation was conducted under the following gradient: 0 min 0.1% (B), 1 min 0.1% (B), 5 min 40% (B), 7.5 min 64% (B), 12 min 71% (B), 12.5 min 82.5% (B), 19 min 85% (B), 19.1 min 99.9% (B), 20.5 min 99.9% (B), 20.6 min 0.1% (B), and 25 min 0.1% (B). The temperature of each sample was maintained at 4°C.

The MS conditions were as follows: ion source gas 1, 50 psi; ion source gas 2, 70 psi; curtain gas, 40 psi; CAD gas, 7; source temperature, 250 °C; spray voltage, 5500 V/−4500 V; QJet declustering potential, ±20 V; collision energy, 42 eV; collision energy spread 15 eV; time bins to sum (MS1), 4; time bins to sum (MS2), 6; MS1 accumulation time, 100 ms; *m/z* range of TOF-MS, 200–1250; *m/z* range of TOF-MS/MS, 100–1250; Q1 resolution, unit; zeno pulsing, enabled; zeno threshold, 2000000 cps. For DDA, the following conditions were used: MS2 accumulation time, 25 ms; maximum candidate ion, 35; Intensity threshold exceeds, 10 counts/s; Exclude former candidate ions, disabled; Inclusion/Exclusion list, disabled. For SWATH-DIA, MS2 accumulation time, 17 ms; isolation window width, 20 Da. For ZT-scan DIA, the following conditions were used: collision energy, ±37 eV; collision energy spread, 0 eV; time bins to sum (MS1), 6; Q1 width, 6.8 Da; Q1 scan speed, 1015 Da/Sec; total MSMS scan time, 1040 ms.

Data analysis was performed using MS-DIAL 5.6 with the same parameter set as described in the previous section. Data curation, reduction, and normalization were performed as previously described. Briefly, the annotation results were manually curated, and lipids that did not satisfy the RT behaviors in reverse-phase LC were excluded. The positive and negative ion mode data were integrated, where only one adduct form was retained; for example, the acetate adduct form was used for PC, whereas the protonated and sodium adduct forms of PC were excluded. Although the normalized MS1 quantitative values reflect the concentrations of lipid molecules in pmol per mg of tissue, the MS2 quantitative values cannot provide such semi-quantitative values without detailed validation experiments. In this study, normalization of MS2 quantitative values was performed through the following process. First, each MS2 peak height was divided by the MS1 peak height of the corresponding internal standard (IS) assigned to the lipid class at the precursor ion level. The correspondence between lipid annotations and internal standards is provided in **Supplementary Data 1/2**. For example, if the MS1 peak was annotated as PC A_B and two MS2-level annotations (PC A_B and PE C_D) were assigned to that precursor, both MS2 peak heights were divided by the MS1 peak height of the phosphatidylcholine internal standard included in EquiSPLASH, i.e., PC 15:0_18:1(d7). In this way, sample-to-sample analytical variation observed at the MS1 level was taken into account. Although this may not represent the ideal normalization strategy, it was considered a practical approach for incorporating analytical variation into MS2-based quantification. In addition, because the absolute intensity scales of MS1 and MS2 data are often substantially different, and because internal standard peak intensities are typically much higher than those of endogenous metabolite peaks, the normalized values after division are often represented as very small decimal values. This makes it difficult for users to intuitively assess the magnitude of the original MS2 peak intensities. To preserve interpretability, the normalized values were further multiplied by the average MS2 peak height of the corresponding annotated lipid across all samples. This rescaling step allows the normalized values to retain the approximate magnitude of the original MS2 intensities while still reflecting normalization by the internal standard. This normalization process was saved with the excel functions in **Supplementary Data 1/2**.

The total number of lipid species reported in this study was calculated by merging lipid annotations obtained at the MS1 and MS2 levels and removing duplicate lipid names. The subset of lipid species supported by MS2-based quantitative values is smaller because MS-DIAL currently expands MS1 features into the MS2 dimension only when the corresponding peak is detected in all samples. The MS2 quantitative values for alignment peak features where gap-filled MS1 features exist are not generated due to the current limitation of MS-DIAL data structure.

### Evaluation of spectral deconvolution processes in lipidomics data

To evaluate the efficiency of Q1Dec and RTDec proceess in lipidomics, the benchimark data were acquired. Lipid extraction was performed according to the Bligh and Dyer technique with slight modification^50^. The extraction solvent, a mixture of methanol (MeOH; FUJIFILM Wako Pure Chemical, Osaka, Japan), chloroform (CHCl_3_; FUJIFILM Wako Pure Chemical, Osaka, Japan), and ultrapure water (H_2_O; UP-0090a-0U1; Organo, Tokyo, Japan) in a volume ratio of 10:4:4 (*v/v/v*), was pre-mixed and sonicated for 5 min before use. All solvents were kept on ice throughout the procedure. 1000 µL of the extraction solvent was added to each 20 µL plasma sample (NIST SRM 1950) in a 2 mL tube (Eppendorf, Enfield, USA). The mixture was vortexed for 1 min and subjected to ultrasonic extraction for 10 min (Bioruptor II, Sonic Bio, Kanagawa, Japan). After centrifugation at 16,000 × g for 5 min at 4°C, 900 µL of the supernatant was transferred to a new 2 mL tube. To this supernatant, 300 µL of CHCl_3_ and 252 µL of H_2_O were added. The mixture was mixed without vigorous vortexing and centrifuged at 16,000 × g for 3 min at 4°C. The lower organic phase was collected into new 1.5 mL tubes, evaporated, and resuspended in 100 µL of MeOH containing EquiSPLASH LIPIDOMIX (Avanti Polar Lipids, USA) and FA 16:0-d₃ / FA 18:0-d₃ (SRL, Canada) as internal standards for normalization. After vortexing for 30s and centrifuged at 16,000 × g for 3 min at 4°C, the supernatant was transferred to LC-MS vials with micro-glass-inserts (Agilent Technologies, Santa Clara, USA).

The LC-MS/MS analysis was performed with an ExionLC system coupled to a ZenoTOF 7600+ mass spectrometer (SCIEX, Toronto, Canada). Lipids were separated on an Imtakt UK-C18 MF column (50 × 2 mm; 3 µm) (Imtakt, Kyoto, Japan) maintained at 45 °C with a flow rate of 0.3 mL/min. The mobile phases consisted of (A) acetonitrile/methanol/water (1:1:3, *v/v/v*) with 5 mM ammonium acetate and 10 nM ethylenediaminetetraacetic acid (EDTA), and (B) water/isopropanol (1:9, *v/v*) with the same additives (all reagents from FUJIFILM Wako Pure Chemical, Osaka, Japan). The injection volumes were 0.5 μL for positive ion mode and 1 μL for negative ion mode. The separation was conducted under the following gradient: 0 min 0.5% (B), 1 min 0.5% (B), 5 min 40% (B), 7.5 min 64% (B), 12 min 71% (B), 12.5 min 82.5% (B), 19 min 85% (B), 20 min 99.0% (B), 22 min 99.0% (B), 22.1 min 0.5% (B), and 27 min 0.5% (B). The temperature of the samples was maintained at 4 °C. The common MS conditions for positive/negative ion modes were as follows: ion source gas 1, 40/50 psi; ion source gas 2, 80/50 psi; curtain gas, 30/35 psi; CAD gas, 7; source temperature, 250 °C/300 °C; spray voltage, 5500 V/−4500 V; *m/z* range of TOF-MS, 75–1250; declustering potential, ±80 V; collision energy, 40 eV/−42 eV; collision energy spread 0 eV; Q1 resolution, unit; zeno pulsing, enabled; zeno threshold, 2000000 cps. For DDA, the following conditions were used: *m/z* range of TOF-MS/MS, 75–1250; total scan time, 788 ms; MS1 accumulation time, 200 ms; MS2 accumulation time, 50 ms; time bins to sum (MS1/MS2), 4; maximum candidate ion, 10; Intensity threshold exceeds, 10 counts/s; Exclude former candidate ions, disabled; Inclusion/Exclusion list, disabled. For ZT Scan DIA, the following conditions were used: cycle time, 1300 ms; MS1 accumulation time, 100 ms; total MSMS scan time, 1140 ms; processed time bins to sum (MS1/MS2), 6. For the analysis of NIST SRM 1950 plasma samples, five different Q1 widths were applied for each ion mode. The *m/z* range of TOF-MS/MS, precursor *m/z* range, and Q1 scan speed were adjusted according to the Q1 width, as detailed in **Supplementary Data 3**.

The raw data were processed using MS-DIAL ver 5.6. The following parameters were used: max number of isotope recognitions, 5; the number of threads, 12; the minimum peak amplitude, 100. The default parameter values were used for the others. For the DDA data of plasma samples (n=3), the peaks were aligned and subjected to manual curation. The curation process involved a visual inspection of both retention time (RT) regularity and MS2 spectral quality. Specifically, we eliminated false-positive annotations where diagnostic ions were incorrectly assigned to stochastic, background noise rather than distinct fragment signals. This curated DDA dataset was defined as the reference annotation set for subsequent evaluations of the ZT Scan DIA data processing methods.

### Data visualization and statistical analyses

All data visualization, significance testing, and multivariate analyses were conducted using the R programming language. For principal component analysis (PCA) and orthogonal partial least squares discriminant analysis (OPLS-DA), variables were log-transformed and standardized by auto-scaling. In OPLS-DA, only the CE2 diet conditions in liver lipidomics dataset was used to compare young (2-month-old) and aged (24-month-old) mice. Variable importance in projection (VIP) values were calculated, and the top 40 variables were visualized as a heatmap.

For liver lipidomics data, statistical comparisons were performed for the following groups: CE2/M2 vs. CE2/M24, HFD/M2 vs. HFD/M24, CE2/M2 vs. HFD/M2, and CE2/M24 vs. HFD/M24. Significance testing was conducted using Student’s t-test (two-sided, equal variance assumed; N = 4), with P values adjusted by the Benjamini–Hochberg method. Lipids with an adjusted P value ≤ 0.05 and a fold change ≥ 1.5 or ≤ 0.5 relative to control were defined as differentially expressed lipids (DELs).

Lipid molecule enrichment analysis was performed using the clusterProfiler package. First, a lipid metadata library (**Supplementary Data 4**) was generated from lipid molecule names in the liver lipidomics dataset. Lipid class names and acyl chain labels were stored in a long-format data object, which was provided as input to clusterProfiler with the DELs information to evaluate enrichment of lipid classes and acyl chains using over-representation analysis (ORA). Lipids belonging to enriched terms were selected for heatmap visualization according to the following criteria: (1) select one lipid molecule from each enriched term, prioritizing those with the highest mean abundance across all samples; (2) repeat selection from terms in order of *P* value from the enrichment analysis until a total of 10 lipids were chosen, prioritizing those with higher abundance in subsequent selections. The relationship between the number of annotated lipids and statistical significance in enrichment analysis was investigated according to the previously reported method^38^. The MS1- and MS2-based metabolite tables (Supplementary Data 2) were merged into a unified dataset. Lipid variables with zero values in at least one sample were excluded from the analysis. Differential analysis between HFD/M2 and CE2/M2 conditions was performed using Student’s t-test, and *P* values were adjusted using the Benjamini–Hochberg method to control the false discovery rate (FDR). Significantly increased and decreased lipid species were then subjected separately to ORA to extract enriched terms (adjusted *P* values were used for interpretation). To evaluate the effect of lipid coverage on enrichment analysis, the total number of lipid variables included in the analysis was subsampled to 100, 500, 1,000, 1,500, and 1,663 (maximum). For each subsampling condition, enrichment analysis was repeated 50 times, and the median *P* value for each term was used as the representative value. Terms containing 10 or fewer lipid molecules were excluded from visualization to avoid instability in enrichment statistics (e.g., the ADGGA subclass, which contained only four lipid species, was excluded).

## Supplementary Figures

**Supplementary Figure 1. Overview of tandem mass spectrometry (MS/MS) data acquisition strategies in liquid chromatography (LC)-MS/MS.** In conventional LC-MS/MS, full precursor ion scanning (MS1) and tandem MS acquisition (MS2) are performed sequentially. In data-dependent acquisition (DDA), precursor ions are isolated with narrow windows (typically <1 Da) to generate high-purity MS2 spectra. In contrast, data-independent acquisition (DIA), such as SWATH-DIA implemented on SCIEX instruments, utilizes wider isolation windows ranging from 5–50 Da. SWATH-DIA applies a block-wise acquisition strategy, pausing briefly (typically <1 ms) between isolation windows. ZT Scan DIA differs fundamentally by employing a continuous sliding precursor isolation strategy, scanning across the *m/z* range with 5–20-Da windows and eliminating the need for pause times between window transitions.

**Supplementary Figure 2. Effect of product ion abundance on Q1Dec filtering performance in hydrophilic metabolomics data.** The relationship between product ion abundance and Q1Dec filtering outcomes was visualized using violin plots. Solid lines indicate median values, and half-solid/half-dashed lines indicate mean values. Four categories were evaluated: (1) contaminant product ions that should be removed and were correctly removed (labeled as “TN: true negative”), (2) product ions that should be retained and were correctly retained (labeled as “TP: true positive”), (3) contaminant product ions that should be removed but were incorrectly retained (labeled as “FP: false positive”), and (4) product ions that should be retained but were incorrectly removed (labeled as “FN: false negative”). The intensity distribution of all product ions (“All”) is also shown.

**Supplementary Figure 3. Showcase of Q1Dec filtering in an untargeted lipidomics data.** (a) The tandem mass spectra before and after Q1Dec filtering method for the peaks eluted at *m/z* 760.58 and 8.25 min region. Extracted ion chromatograms (EIC) for *m/z* 760.5841, 768.5594, and 770.6047 are shown. An ion heatmap of *m/z* 184.072 in the MS2 spectrum of protonated phosphatidylcholine (PC) 16:0_18:1 (*m/z* 760.5841) is presented along retention time (RT) and quadrupole (Q1) axes. MS2 spectra corresponding to the three peaks were extracted from both data-dependent acquisition (DDA) and ZT Scan DIA data, with Q1 axis-based spectral processing (Q1Dec) applied to the latter.

**Supplementary Figure 4. Mis-exclusion rates of diagnostic fragment ions for lipid annotations.** Recalls of diagnostic fragment ions in ZT Scan DIA data processed with under different conditions (no deconvolution, None; only RTDec used, RTDec; only Q1Dec used, Q1Dec; both Q1Dec and RTDec used, Q1/RTDec) were described.

**Supplementary Figure 5. MS-DIAL function for lipid name–centric deconvolution.** This illustration shows the conceptual workflow of lipid name–centric deconvolution in analyzing six biological samples. In samples 2, 3, and 4, two molecular species–level annotation candidates are assumed to coexist within a single peak feature defined by the same precursor *m/z* and retention time. Across the six samples, three lipid candidates (PC 16:1_22:5, PC 18:2_20:4, and PC 18:1_20:5) are included in the same alignment peak feature. By extracting diagnostic product ions specific to each lipid molecule and using them for quantification, structural isomers that cannot be distinguished at the MS1 feature level can be resolved as independent MS2 features.

**Supplementary Figure 6. Orthogonal partial least squares discriminant analysis (OPLS-DA) and variable importance in projection (VIP) analysis for age-related lipidomic differences.** OPLS-DA score plots were generated using auto-scaled lipid variables to capture age-related differences in the mouse liver lipidome. In addition, a heatmap of the representative lipid molecules ranked by variable importance in projection (VIP) scores is shown.

**Supplementary Figure 7. Summary of dynamic MS2 data loading in the new MS-DIAL program (version 5.6 series).** In previous versions of MS-DIAL, all MS1 and MS2 data from a single raw file were loaded entirely into random access memory (RAM) before data processing was initiated. However, for datasets such as ZT Scan DIA, where extremely large amounts of MS2 data are contained and only a limited portion of the MS2 space contains analytically meaningful information, this approach is not computationally efficient. In such cases, peak detection in the MS1 dimension can be performed immediately after loading only the MS1 data into RAM, without requiring full MS2 loading. Similarly, during deconvolution processes such as Q1Dec and RTDec, extracting only the required MS2 regions on demand improves computational efficiency and substantially reduces RAM usage. This functionality required major structural changes to the original MS-DIAL source code and involved substantial development effort. Although the dynamic MS2 reading platform is still under active development, it provides the foundation for practical and scalable processing of large DIA datasets.

**Supplementary Figure 8. Summary of spectrum-centric deconvolution strategy.** This illustration assumes the analysis of six biological samples. MS2 spectra with high similarity and substantial shared fragment ions are grouped and defined as consensus spectra (ConSpec). In this example, for a single alignment peak feature defined by the same precursor *m/z* and retention time, the MS2 spectra from the six samples are classified into three consensus spectra, A, B, and C, indicating that three distinct co-eluting components with different MS2 patterns are present. The definition of ConSpec is based on spectral similarity across samples and can be adjusted by parameter settings in MS-DIAL. In this illustration, product ions were binned with a width of 0.5 Da, and spectra with a dot product score greater than 0.8 were grouped into the same ConSpec. To export quantitative values for components A, B, and C, product ions used for quantification must be selected from each consensus spectrum. Two strategies are currently supported. The default strategy requires that the product ion be detected in all samples belonging to the corresponding ConSpec. Alternatively, if the ConSpec can be annotated using an authentic standard spectral library, product ions with relative abundance greater than 50% in the reference spectrum can be selected as quantitative ions (optional setting). However, because ConSpecs cannot always be annotated by standard spectral libraries, quantitative product ions must also be defined for unannotated components. In such cases, it is practical to prioritize high-intensity product ions that are unique to one component and absent from the others. Therefore, the current MS-DIAL implementation uses the strategy illustrated in the lower panel. First, each spectrum of ConSpec A, B, and C is binned at 0.5 Da intervals. For each *m/z* bin, the presence or absence of signals across the consensus spectra is evaluated. In this example, eight *m/z* bins are shown. For example, *m/z* bin 1 is detected only in ConSpec C, whereas *m/z* bin 2 is shared among all three ConSpecs (A, B, and C). A product ion is defined as “detected” when its relative abundance exceeds 70% within the same MS2 spectrum (default setting). In this illustration, ConSpec C contains unique product ions, whereas A and B do not have any unique ions. In such cases, the current MS-DIAL strategy treats A and B as a single quantitative group and exports them together as a combined feature (A + B).

## Data availability

All raw MS data are available in the MB-POST repository (https://repository.massbank.jp/) under index number MPST000090, MPST000175, MPST000176, and MPST000179. The processed table data used to create the figures were available at Zenodo (https://doi.org/10.5281/zenodo.20763288), and the original peak lists and alignment tables were deposited in MB-POST together with the raw data. Lipidomic data from SRM 1950 human plasma and mouse liver tissues are available in Supplementary Data 1 and Supplementary Data 2, respectively. The table data for lipid enrichment analysis is available in Supplementary Data 4. Details of the raw data acquired on the pre-commercial ZenoTOF 8600, ZenoTOF 8600, and ZenoTOF 7600+ are provided in Supplementary Data 5. The lipidomics minimal reporting checklist for liver lipidomics is available in Supplementary Note 1.

## Code availability

MS-DIAL source code is available at https://github.com/systemsomicslab/MsdialWorkbench.

## Acknowledgments

This research was supported by the Japan Agency for Medical Research and Development (AMED) under Infectious Diseases Research and Infrastructure (JP25wm0325071, H.T.), the Japan Science and Technology Agency (JST) Exploratory Research for Advanced Technology (ERATO) (JPMJER2101 to H. T.), JST FOREST program (JPMJFR230H to H.T.), JST NBDC (JPMJND2305 to H.T.), JST ASPIRE program (JPMJAP2505 to H.T.), the JSPS KAKENHI (24K02011, 24H00043, 24H00392, 24K21269, 25H01425, and 25H01426 to H.T.), the Inserm project (international research project AtypicoLipid to T.H. and H.T.), and Technologically Advanced research through Marriage of Agriculture and engineering as Groundbreaking Organization (TAMAGO to H.T. and J.M.).

## Author Contributions

L.D. and H.T. designed the study. T.H. and H.T. organized the lipidomics experiments and the relevant data analyses. Y.M., Mikiko T., and H.T. developed the MS-DIAL environment. K.T., R.Y., M.H., and J.M. performed biological experiments. K.T., B.B., T.O., T.H., and H.T. analyzed the data. Manami T., U.T., G.I., D.C., P.B., A.C., N.B., and L.D. organized the ZT Scan DIA instruments and performed the LC-MS/MS analyses. H.T. wrote the manuscript, and G.I. and T.H. improved the draft. All authors have thoroughly discussed this project and helped improve the manuscript.

