## Supplementary Figures for "A dual-dimensional deconvolution environment for ZT Scan DIA in metabolomics"

Supplementary Figure 1

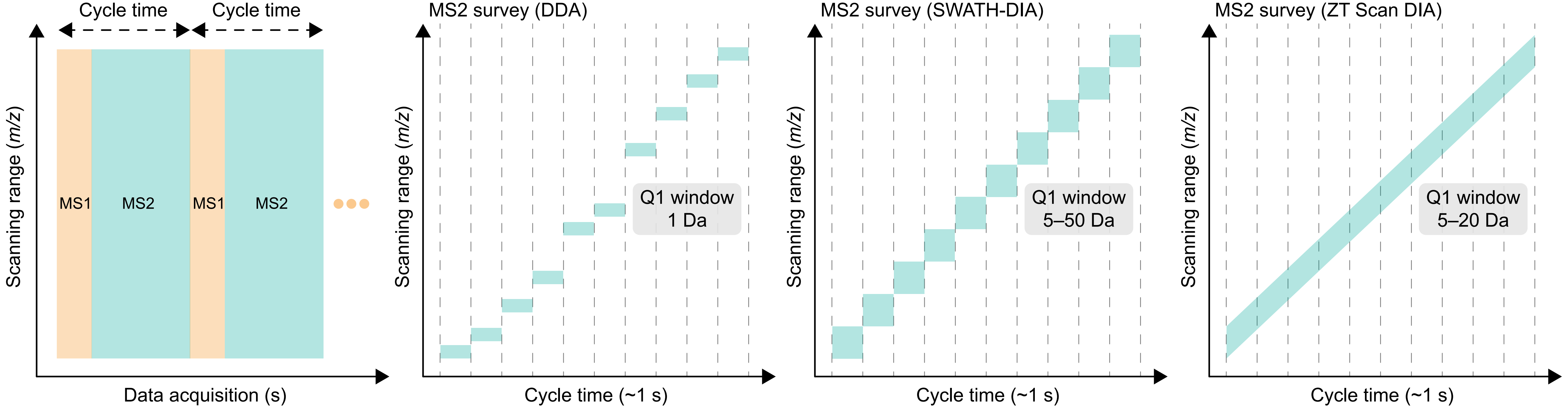

Supplementary Figure 2

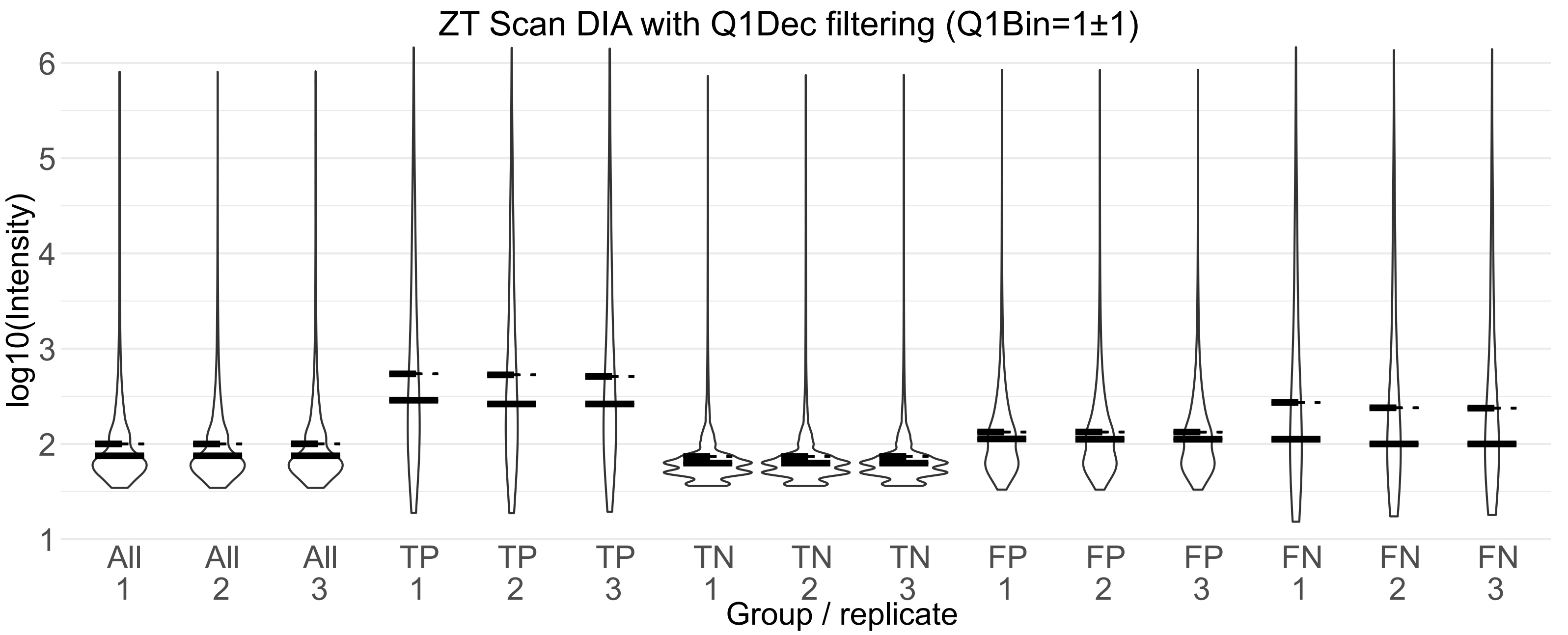

Supplementary Figure 3

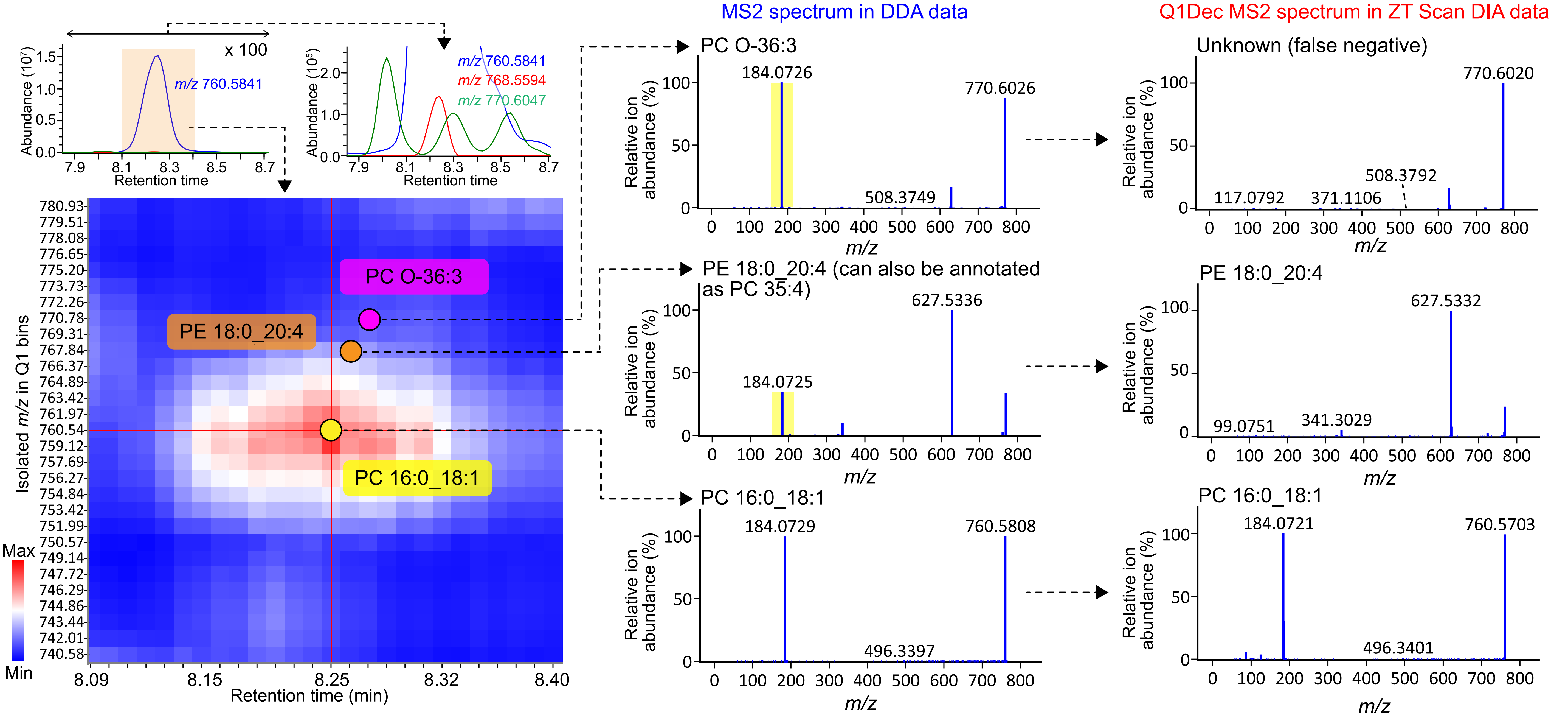

Supplementary Figure 4

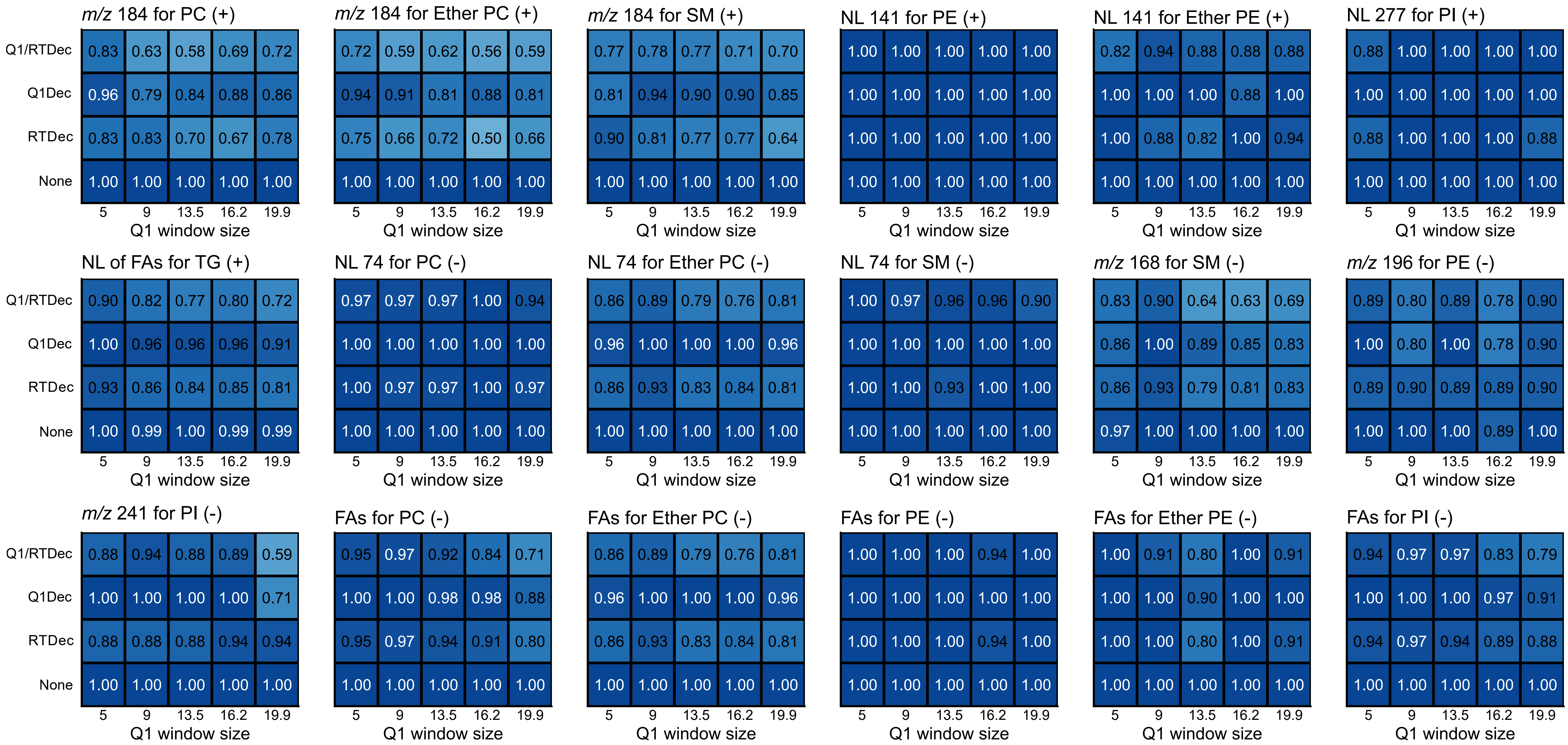

Supplementary Figure 5

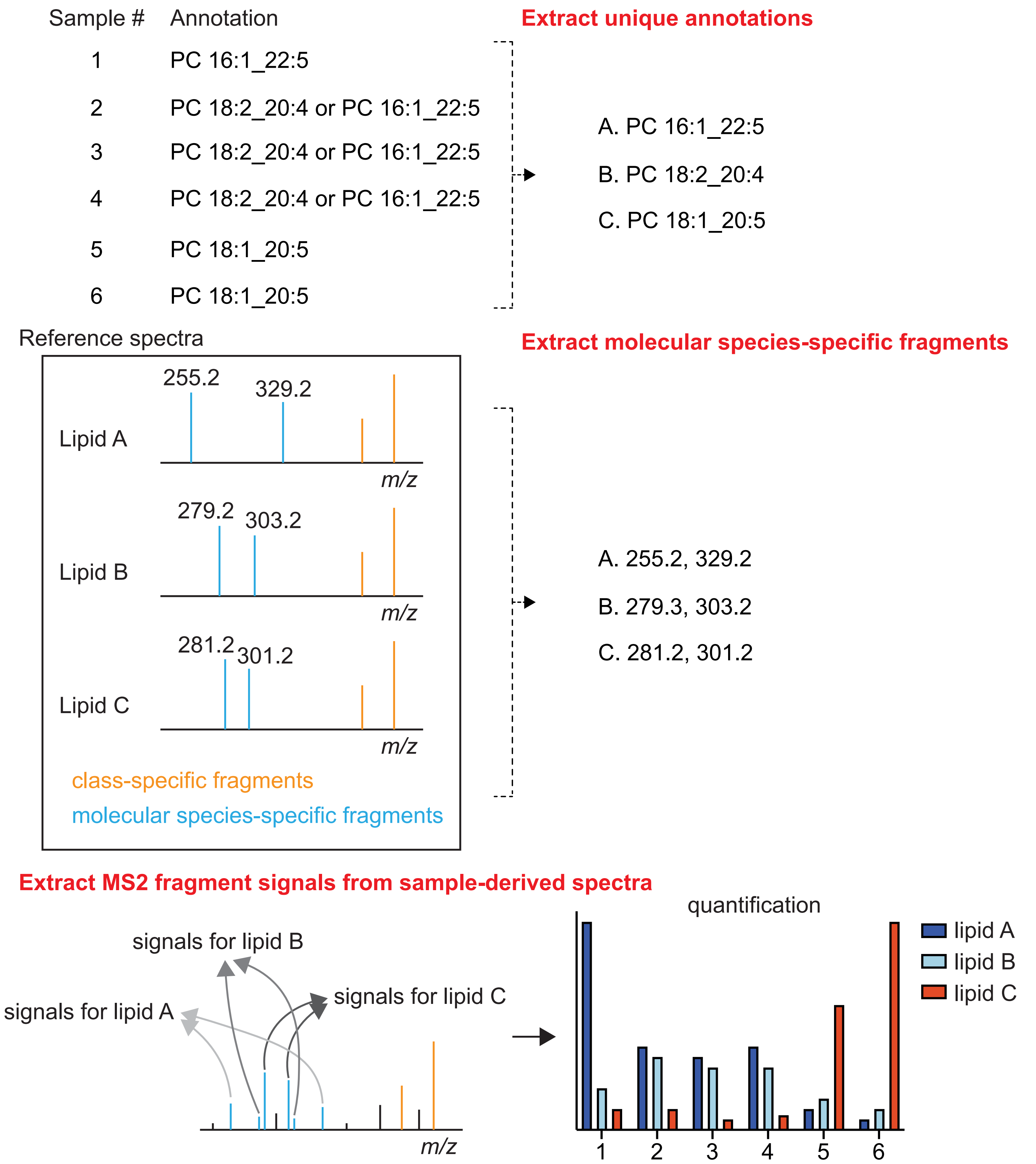

Supplementary Figure 6

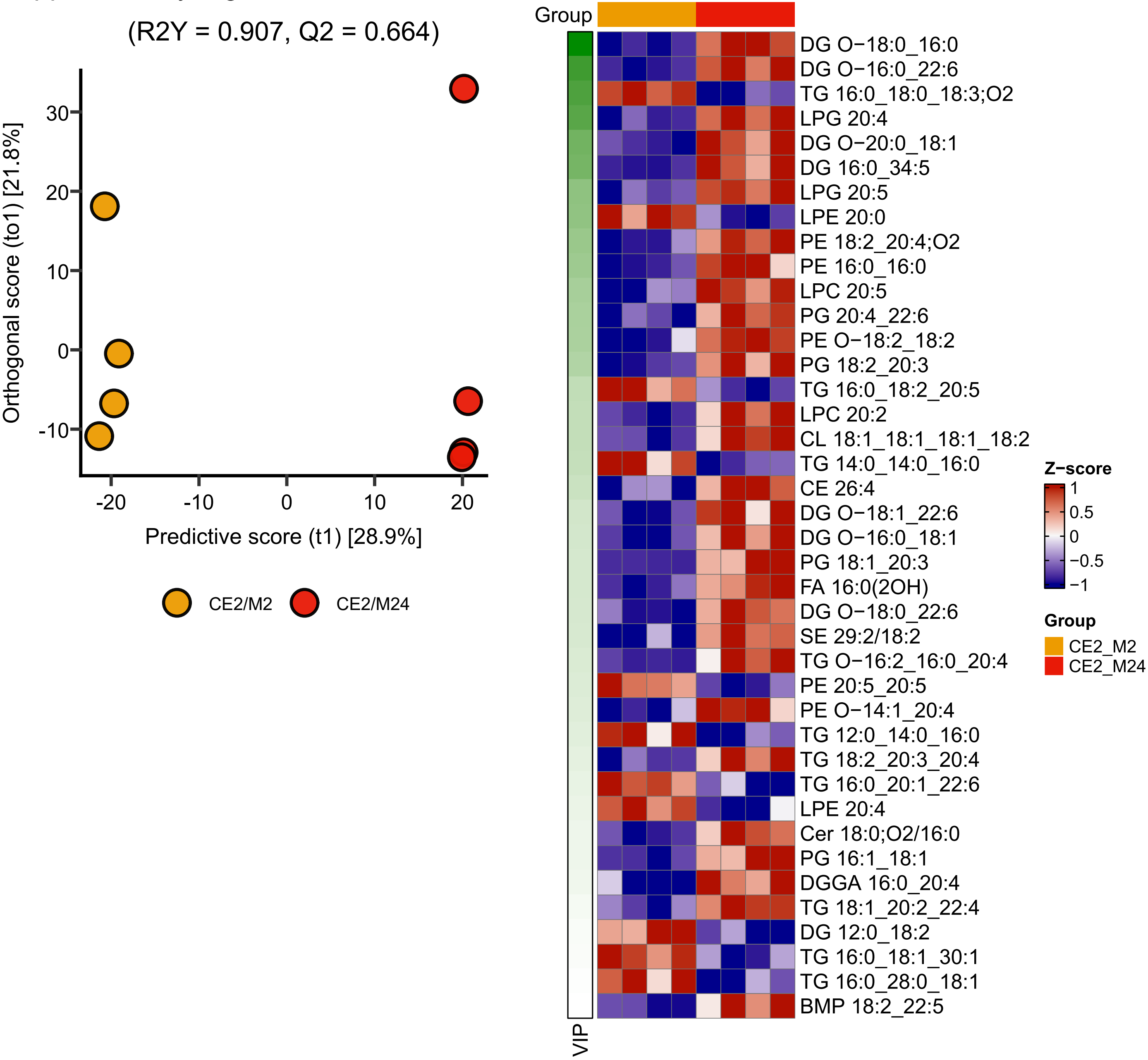

Supplementary Figure 7

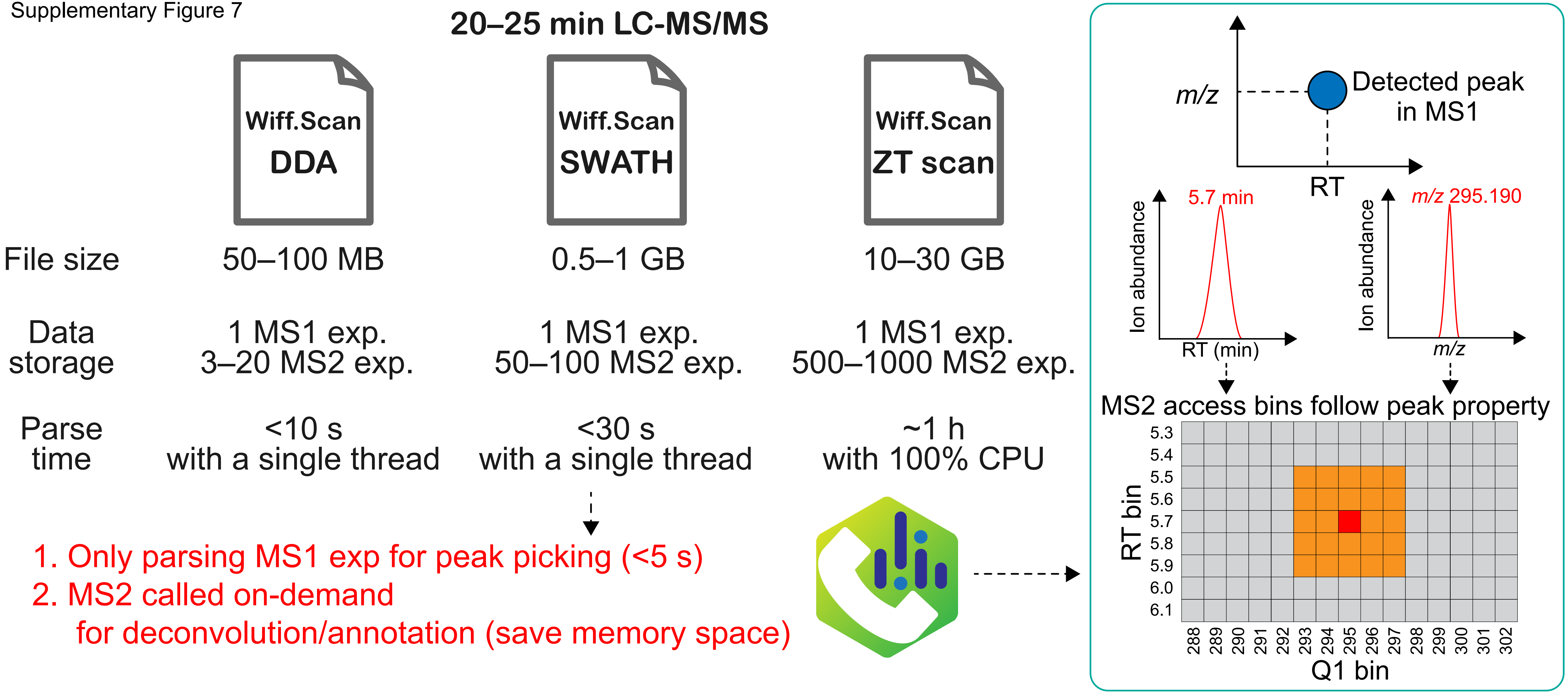

Supplementary Figure 8

### Spectrum-centric deconvolution

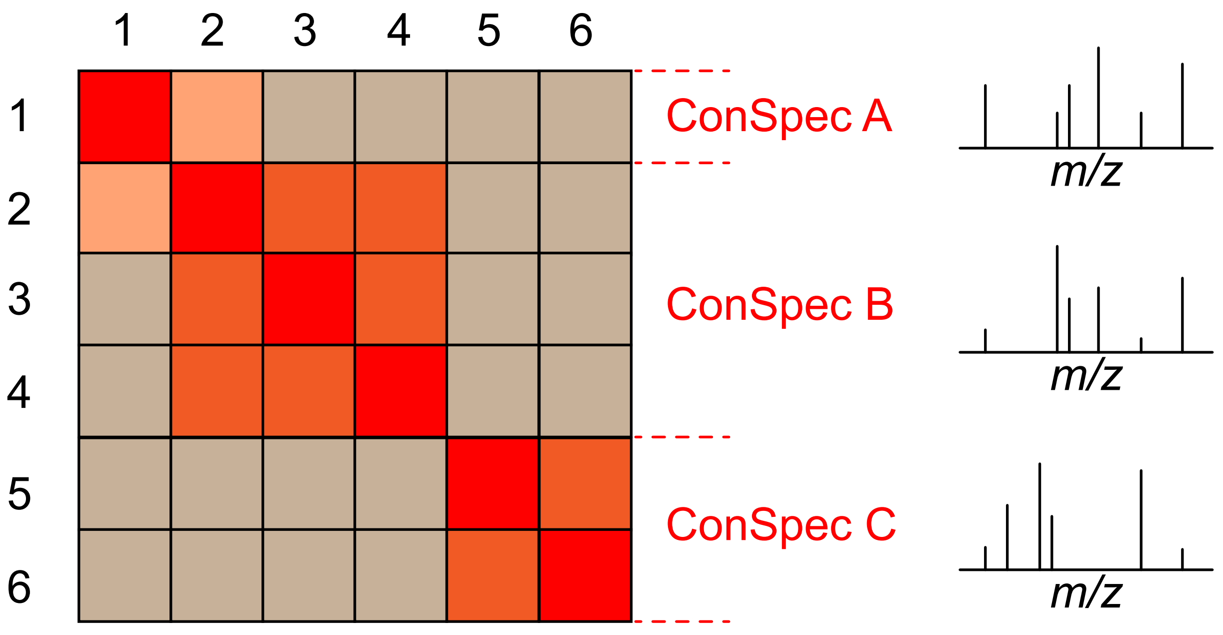


ConSpec: Consensus spectrum
-Peak summed by 0.5 Da bin (default)
-Dot product > 0.8 (default)
-Shared by 100% samples of a group (default)
 or *m/z* in Ref. spec (>0.5% rel.abs) (option)

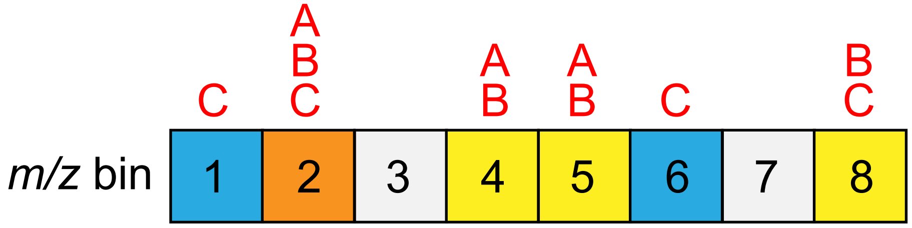


Unique mass definition and MS2 quantification
-mapping *m/z* of 70% rel.abs in 0.5 Da bin (default)
-metabolites grouping (AB and C in the illustration)
-calculate MS2 peak height/area by unique masses
