## Supplementary Note 1 for "A dual-dimensional deconvolution environment for ZT Scan DIA in metabolomics"

### Contents of Report

Created by https://lipidomicstandards.org, version v2.5.0

**Separation Workflow** 5

### Separation Workflow

#### Overall study design

| | | | |
|---|---|---|---|
| Title of the study | Mouse liver aging lipidome analysis in CE2 and HFD dietary conditions | | |
| Document creation date | 08/20/2025 | Corresponding Email | |
| Principal investigator | Hiroshi Tsugawa | Is the workflow targeted or untargeted? | Untargeted |
| Institution | Tokyo University of Agriculture and Technology | Clinical | No |

#### Lipid extraction

| | | | |
|---|---|---|---|
| Extraction method | 2-phase system | Were internal standards used? | Yes |
| pH adjustment | None | Deposition method | Added IS mixture in the extraction solvent |
| 2-phase system | MTBE | Internal standards used | EquiSPLASH and FA 16:0-d3 and FA 18:0-d3 |

#### Analytical platform

| | | | |
|---|---|---|---|
| Ionizaton additives | Ammonium acetate | Resolution at m/z 200 at $MS^1$ | 26232 |
| Number of separation dimensions | One dimension | Mass accuracy in ppm at $MS^1$ | 0.70434 |
| Separation type 1 | LC | Recording mode of raw data at $MS^1$ | Profile mode |
| Separation mode 1 (liquid) | RP | Mass window for precursor ion isolation (in Da total isolation window) | 1 |
| Detector | Mass spectrometer | Mass resolution for detected ion at $MS^2$ | High resolution |
| MS type | QTOF | Resolution at m/z 200 at $MS^2$ | 31418 |
| MS vendor | SCIEX | Mass accuracy in ppm at $MS^2$ | 0.07117 |
| Ion source | ESI | Recording mode of raw data at $MS^2$ | Profile mode |
| MS Level | $MS^1$, $MS^2$ | Was/Were additional dimension/techniques used | No |
| Mass resolution for detected ion at $MS^1$ | High resolution | | |

#### Quality control

| | | | |
|---|---|---|---|
| Blanks | Yes | Quality control | Yes |
| Type of Blanks | Extraction blank | Type of QC sample | Reference material |

##### Method qualification and validation

| | | | |
|---|---|---|---|
| Method validation | Yes | Precision | Yes |
| Lipid recovery | Yes | Accuracy | Yes |
| Dynamic quantification range | No | Guidelines followed | None |
| Limit of quantitation (LOQ)/Limit of detection (LOD) | No | | |

##### Reporting

| | | | |
|---|---|---|---|
| Are reported raw data uploaded into repository? | Yes | Summary data | Quantification and identification data |
| Link to repository / ID to entry | MB-POST repository (https://repository.massbank.jp/) /MPST000090.2 | Raw data upload | Yes |
| Are metadata available? | Yes | Additional comments | The raw data is available under the index of MPST000090.2 at the MB-POST repository (https://repository.massbank.jp/) . The resolution and accuracy (ppm) for MS1 are those of m/z 118.0863 in positive ion mode which were determined by the mass calibration in SCIEX OS. In addition, the resolution and accuracy for MS2 are those of m/z 185.1285 in positive ion mode, which were also determined by the mass calibration in SCIEX OS. |

#### Sample Descriptions

##### Mouse liver / Mouse / Tissues (e.g., liver, heart, brain)

| | | | |
|---|---|---|---|
| Tissue type | Liver | Time to freeze | between 5 and 10 minutes |
| Storage and collection conditions | Available | Snap freezing in liquid N2 | Yes |
| Sample homogenization | Yes | Storage temperature | -80 °C |
| Sample homogenization solvent | Methanol | Storage time (month) | 36 |
| Provided preanalytical information | Time to freeze, Storage time (month), Freeze-thaw cycles | Freeze-thaw cycles | 0 |
| Temperature handling original sample | 4-8 °C | Additives | None |
| Instant sample preparation | Yes | | |

### Lipid Class Descriptions

#### 1) FA / Lipid identification

| | | | |
|---|---|---|---|
| Lipid class | FA | Limit of detection | No |
| MS Level for identification | $MS^1$ | RT verified by standard | Yes |
| Identification level | Species level | Separation of isobaric/isomeric interferece confirmed | Yes |
| Isotope correction at $MS^1$ | No | Model for separation prediction | Yes |
| $MS^1$ verified by standard | Yes | Lipid Identification Software | MS-DIAL |
| Background check at $MS^1$ | Yes | Data manipulation | Smoothing, Centroiding |
| Did you presume assumptions for identification? | No | Nomenclature for intact lipid molecule | Yes |

#### 1) FA / Lipid quantification

| | | | |
|---|---|---|---|
| Quantitative | Yes | Type I isotope correction | No |
| MS Level for quantification | $MS^1$ | Limit of quantification | No |
| Internal lipid standard(s) $MS^1$ | | Normalization to reference | No |

| Internal standard | Endogenous subclass |
|---|---|
| FA 18:0(d3) | FA subclass |

| | | | |
|---|---|---|---|
| Type of quantification | Internal standard amount | Lipid Quantification Software | MS-DIAL |
| Response correction | No | Batch correction | No |

#### 2) DG / Lipid identification

| | | | |
|---|---|---|---|
| Lipid class | DG | Did you presume assumptions for identification? | No |
| MS Level for identification | $MS^1$, $MS^2$ | Limit of detection | No |
| Identification level | Molecular species level | RT verified by standard | Yes |
| Isotope correction at $MS^1$ | No | Separation of isobaric/isomeric interferece confirmed | Yes |
| Fragments for identification | | Model for separation prediction | Yes |

| Fragment name |
|---|
| Dehydro-monoacyl glycerols |
| Neutral loss of H2O |

| | | | |
|---|---|---|---|
| Isotope correction at $MS^2$ | No | Lipid Identification Software | MS-DIAL |
| $MS^1$ verified by standard | Yes | Data manipulation | Smoothing, Centroiding |
| $MS^2$ verified by standard | Yes | Nomenclature for intact lipid molecule | Yes |
| Background check at $MS^1$ | Yes | Nomenclature for fragment ions | No |
| Background check at $MS^2$ | No | | |

#### 2) DG / Lipid quantification

| | | | |
|---|---|---|---|
| Quantitative | Yes | Type I isotope correction | No |
| MS Level for quantification | $MS^1$ | Limit of quantification | No |
| Internal lipid standard(s) $MS^1$ | | Normalization to reference | No |

| Internal standard | Endogenous subclass |
|---|---|
| DG 15:0_18:1(d7) | DG subclass |

| | | | |
|---|---|---|---|
| Type of quantification | Internal standard amount | Lipid Quantification Software | MS-DIAL |
| Response correction | No | Batch correction | No |

#### 3) DGDG / Lipid identification

| | | | |
|---|---|---|---|
| Lipid class | DGDG | Did you presume assumptions for identification? | No |
| MS Level for identification | $MS^1$, $MS^2$ | Limit of detection | No |
| Identification level | Molecular species level | RT verified by standard | Yes |
| Isotope correction at $MS^1$ | No | Separation of isobaric/isomeric interferece confirmed | Yes |
| Fragments for identification | | Model for separation prediction | Yes |

| Fragment name |
|---|
| Fatty acid fragment |
| Neutral loss of fatty acids |

| | | | |
|---|---|---|---|
| Isotope correction at $MS^2$ | No | Lipid Identification Software | MS-DIAL |
| $MS^1$ verified by standard | No | Data manipulation | Smoothing, Centroiding |
| $MS^2$ verified by standard | No | Nomenclature for intact lipid molecule | Yes |
| Background check at $MS^1$ | Yes | Nomenclature for fragment ions | No |
| Background check at $MS^2$ | No | | |

#### 3) DGDG / Lipid quantification

| | | | |
|---|---|---|---|
| Quantitative | Yes | Type I isotope correction | No |
| MS Level for quantification | $MS^1$ | Limit of quantification | No |
| Internal lipid standard(s) $MS^1$ | | Normalization to reference | No |

| Internal standard | Endogenous subclass |
|---|---|
| LPC 18:1(d7) | DGDG subclass |

| | | | |
|---|---|---|---|
| Type of quantification | Internal standard amount | Lipid Quantification Software | MS-DIAL |
| Response correction | No | Batch correction | No |

#### 4) Sulfonolipid (SL) / Lipid identification

| | | | |
|---|---|---|---|
| Lipid class | Sulfonolipid (SL) | Did you presume assumptions for identification? | No |
| MS Level for identification | $MS^1$, $MS^2$ | Limit of detection | No |
| Identification level | Molecular species level | RT verified by standard | Yes |
| Isotope correction at $MS^1$ | No | Separation of isobaric/isomeric interferece confirmed | Yes |
| Fragments for identification | | Model for separation prediction | Yes |

| Fragment name |
|---|
| Sulfite |
| Neutral loss of N-acyl chain |

| | | | |
|---|---|---|---|
| Isotope correction at $MS^2$ | No | Lipid Identification Software | MS-DIAL |
| $MS^1$ verified by standard | No | Data manipulation | Smoothing, Centroiding |
| $MS^2$ verified by standard | No | Nomenclature for intact lipid molecule | Yes |
| Background check at $MS^1$ | Yes | Nomenclature for fragment ions | No |
| Background check at $MS^2$ | No | | |

#### 4) Sulfonolipid (SL) / Lipid quantification

| | | | |
|---|---|---|---|
| Quantitative | Yes | Type I isotope correction | No |
| MS Level for quantification | $MS^1$ | Limit of quantification | No |
| Internal lipid standard(s) $MS^1$ | | Normalization to reference | No |

| Internal standard | Endogenous subclass |
|---|---|
| Cer 18:1;2O/15:0(d7) | SL subclass |

| | | | |
|---|---|---|---|
| Type of quantification | Internal standard amount | Lipid Quantification Software | MS-DIAL |
| Response correction | No | Batch correction | No |

#### 5) Acyl diacylglyceryl glucuronide(ADGGA) / Lipid identification

| | | | |
|---|---|---|---|
| Lipid class | Acyl diacylglyceryl glucuronide(ADGGA) | Did you presume assumptions for identification? | No |
| MS Level for identification | $MS^1$, $MS^2$ | Limit of detection | No |
| Identification level | Molecular species level | RT verified by standard | Yes |
| Isotope correction at $MS^1$ | No | Separation of isobaric/isomeric interferece confirmed | Yes |
| Fragments for identification | | Model for separation prediction | Yes |

| Fragment name |
|---|
| Fatty acid fragment |

| | | | |
|---|---|---|---|
| Isotope correction at $MS^2$ | No | Lipid Identification Software | MS-DIAL |
| $MS^1$ verified by standard | No | Data manipulation | Smoothing, Centroiding |
| $MS^2$ verified by standard | No | Nomenclature for intact lipid molecule | Yes |
| Background check at $MS^1$ | Yes | Nomenclature for fragment ions | No |
| Background check at $MS^2$ | No | | |

#### 5) Acyl diacylglyceryl glucuronide(ADGGA) / Lipid quantification

| | | | |
|---|---|---|---|
| Quantitative | Yes | Type I isotope correction | No |
| MS Level for quantification | $MS^1$ | Limit of quantification | No |
| Internal lipid standard(s) $MS^1$ | | Normalization to reference | No |

| Internal standard | Endogenous subclass |
|---|---|
| LPC 18:1(d7) | ADGGA subclass |

| | | | |
|---|---|---|---|
| Type of quantification | Internal standard amount | Lipid Quantification Software | MS-DIAL |
| Response correction | No | Batch correction | No |

#### 6) Acylcarnitine (CAR) / Lipid identification

| | | | |
|---|---|---|---|
| Lipid class | Acylcarnitine (CAR) | Did you presume assumptions for identification? | No |
| MS Level for identification | $MS^1$, $MS^2$ | Limit of detection | No |
| Identification level | Molecular species level | RT verified by standard | Yes |
| Isotope correction at $MS^1$ | No | Separation of isobaric/isomeric interferece confirmed | Yes |
| Fragments for identification | | Model for separation prediction | Yes |

| Fragment name |
|---|
| Characteristic fragment (C4H5O2+) |

| | | | |
|---|---|---|---|
| Isotope correction at $MS^2$ | No | Lipid Identification Software | MS-DIAL |
| $MS^1$ verified by standard | No | Data manipulation | Smoothing, Centroiding |
| $MS^2$ verified by standard | No | Nomenclature for intact lipid molecule | Yes |
| Background check at $MS^1$ | Yes | Nomenclature for fragment ions | No |
| Background check at $MS^2$ | No | | |

#### 6) Acylcarnitine (CAR) / Lipid quantification

| | | | |
|---|---|---|---|
| Quantitative | Yes | Type I isotope correction | No |
| MS Level for quantification | $MS^1$ | Limit of quantification | No |
| Internal lipid standard(s) $MS^1$ | | Normalization to reference | No |

| Internal standard | Endogenous subclass |
|---|---|
| LPC 18:1(d7) | CAR subclass |

| | | | |
|---|---|---|---|
| Type of quantification | Internal standard amount | Lipid Quantification Software | MS-DIAL |
| Response correction | No | Batch correction | No |

#### 7) Acylhexosyl brassicasterol (AHexBRS) / Lipid identification

| | | | |
|---|---|---|---|
| Lipid class | Acylhexosyl brassicasterol (AHexBRS) | Did you presume assumptions for identification? | No |
| MS Level for identification | $MS^1$, $MS^2$ | Limit of detection | No |
| Identification level | Molecular species level | RT verified by standard | Yes |
| Isotope correction at $MS^1$ | No | Separation of isobaric/isomeric interferece confirmed | Yes |
| Fragments for identification | | Model for separation prediction | Yes |
| Isotope correction at $MS^2$ | No | Lipid Identification Software | MS-DIAL |
| $MS^1$ verified by standard | No | Data manipulation | Smoothing, Centroiding |
| $MS^2$ verified by standard | No | Nomenclature for intact lipid molecule | Yes |
| Background check at $MS^1$ | Yes | Nomenclature for fragment ions | No |
| Background check at $MS^2$ | No | | |

Fragments for identification:

| Fragment name |
|---|
| Fatty acid fragment |

#### 7) Acylhexosyl brassicasterol (AHexBRS) / Lipid quantification

| | | | |
|---|---|---|---|
| Quantitative | Yes | Type I isotope correction | No |
| MS Level for quantification | $MS^1$ | Limit of quantification | No |
| Internal lipid standard(s) $MS^1$ | | Normalization to reference | No |
| Type of quantification | Internal standard amount | Lipid Quantification Software | MS-DIAL |
| Response correction | No | Batch correction | No |

Internal lipid standard(s) $MS^1$:

| Internal standard | Endogenous subclass |
|---|---|
| CE 18:1(d7) | AHexBRS subclass |

#### 8) Acylhexosyl campesterol (AHexCAS) / Lipid identification

| | | | |
|---|---|---|---|
| Lipid class | Acylhexosyl campesterol (AHexCAS) | Did you presume assumptions for identification? | No |
| MS Level for identification | $MS^1$, $MS^2$ | Limit of detection | No |
| Identification level | Molecular species level | RT verified by standard | Yes |
| Isotope correction at $MS^1$ | No | Separation of isobaric/isomeric interferece confirmed | Yes |
| Fragments for identification | | Model for separation prediction | Yes |
| Isotope correction at $MS^2$ | No | Lipid Identification Software | MS-DIAL |
| $MS^1$ verified by standard | No | Data manipulation | Smoothing, Centroiding |
| $MS^2$ verified by standard | No | Nomenclature for intact lipid molecule | Yes |
| Background check at $MS^1$ | Yes | Nomenclature for fragment ions | No |
| Background check at $MS^2$ | No | | |

Fragments for identification:

| Fragment name |
|---|
| Fatty acid fragment |

#### 8) Acylhexosyl campesterol (AHexCAS) / Lipid quantification

| | | | |
|---|---|---|---|
| Quantitative | Yes | Type I isotope correction | No |
| MS Level for quantification | $MS^1$ | Limit of quantification | No |
| Internal lipid standard(s) $MS^1$ | | Normalization to reference | No |

| Internal standard | Endogenous subclass |
|---|---|
| CE 18:1(d7) | AHexCAS subclass |

| | | | |
|---|---|---|---|
| Type of quantification | Internal standard amount | Lipid Quantification Software | MS-DIAL |
| Response correction | No | Batch correction | No |

#### 9) Acylhexosyl cholesterol (AHexCS) / Lipid identification

| | | | |
|---|---|---|---|
| Lipid class | Acylhexosyl cholesterol (AHexCS) | Did you presume assumptions for identification? | No |
| MS Level for identification | $MS^1$, $MS^2$ | Limit of detection | No |
| Identification level | Molecular species level | RT verified by standard | Yes |
| Isotope correction at $MS^1$ | No | Separation of isobaric/isomeric interferece confirmed | Yes |
| Fragments for identification | | Model for separation prediction | Yes |

| Fragment name |
|---|
| Fatty acid fragment |

| | | | |
|---|---|---|---|
| Isotope correction at $MS^2$ | No | Lipid Identification Software | MS-DIAL |
| $MS^1$ verified by standard | No | Data manipulation | Smoothing, Centroiding |
| $MS^2$ verified by standard | No | Nomenclature for intact lipid molecule | Yes |
| Background check at $MS^1$ | Yes | Nomenclature for fragment ions | No |
| Background check at $MS^2$ | No | | |

#### 9) Acylhexosyl cholesterol (AHexCS) / Lipid quantification

| | | | |
|---|---|---|---|
| Quantitative | Yes | Type I isotope correction | No |
| MS Level for quantification | $MS^1$ | Limit of quantification | No |
| Internal lipid standard(s) $MS^1$ | | Normalization to reference | No |

| Internal standard | Endogenous subclass |
|---|---|
| CE 18:1(d7) | AHexCS subclass |

| | | | |
|---|---|---|---|
| Type of quantification | Internal standard amount | Lipid Quantification Software | MS-DIAL |
| Response correction | No | Batch correction | No |

#### 10) Acylhexosyl sitosterol (AHexSIS) / Lipid identification

| | | | |
|---|---|---|---|
| Lipid class | Acylhexosyl sitosterol (AHexSIS) | Did you presume assumptions for identification? | No |
| MS Level for identification | $MS^1$, $MS^2$ | Limit of detection | No |
| Identification level | Molecular species level | RT verified by standard | Yes |
| Isotope correction at $MS^1$ | No | Separation of isobaric/isomeric interferece confirmed | Yes |
| Fragments for identification | | Model for separation prediction | Yes |
| Isotope correction at $MS^2$ | No | Lipid Identification Software | MS-DIAL |
| $MS^1$ verified by standard | No | Data manipulation | Smoothing, Centroiding |
| $MS^2$ verified by standard | No | Nomenclature for intact lipid molecule | Yes |
| Background check at $MS^1$ | Yes | Nomenclature for fragment ions | No |
| Background check at $MS^2$ | No | | |

| Fragment name |
|---|
| Fatty acid fragment |

#### 10) Acylhexosyl sitosterol (AHexSIS) / Lipid quantification

| | | | |
|---|---|---|---|
| Quantitative | Yes | Type I isotope correction | No |
| MS Level for quantification | $MS^1$ | Limit of quantification | No |
| Internal lipid standard(s) $MS^1$ | | Normalization to reference | No |
| Type of quantification | Internal standard amount | Lipid Quantification Software | MS-DIAL |
| Response correction | No | Batch correction | No |

| Internal standard | Endogenous subclass |
|---|---|
| CE 18:1(d7) | AHexSIS subclass |

#### 11) Acylhexosyl stigmasterol (AHexSTS) / Lipid identification

| | | | |
|---|---|---|---|
| Lipid class | Acylhexosyl stigmasterol (AHexSTS) | Did you presume assumptions for identification? | No |
| MS Level for identification | $MS^1$, $MS^2$ | Limit of detection | No |
| Identification level | Molecular species level | RT verified by standard | Yes |
| Isotope correction at $MS^1$ | No | Separation of isobaric/isomeric interferece confirmed | Yes |
| Fragments for identification | | Model for separation prediction | Yes |
| Isotope correction at $MS^2$ | No | Lipid Identification Software | MS-DIAL |
| $MS^1$ verified by standard | No | Data manipulation | Smoothing, Centroiding |
| $MS^2$ verified by standard | No | Nomenclature for intact lipid molecule | Yes |
| Background check at $MS^1$ | Yes | Nomenclature for fragment ions | No |
| Background check at $MS^2$ | No | | |

| Fragment name |
|---|
| Fatty acid fragment |

#### 11) Acylhexosyl stigmasterol (AHexSTS) / Lipid quantification

| | | | |
|---|---|---|---|
| Quantitative | Yes | Type I isotope correction | No |
| MS Level for quantification | $MS^1$ | Limit of quantification | No |
| Internal lipid standard(s) $MS^1$ | | Normalization to reference | No |

| Internal standard | Endogenous subclass |
|---|---|
| CE 18:1(d7) | AHexSTS subclass |

| | | | |
|---|---|---|---|
| Type of quantification | Internal standard amount | Lipid Quantification Software | MS-DIAL |
| Response correction | No | Batch correction | No |

#### 12) Acylhexosylceramide (AHexCer) / Lipid identification

| | | | |
|---|---|---|---|
| Lipid class | Acylhexosylceramide (AHexCer) | Did you presume assumptions for identification? | No |
| MS Level for identification | $MS^1$, $MS^2$ | Limit of detection | No |
| Identification level | Molecular species level | RT verified by standard | Yes |
| Isotope correction at $MS^1$ | No | Separation of isobaric/isomeric interferece confirmed | Yes |
| Fragments for identification | | Model for separation prediction | Yes |

| Fragment name |
|---|
| Neutral loss of fatty acid |
| Neutral loss of fatty acid and hexose |
| Fatty acid fragment |
| Neutral loss of N- acyl |
| Neutral loss of fatty acid and H2O |
| Neutral loss of N-acyl and H2O |

| | | | |
|---|---|---|---|
| Isotope correction at $MS^2$ | No | Lipid Identification Software | MS-DIAL |
| $MS^1$ verified by standard | No | Data manipulation | Smoothing, Centroiding |
| $MS^2$ verified by standard | No | Nomenclature for intact lipid molecule | Yes |
| Background check at $MS^1$ | Yes | Nomenclature for fragment ions | No |
| Background check at $MS^2$ | No | | |

#### 12) Acylhexosylceramide (AHexCer) / Lipid quantification

| | | | |
|---|---|---|---|
| Quantitative | Yes | Type I isotope correction | No |
| MS Level for quantification | $MS^1$ | Limit of quantification | No |
| Internal lipid standard(s) $MS^1$ | | Normalization to reference | No |

| Internal standard | Endogenous subclass |
|---|---|
| Cer 18:1;2O/15:0(d7) | AHexCer subclass |

| | | | |
|---|---|---|---|
| Type of quantification | Internal standard amount | Lipid Quantification Software | MS-DIAL |
| Response correction | No | Batch correction | No |

#### 13) Acylsphingomyelin (ASM) / Lipid identification

| | | | |
|---|---|---|---|
| Lipid class | Acylsphingomyelin (ASM) | Did you presume assumptions for identification? | No |
| MS Level for identification | $MS^1$, $MS^2$ | Limit of detection | No |
| Identification level | Molecular species level | RT verified by standard | Yes |
| Isotope correction at $MS^1$ | No | Separation of isobaric/isomeric interferece confirmed | Yes |
| Fragments for identification | | Model for separation prediction | Yes |

| Fragment name |
|---|
| Neutral loss of methyl |
| Neutral loss of methyl and fatty acid and H2O |
| Fatty acid fragment |
| Acyl amide |

| | | | |
|---|---|---|---|
| Isotope correction at $MS^2$ | No | Lipid Identification Software | MS-DIAL |
| $MS^1$ verified by standard | No | Data manipulation | Smoothing, Centroiding |
| $MS^2$ verified by standard | No | Nomenclature for intact lipid molecule | Yes |
| Background check at $MS^1$ | Yes | Nomenclature for fragment ions | No |
| Background check at $MS^2$ | No | | |

#### 13) Acylsphingomyelin (ASM) / Lipid quantification

| | | | |
|---|---|---|---|
| Quantitative | Yes | Type I isotope correction | No |
| MS Level for quantification | $MS^1$ | Limit of quantification | No |
| Internal lipid standard(s) $MS^1$ | | Normalization to reference | No |

| Internal standard | Endogenous subclass |
|---|---|
| SM 18:1;2O/18:1(d9) | ASM subclass |

| | | | |
|---|---|---|---|
| Type of quantification | Internal standard amount | Lipid Quantification Software | MS-DIAL |
| Response correction | No | Batch correction | No |

#### 14) BMP / Lipid identification

| | | | |
|---|---|---|---|
| Lipid class | BMP | Did you presume assumptions for identification? | No |
| MS Level for identification | $MS^1$, $MS^2$ | Limit of detection | No |
| Identification level | Molecular species level | RT verified by standard | Yes |
| Isotope correction at $MS^1$ | No | Separation of isobaric/isomeric interferece confirmed | Yes |
| Fragments for identification | | Model for separation prediction | Yes |

| Fragment name |
|---|
| Dehydro-monoacyl glycerols |
| Neutral loss of glycerophosphate |

| | | | |
|---|---|---|---|
| Isotope correction at $MS^2$ | No | Lipid Identification Software | MS-DIAL |
| $MS^1$ verified by standard | No | Data manipulation | Smoothing, Centroiding |
| $MS^2$ verified by standard | No | Nomenclature for intact lipid molecule | Yes |
| Background check at $MS^1$ | Yes | Nomenclature for fragment ions | No |
| Background check at $MS^2$ | No | | |

#### 14) BMP / Lipid quantification

| | | | |
|---|---|---|---|
| Quantitative | Yes | Type I isotope correction | No |
| MS Level for quantification | $MS^1$ | Limit of quantification | No |
| Internal lipid standard(s) $MS^1$ | | Normalization to reference | No |

| Internal standard | Endogenous subclass |
|---|---|
| PG 15:0_18:1(d7) | BMP subclass |

| | | | |
|---|---|---|---|
| Type of quantification | Internal standard amount | Lipid Quantification Software | MS-DIAL |
| Response correction | No | Batch correction | No |

#### 15) Brassicasterol / Lipid identification

| | | | |
|---|---|---|---|
| Lipid class | Brassicasterol | Limit of detection | No |
| MS Level for identification | $MS^1$ | RT verified by standard | Yes |
| Identification level | Species level | Separation of isobaric/isomeric interferece confirmed | Yes |
| Isotope correction at $MS^1$ | No | Model for separation prediction | Yes |
| $MS^1$ verified by standard | No | Lipid Identification Software | MS-DIAL |
| Background check at $MS^1$ | Yes | Data manipulation | Smoothing, Centroiding |
| Did you presume assumptions for identification? | No | Nomenclature for intact lipid molecule | Yes |

#### 15) Brassicasterol / Lipid quantification

| | | | |
|---|---|---|---|
| Quantitative | Yes | Type I isotope correction | No |
| MS Level for quantification | $MS^1$ | Limit of quantification | No |
| Internal lipid standard(s) $MS^1$ | | Normalization to reference | No |

| Internal standard | Endogenous subclass |
|---|---|
| CE 18:1(d7) | Brassicasterol |

| | | | |
|---|---|---|---|
| Type of quantification | Internal standard amount | Lipid Quantification Software | MS-DIAL |
| Response correction | No | Batch correction | No |

#### 16) Brassicasterol ester (BRSE) / Lipid identification

| | | | |
|---|---|---|---|
| Lipid class | Brassicasterol ester (BRSE) | Did you presume assumptions for identification? | No |
| MS Level for identification | $MS^1$, $MS^2$ | Limit of detection | No |
| Identification level | Molecular species level | RT verified by standard | Yes |
| Isotope correction at $MS^1$ | No | Separation of isobaric/isomeric interferece confirmed | Yes |
| Fragments for identification | | Model for separation prediction | Yes |

| Fragment name |
|---|
| Neutral loss of fatty acid |

| | | | |
|---|---|---|---|
| Isotope correction at $MS^2$ | No | Lipid Identification Software | MS-DIAL |
| $MS^1$ verified by standard | No | Data manipulation | Smoothing, Centroiding |
| $MS^2$ verified by standard | No | Nomenclature for intact lipid molecule | Yes |
| Background check at $MS^1$ | Yes | Nomenclature for fragment ions | No |
| Background check at $MS^2$ | No | | |

#### 16) Brassicasterol ester (BRSE) / Lipid quantification

| | | | |
|---|---|---|---|
| Quantitative | Yes | Type I isotope correction | No |
| MS Level for quantification | $MS^1$ | Limit of quantification | No |
| Internal lipid standard(s) $MS^1$ | | Normalization to reference | No |

| Internal standard | Endogenous subclass |
|---|---|
| CE 18:1(d7) | BRSE subclass |

| | | | |
|---|---|---|---|
| Type of quantification | Internal standard amount | Lipid Quantification Software | MS-DIAL |
| Response correction | No | Batch correction | No |

#### 17) Campesterol ester (CASE) / Lipid identification

| | | | |
|---|---|---|---|
| Lipid class | Campesterol ester (CASE) | Did you presume assumptions for identification? | No |
| MS Level for identification | $MS^1$, $MS^2$ | Limit of detection | No |
| Identification level | Molecular species level | RT verified by standard | Yes |
| Isotope correction at $MS^1$ | No | Separation of isobaric/isomeric interferece confirmed | Yes |
| Fragments for identification | | Model for separation prediction | Yes |

| Fragment name |
|---|
| Neutral loss of fatty acid |

| | | | |
|---|---|---|---|
| Isotope correction at $MS^2$ | No | Lipid Identification Software | MS-DIAL |
| $MS^1$ verified by standard | No | Data manipulation | Smoothing, Centroiding |
| $MS^2$ verified by standard | No | Nomenclature for intact lipid molecule | Yes |
| Background check at $MS^1$ | Yes | Nomenclature for fragment ions | No |
| Background check at $MS^2$ | No | | |

#### 17) Campesterol ester (CASE) / Lipid quantification

| | | | |
|---|---|---|---|
| Quantitative | Yes | Type I isotope correction | No |
| MS Level for quantification | $MS^1$ | Limit of quantification | No |
| Internal lipid standard(s) $MS^1$ | | Normalization to reference | No |

| Internal standard | Endogenous subclass |
|---|---|
| CE 18:1(d7) | CASE subclass |

| | | | |
|---|---|---|---|
| Type of quantification | Internal standard amount | Lipid Quantification Software | MS-DIAL |
| Response correction | No | Batch correction | No |

#### 18) CL / Lipid identification

| | | | |
|---|---|---|---|
| Lipid class | CL | Did you presume assumptions for identification? | No |
| MS Level for identification | $MS^1$, $MS^2$ | Limit of detection | No |
| Identification level | Molecular species level | RT verified by standard | Yes |
| Isotope correction at $MS^1$ | No | Separation of isobaric/isomeric interferece confirmed | Yes |
| Fragments for identification | | Model for separation prediction | Yes |

| Fragment name |
|---|
| Phosphoglycerol - H2O |
| Fatty acid fragment |
| Phosphatidic acid |

| | | | |
|---|---|---|---|
| Isotope correction at $MS^2$ | No | Lipid Identification Software | MS-DIAL |
| $MS^1$ verified by standard | No | Data manipulation | Smoothing, Centroiding |
| $MS^2$ verified by standard | No | Nomenclature for intact lipid molecule | Yes |
| Background check at $MS^1$ | Yes | Nomenclature for fragment ions | No |
| Background check at $MS^2$ | No | | |

#### 18) CL / Lipid quantification

| | | | |
|---|---|---|---|
| Quantitative | Yes | Type I isotope correction | No |
| MS Level for quantification | $MS^1$ | Limit of quantification | No |
| Internal lipid standard(s) $MS^1$ | | Normalization to reference | No |

| Internal standard | Endogenous subclass |
|---|---|
| PG 15:0_18:1(d7) | CL subclass |

| | | | |
|---|---|---|---|
| Type of quantification | Internal standard amount | Lipid Quantification Software | MS-DIAL |
| Response correction | No | Batch correction | No |

#### 19) Ceramide alpha-hydroxy fatty acid-dihydrosphingosine (Cer_ADS) / Lipid identification

| | | | |
|---|---|---|---|
| Lipid class | Ceramide alpha-hydroxy fatty acid-dihydrosphingosine (Cer_ADS) | Did you presume assumptions for identification? | No |
| MS Level for identification | $MS^1$, $MS^2$ | Limit of detection | No |
| Identification level | Molecular species level | RT verified by standard | Yes |
| Isotope correction at $MS^1$ | No | Separation of isobaric/isomeric interferece confirmed | Yes |
| Fragments for identification | | Model for separation prediction | Yes |

| Fragment name |
|---|
| Neutral loss of CH4O2 |
| Neutral loss of H2O |
| Sphinganine |
| Sphinganine -H2O fragment |
| Oxidized acyl -2H fragment |
| Oxidized acyl -CH2O fragment |
| Sphinganine -C2H7NO fragment |

| | | | |
|---|---|---|---|
| Isotope correction at $MS^2$ | No | Lipid Identification Software | MS-DIAL |
| $MS^1$ verified by standard | No | Data manipulation | Smoothing, Centroiding |
| $MS^2$ verified by standard | No | Nomenclature for intact lipid molecule | Yes |
| Background check at $MS^1$ | Yes | Nomenclature for fragment ions | No |
| Background check at $MS^2$ | No | | |

#### 19) Ceramide alpha-hydroxy fatty acid-dihydrosphingosine (Cer_ADS) / Lipid quantification

| | | | |
|---|---|---|---|
| Quantitative | Yes | Type I isotope correction | No |
| MS Level for quantification | $MS^1$ | Limit of quantification | No |
| Internal lipid standard(s) $MS^1$ | | Normalization to reference | No |

| Internal standard | Endogenous subclass |
|---|---|
| Cer 18:1;2O/15:0(d7) | Cer_ADS subclass |

| | | | |
|---|---|---|---|
| Type of quantification | Internal standard amount | Lipid Quantification Software | MS-DIAL |
| Response correction | No | Batch correction | No |

#### 20) Ceramide alpha-hydroxy fatty acid-phytospingosine (Cer_AP) / Lipid identification

| | | | |
|---|---|---|---|
| Lipid class | Ceramide alpha-hydroxy fatty acid-phytospingosine (Cer_AP) | Did you presume assumptions for identification? | No |
| MS Level for identification | $MS^1$, $MS^2$ | Limit of detection | No |
| Identification level | Molecular species level | RT verified by standard | Yes |
| Isotope correction at $MS^1$ | No | Separation of isobaric/isomeric interferece confirmed | Yes |
| Fragments for identification | | Model for separation prediction | Yes |

| Fragment name |
|---|
| Oxidized fatty acyl -CH2O fragment |
| Oxidized fatty acyl +O fragment |
| Oxidized fatty acyl +C3H5NO fragment |

| | | | |
|---|---|---|---|
| Isotope correction at $MS^2$ | No | Lipid Identification Software | MS-DIAL |
| $MS^1$ verified by standard | No | Data manipulation | Smoothing, Centroiding |
| $MS^2$ verified by standard | No | Nomenclature for intact lipid molecule | Yes |
| Background check at $MS^1$ | Yes | Nomenclature for fragment ions | No |
| Background check at $MS^2$ | No | | |

#### 20) Ceramide alpha-hydroxy fatty acid-phytospingosine (Cer_AP) / Lipid quantification

| | | | |
|---|---|---|---|
| Quantitative | Yes | Type I isotope correction | No |
| MS Level for quantification | $MS^1$ | Limit of quantification | No |
| Internal lipid standard(s) $MS^1$ | | Normalization to reference | No |

| Internal standard | Endogenous subclass |
|---|---|
| Cer 18:1;2O/15:0(d7) | Cer_AP subclass |

| | | | |
|---|---|---|---|
| Type of quantification | Internal standard amount | Lipid Quantification Software | MS-DIAL |
| Response correction | No | Batch correction | No |

#### 21) Ceramide alpha-hydroxy fatty acid-sphingosine (Cer_AS) / Lipid identification

| | | | |
|---|---|---|---|
| Lipid class | Ceramide alpha-hydroxy fatty acid-sphingosine (Cer_AS) | Did you presume assumptions for identification? | No |
| MS Level for identification | $MS^1$, $MS^2$ | Limit of detection | No |
| Identification level | Molecular species level | RT verified by standard | Yes |
| Isotope correction at $MS^1$ | No | Separation of isobaric/isomeric interferece confirmed | Yes |
| Fragments for identification | | Model for separation prediction | Yes |

| Fragment name |
|---|
| Neutral loss of CH4O2 |
| Neutral loss of H2O |
| Sphingosine -H2O fragment |
| Sphingosine -C2H7NO fragment |
| Oxidized fatty acyl -CH2O fragment |
| Oxidized fatty acyl -2H fragment |

| | | | |
|---|---|---|---|
| Isotope correction at $MS^2$ | No | Lipid Identification Software | MS-DIAL |
| $MS^1$ verified by standard | No | Data manipulation | Smoothing, Centroiding |
| $MS^2$ verified by standard | No | Nomenclature for intact lipid molecule | Yes |
| Background check at $MS^1$ | Yes | Nomenclature for fragment ions | No |
| Background check at $MS^2$ | No | | |

#### 21) Ceramide alpha-hydroxy fatty acid-sphingosine (Cer_AS) / Lipid quantification

| | | | |
|---|---|---|---|
| Quantitative | Yes | Type I isotope correction | No |
| MS Level for quantification | $MS^1$ | Limit of quantification | No |
| Internal lipid standard(s) $MS^1$ | | Normalization to reference | No |

| Internal standard | Endogenous subclass |
|---|---|
| Cer 18:1;2O/15:0(d7) | Cer_AS subclass |

| | | | |
|---|---|---|---|
| Type of quantification | Internal standard amount | Lipid Quantification Software | MS-DIAL |
| Response correction | No | Batch correction | No |

#### 22) Ceramide beta-hydroxy fatty acid-dihydrosphingosine (Cer_BDS) / Lipid identification

| | | | |
|---|---|---|---|
| Lipid class | Ceramide beta-hydroxy fatty acid-dihydrosphingosine (Cer_BDS) | Did you presume assumptions for identification? | No |
| MS Level for identification | $MS^1$, $MS^2$ | Limit of detection | No |
| Identification level | Molecular species level | RT verified by standard | Yes |
| Isotope correction at $MS^1$ | No | Separation of isobaric/isomeric interferece confirmed | Yes |
| Fragments for identification | | Model for separation prediction | Yes |
| Isotope correction at $MS^2$ | No | Lipid Identification Software | MS-DIAL |
| $MS^1$ verified by standard | No | Data manipulation | Smoothing, Centroiding |
| $MS^2$ verified by standard | No | Nomenclature for intact lipid molecule | Yes |
| Background check at $MS^1$ | Yes | Nomenclature for fragment ions | No |
| Background check at $MS^2$ | No | | |

Fragments for identification:

| Fragment name |
|---|
| Sphinganine |
| Sphinganine +C2H2O fragment |
| Sphinganine +C2H2O -CH4O fragment |
| Sphinganine -C2H7NO fragment |

#### 22) Ceramide beta-hydroxy fatty acid-dihydrosphingosine (Cer_BDS) / Lipid quantification

| | | | |
|---|---|---|---|
| Quantitative | Yes | Type I isotope correction | No |
| MS Level for quantification | $MS^1$ | Limit of quantification | No |
| Internal lipid standard(s) $MS^1$ | | Normalization to reference | No |
| Type of quantification | Internal standard amount | Lipid Quantification Software | MS-DIAL |
| Response correction | No | Batch correction | No |

Internal lipid standard(s) $MS^1$:

| Internal standard | Endogenous subclass |
|---|---|
| Cer 18:1;2O/15:0(d7) | Cer_BDS subclass |

#### 23) Ceramide beta-hydroxy fatty acid-sphingosine (Cer_BS) / Lipid identification

| | | | |
|---|---|---|---|
| Lipid class | Ceramide beta-hydroxy fatty acid-sphingosine (Cer_BS) | Did you presume assumptions for identification? | No |
| MS Level for identification | $MS^1$, $MS^2$ | Limit of detection | No |
| Identification level | Molecular species level | RT verified by standard | Yes |
| Isotope correction at $MS^1$ | No | Separation of isobaric/isomeric interferece confirmed | Yes |
| Fragments for identification | | Model for separation prediction | Yes |

| Fragment name |
|---|
| Sphingosine -C2H7NO fragment |
| Sphingosine +C2H2O fragment |
| Sphinganine +C2H2O -CH2O fragment |

| | | | |
|---|---|---|---|
| Isotope correction at $MS^2$ | No | Lipid Identification Software | MS-DIAL |
| $MS^1$ verified by standard | No | Data manipulation | Smoothing, Centroiding |
| $MS^2$ verified by standard | No | Nomenclature for intact lipid molecule | Yes |
| Background check at $MS^1$ | Yes | Nomenclature for fragment ions | No |
| Background check at $MS^2$ | No | | |

#### 23) Ceramide beta-hydroxy fatty acid-sphingosine (Cer_BS) / Lipid quantification

| | | | |
|---|---|---|---|
| Quantitative | Yes | Type I isotope correction | No |
| MS Level for quantification | $MS^1$ | Limit of quantification | No |
| Internal lipid standard(s) $MS^1$ | | Normalization to reference | No |

| Internal standard | Endogenous subclass |
|---|---|
| Cer 18:1;2O/15:0(d7) | Cer_BS subclass |

| | | | |
|---|---|---|---|
| Type of quantification | Internal standard amount | Lipid Quantification Software | MS-DIAL |
| Response correction | No | Batch correction | No |

#### 24) Ceramide Esterified beta-hydroxy fatty acid-dihydrosphingosine (Cer_EBDS) / Lipid identification

| | | | |
|---|---|---|---|
| Lipid class | Ceramide Esterified beta-hydroxy fatty acid-dihydrosphingosine (Cer_EBDS) | Did you presume assumptions for identification? | No |
| MS Level for identification | $MS^1$, $MS^2$ | Limit of detection | No |
| Identification level | Molecular species level | RT verified by standard | Yes |
| Isotope correction at $MS^1$ | No | Separation of isobaric/isomeric interferece confirmed | Yes |
| Fragments for identification | | Model for separation prediction | Yes |
| Isotope correction at $MS^2$ | No | Lipid Identification Software | MS-DIAL |
| $MS^1$ verified by standard | No | Data manipulation | Smoothing, Centroiding |
| $MS^2$ verified by standard | No | Nomenclature for intact lipid molecule | Yes |
| Background check at $MS^1$ | Yes | Nomenclature for fragment ions | No |
| Background check at $MS^2$ | No | | |

| Fragment name |
|---|
| Sphingosine +C2H2O fragment |
| Fatty acid fragment |
| Neutral loss of fatty acyl and H2O |

#### 24) Ceramide Esterified beta-hydroxy fatty acid-dihydrosphingosine (Cer_EBDS) / Lipid quantification

| | | | |
|---|---|---|---|
| Quantitative | Yes | Type I isotope correction | No |
| MS Level for quantification | $MS^1$ | Limit of quantification | No |
| Internal lipid standard(s) $MS^1$ | | Normalization to reference | No |
| Type of quantification | Internal standard amount | Lipid Quantification Software | MS-DIAL |
| Response correction | No | Batch correction | No |

| Internal standard | Endogenous subclass |
|---|---|
| Cer 18:1;2O/15:0(d7) | Cer_EBDS subclass |

#### 25) Ceramide Esterified omega-hydroxy fatty acid-dihydrosphingosine (Cer_EODS) / Lipid identification

| | | | |
|---|---|---|---|
| Lipid class | Ceramide Esterified omega-hydroxy fatty acid-dihydrosphingosine (Cer_EODS) | Did you presume assumptions for identification? | No |
| MS Level for identification | $MS^1$, $MS^2$ | Limit of detection | No |
| Identification level | Molecular species level | RT verified by standard | Yes |
| Isotope correction at $MS^1$ | No | Separation of isobaric/isomeric interferece confirmed | Yes |
| Fragments for identification | | Model for separation prediction | Yes |
| Isotope correction at $MS^2$ | No | Lipid Identification Software | MS-DIAL |
| $MS^1$ verified by standard | No | Data manipulation | Smoothing, Centroiding |
| $MS^2$ verified by standard | No | Nomenclature for intact lipid molecule | Yes |
| Background check at $MS^1$ | Yes | Nomenclature for fragment ions | No |
| Background check at $MS^2$ | No | | |

Fragments for identification:

| Fragment name |
|---|
| Fatty acid fragment |
| Neutral loss of fatty acyl |
| Acyl amide |

#### 25) Ceramide Esterified omega-hydroxy fatty acid-dihydrosphingosine (Cer_EODS) / Lipid quantification

| | | | |
|---|---|---|---|
| Quantitative | Yes | Type I isotope correction | No |
| MS Level for quantification | $MS^1$ | Limit of quantification | No |
| Internal lipid standard(s) $MS^1$ | | Normalization to reference | No |
| Type of quantification | Internal standard amount | Lipid Quantification Software | MS-DIAL |
| Response correction | No | Batch correction | No |

Internal lipid standard(s) $MS^1$:

| Internal standard | Endogenous subclass |
|---|---|
| Cer 18:1;2O/15:0(d7) | Cer_EODS subclass |

#### 26) Ceramide Esterified omega-hydroxy fatty acid-sphingosine (Cer_EOS) / Lipid identification

| | | | |
|---|---|---|---|
| Lipid class | Ceramide Esterified omega-hydroxy fatty acid-sphingosine (Cer_EOS) | Did you presume assumptions for identification? | No |
| MS Level for identification | $MS^1$, $MS^2$ | Limit of detection | No |
| Identification level | Molecular species level | RT verified by standard | Yes |
| Isotope correction at $MS^1$ | No | Separation of isobaric/isomeric interferece confirmed | Yes |
| Fragments for identification | | Model for separation prediction | Yes |
| Isotope correction at $MS^2$ | No | Lipid Identification Software | MS-DIAL |
| $MS^1$ verified by standard | No | Data manipulation | Smoothing, Centroiding |
| $MS^2$ verified by standard | No | Nomenclature for intact lipid molecule | Yes |
| Background check at $MS^1$ | Yes | Nomenclature for fragment ions | No |
| Background check at $MS^2$ | No | | |

Fragments for identification:

| Fragment name |
|---|
| Acyl amide |
| Fatty acid fragment |
| Neutral loss of fatty acyl |

#### 26) Ceramide Esterified omega-hydroxy fatty acid-sphingosine (Cer_EOS) / Lipid quantification

| | | | |
|---|---|---|---|
| Quantitative | Yes | Type I isotope correction | No |
| MS Level for quantification | $MS^1$ | Limit of quantification | No |
| Internal lipid standard(s) $MS^1$ | | Normalization to reference | No |
| Type of quantification | Internal standard amount | Lipid Quantification Software | MS-DIAL |
| Response correction | No | Batch correction | No |

Internal lipid standard(s) $MS^1$:

| Internal standard | Endogenous subclass |
|---|---|
| Cer 18:1;2O/15:0(d7) | Cer_EOS subclass |

#### 27) Ceramide non-hydroxyfatty acid-dihydrosphingosine (Cer_NDS) / Lipid identification

| | | | |
|---|---|---|---|
| Lipid class | Ceramide non-hydroxyfatty acid-dihydrosphingosine (Cer_NDS) | Did you presume assumptions for identification? | No |
| MS Level for identification | $MS^1$, $MS^2$ | Limit of detection | No |
| Identification level | Molecular species level | RT verified by standard | Yes |
| Isotope correction at $MS^1$ | No | Separation of isobaric/isomeric interferece confirmed | Yes |
| Fragments for identification | | Model for separation prediction | Yes |

| Fragment name |
|---|
| Neutral loss of CH4O |
| Neutral loss of CH4O2 |
| Sphinganine -C2H7NO fragment |
| Fatty acyl +C2H3N fragment |
| Fatty acyl -2H fragment |

| | | | |
|---|---|---|---|
| Isotope correction at $MS^2$ | No | Lipid Identification Software | MS-DIAL |
| $MS^1$ verified by standard | No | Data manipulation | Smoothing, Centroiding |
| $MS^2$ verified by standard | No | Nomenclature for intact lipid molecule | Yes |
| Background check at $MS^1$ | Yes | Nomenclature for fragment ions | No |
| Background check at $MS^2$ | No | | |

#### 27) Ceramide non-hydroxyfatty acid-dihydrosphingosine (Cer_NDS) / Lipid quantification

| | | | |
|---|---|---|---|
| Quantitative | Yes | Type I isotope correction | No |
| MS Level for quantification | $MS^1$ | Limit of quantification | No |
| Internal lipid standard(s) $MS^1$ | | Normalization to reference | No |

| Internal standard | Endogenous subclass |
|---|---|
| Cer 18:1;2O/15:0(d7) | Cer_NDS subclass |

| | | | |
|---|---|---|---|
| Type of quantification | Internal standard amount | Lipid Quantification Software | MS-DIAL |
| Response correction | No | Batch correction | No |

#### 28) Ceramide non-hydroxyfatty acid-phytospingosine (Cer_NP) / Lipid identification

| | | | |
|---|---|---|---|
| Lipid class | Ceramide non-hydroxyfatty acid-phytospingosine (Cer_NP) | Did you presume assumptions for identification? | No |
| MS Level for identification | $MS^1$, $MS^2$ | Limit of detection | No |
| Identification level | Molecular species level | RT verified by standard | Yes |
| Isotope correction at $MS^1$ | No | Separation of isobaric/isomeric interferece confirmed | Yes |
| Fragments for identification | | Model for separation prediction | Yes |

| Fragment name |
|---|
| Neutral loss of H2O |
| Neutral loss of 2H2O |
| Sphinganine -CH7NO fragment |
| Fatty acyl +C3H5NO fragment |
| Acyl amide |

| | | | |
|---|---|---|---|
| Isotope correction at $MS^2$ | No | Lipid Identification Software | MS-DIAL |
| $MS^1$ verified by standard | No | Data manipulation | Smoothing, Centroiding |
| $MS^2$ verified by standard | No | Nomenclature for intact lipid molecule | Yes |
| Background check at $MS^1$ | Yes | Nomenclature for fragment ions | No |
| Background check at $MS^2$ | No | | |

#### 28) Ceramide non-hydroxyfatty acid-phytospingosine (Cer_NP) / Lipid quantification

| | | | |
|---|---|---|---|
| Quantitative | Yes | Type I isotope correction | No |
| MS Level for quantification | $MS^1$ | Limit of quantification | No |
| Internal lipid standard(s) $MS^1$ | | Normalization to reference | No |

| Internal standard | Endogenous subclass |
|---|---|
| Cer 18:1;2O/15:0(d7) | Cer_NP subclass |

| | | | |
|---|---|---|---|
| Type of quantification | Internal standard amount | Lipid Quantification Software | MS-DIAL |
| Response correction | No | Batch correction | No |

#### 29) Ceramide non-hydroxyfatty acid-sphingosine (Cer_NS) / Lipid identification

| | | | |
|---|---|---|---|
| Lipid class | Ceramide non-hydroxyfatty acid-sphingosine (Cer_NS) | Did you presume assumptions for identification? | No |
| MS Level for identification | $MS^1$, $MS^2$ | Limit of detection | No |
| Identification level | Molecular species level | RT verified by standard | Yes |
| Isotope correction at $MS^1$ | No | Separation of isobaric/isomeric interferece confirmed | Yes |
| Fragments for identification | | Model for separation prediction | Yes |

| Fragment name |
|---|
| Neutral loss of H2O |
| Neutral loss of CH2O |
| Sphingosine -C2H7NO fragment |
| Fatty acyl +C2H3N fragment |
| Fatty acyl -2H fragment |

| | | | |
|---|---|---|---|
| Isotope correction at $MS^2$ | No | Lipid Identification Software | MS-DIAL |
| $MS^1$ verified by standard | Yes | Data manipulation | Smoothing, Centroiding |
| $MS^2$ verified by standard | Yes | Nomenclature for intact lipid molecule | Yes |
| Background check at $MS^1$ | Yes | Nomenclature for fragment ions | No |
| Background check at $MS^2$ | No | | |

#### 29) Ceramide non-hydroxyfatty acid-sphingosine (Cer_NS) / Lipid quantification

| | | | |
|---|---|---|---|
| Quantitative | Yes | Type I isotope correction | No |
| MS Level for quantification | $MS^1$ | Limit of quantification | No |
| Internal lipid standard(s) $MS^1$ | | Normalization to reference | No |

| Internal standard | Endogenous subclass |
|---|---|
| Cer 18:1;2O/15:0(d7) | Cer_NS subclass |

| | | | |
|---|---|---|---|
| Type of quantification | Internal standard amount | Lipid Quantification Software | MS-DIAL |
| Response correction | No | Batch correction | No |

#### 30) Ceramide phosphoethanolamine (PE_Cer) / Lipid identification

| | | | |
|---|---|---|---|
| Lipid class | Ceramide phosphoethanolamine (PE_Cer) | Did you presume assumptions for identification? | No |
| MS Level for identification | $MS^1$, $MS^2$ | Limit of detection | No |
| Identification level | Molecular species level | RT verified by standard | Yes |
| Isotope correction at $MS^1$ | No | Separation of isobaric/isomeric interferece confirmed | Yes |
| Fragments for identification | | Model for separation prediction | Yes |

| Fragment name |
|---|
| Characteristic fragment (C2H7NO4P-) |
| Neutral loss of fatty acyl |

| | | | |
|---|---|---|---|
| Isotope correction at $MS^2$ | No | Lipid Identification Software | MS-DIAL |
| $MS^1$ verified by standard | No | Data manipulation | Smoothing, Centroiding |
| $MS^2$ verified by standard | No | Nomenclature for intact lipid molecule | Yes |
| Background check at $MS^1$ | Yes | Nomenclature for fragment ions | No |
| Background check at $MS^2$ | No | | |

#### 30) Ceramide phosphoethanolamine (PE_Cer) / Lipid quantification

| | | | |
|---|---|---|---|
| Quantitative | Yes | Type I isotope correction | No |
| MS Level for quantification | $MS^1$ | Limit of quantification | No |
| Internal lipid standard(s) $MS^1$ | | Normalization to reference | No |

| Internal standard | Endogenous subclass |
|---|---|
| Cer 18:1;2O/15:0(d7) | PE_Cer subclass |

| | | | |
|---|---|---|---|
| Type of quantification | Internal standard amount | Lipid Quantification Software | MS-DIAL |
| Response correction | No | Batch correction | No |

#### 31) Ceramide phosphoinositol (PI_Cer) / Lipid identification

| | | | |
|---|---|---|---|
| Lipid class | Ceramide phosphoinositol (PI_Cer) | Did you presume assumptions for identification? | No |
| MS Level for identification | $MS^1$, $MS^2$ | Limit of detection | No |
| Identification level | Molecular species level | RT verified by standard | Yes |
| Isotope correction at $MS^1$ | No | Separation of isobaric/isomeric interferece confirmed | Yes |
| Fragments for identification | | Model for separation prediction | Yes |

| Fragment name |
|---|
| Phosphoinositol - H2O |
| Neutral loss of inositol |
| Neutral loss of fatty acyl |

| | | | |
|---|---|---|---|
| Isotope correction at $MS^2$ | No | Lipid Identification Software | MS-DIAL |
| $MS^1$ verified by standard | No | Data manipulation | Smoothing, Centroiding |
| $MS^2$ verified by standard | No | Nomenclature for intact lipid molecule | Yes |
| Background check at $MS^1$ | Yes | Nomenclature for fragment ions | No |
| Background check at $MS^2$ | No | | |

#### 31) Ceramide phosphoinositol (PI_Cer) / Lipid quantification

| | | | |
|---|---|---|---|
| Quantitative | Yes | Type I isotope correction | No |
| MS Level for quantification | $MS^1$ | Limit of quantification | No |
| Internal lipid standard(s) $MS^1$ | | Normalization to reference | No |

| Internal standard | Endogenous subclass |
|---|---|
| Cer 18:1;2O/15:0(d7) | PI_Cer subclass |

| | | | |
|---|---|---|---|
| Type of quantification | Internal standard amount | Lipid Quantification Software | MS-DIAL |
| Response correction | No | Batch correction | No |

#### 32) CAR / Lipid identification

| | | | |
|---|---|---|---|
| Lipid class | CAR | Did you presume assumptions for identification? | No |
| MS Level for identification | $MS^1$, $MS^2$ | Limit of detection | No |
| Identification level | Molecular species level | RT verified by standard | Yes |
| Isotope correction at $MS^1$ | No | Separation of isobaric/isomeric interferece confirmed | Yes |
| Fragments for identification | | Model for separation prediction | Yes |

| Fragment name |
|---|
| Neutral loss of fatty acyl |

| | | | |
|---|---|---|---|
| Isotope correction at $MS^2$ | No | Lipid Identification Software | MS-DIAL |
| $MS^1$ verified by standard | Yes | Data manipulation | Smoothing, Centroiding |
| $MS^2$ verified by standard | Yes | Nomenclature for intact lipid molecule | Yes |
| Background check at $MS^1$ | Yes | Nomenclature for fragment ions | No |
| Background check at $MS^2$ | No | | |

#### 32) CAR / Lipid quantification

| | | | |
|---|---|---|---|
| Quantitative | Yes | Type I isotope correction | No |
| MS Level for quantification | $MS^1$ | Limit of quantification | No |
| Internal lipid standard(s) $MS^1$ | | Normalization to reference | No |

| Internal standard | Endogenous subclass |
|---|---|
| CE 18:1(d7) | CE subclass |

| | | | |
|---|---|---|---|
| Type of quantification | Internal standard amount | Lipid Quantification Software | MS-DIAL |
| Response correction | No | Batch correction | No |

#### 33) Cholic acid (BileAcid) / Lipid identification

| | | | |
|---|---|---|---|
| Lipid class | Cholic acid (BileAcid) | Limit of detection | No |
| MS Level for identification | $MS^1$ | RT verified by standard | Yes |
| Identification level | Species level | Separation of isobaric/isomeric interferece confirmed | Yes |
| Isotope correction at $MS^1$ | No | Model for separation prediction | Yes |
| $MS^1$ verified by standard | No | Lipid Identification Software | MS-DIAL |
| Background check at $MS^1$ | Yes | Data manipulation | Smoothing, Centroiding |
| Did you presume assumptions for identification? | No | Nomenclature for intact lipid molecule | Yes |

#### 33) Cholic acid (BileAcid) / Lipid quantification

| | | | |
|---|---|---|---|
| Quantitative | Yes | Type I isotope correction | No |
| MS Level for quantification | $MS^1$ | Limit of quantification | No |
| Internal lipid standard(s) $MS^1$ | | Normalization to reference | No |

| Internal standard | Endogenous subclass |
|---|---|
| LPC 18:1(d7) | BileAcid subclass |

| | | | |
|---|---|---|---|
| Type of quantification | Internal standard amount | Lipid Quantification Software | MS-DIAL |
| Response correction | No | Batch correction | No |

#### 34) Cholic acid sulfate (BASulfate) / Lipid identification

| | | | |
|---|---|---|---|
| Lipid class | Cholic acid sulfate (BASulfate) | Did you presume assumptions for identification? | No |
| MS Level for identification | $MS^1$, $MS^2$ | Limit of detection | No |
| Identification level | Molecular species level | RT verified by standard | Yes |
| Isotope correction at $MS^1$ | No | Separation of isobaric/isomeric interferece confirmed | Yes |
| Fragments for identification | | Model for separation prediction | Yes |

| Fragment name |
|---|
| Hydrogensulfate |

| | | | |
|---|---|---|---|
| Isotope correction at $MS^2$ | No | Lipid Identification Software | MS-DIAL |
| $MS^1$ verified by standard | No | Data manipulation | Smoothing, Centroiding |
| $MS^2$ verified by standard | No | Nomenclature for intact lipid molecule | Yes |
| Background check at $MS^1$ | Yes | Nomenclature for fragment ions | No |
| Background check at $MS^2$ | No | | |

#### 34) Cholic acid sulfate (BASulfate) / Lipid quantification

| | | | |
|---|---|---|---|
| Quantitative | Yes | Type I isotope correction | No |
| MS Level for quantification | $MS^1$ | Limit of quantification | No |
| Internal lipid standard(s) $MS^1$ | | Normalization to reference | No |

| Internal standard | Endogenous subclass |
|---|---|
| LPC 18:1(d7) | BASulfate subclass |

| | | | |
|---|---|---|---|
| Type of quantification | Internal standard amount | Lipid Quantification Software | MS-DIAL |
| Response correction | No | Batch correction | No |

#### 35) Coenzyme Q (CoQ) / Lipid identification

| | | | |
|---|---|---|---|
| Lipid class | Coenzyme Q (CoQ) | Did you presume assumptions for identification? | No |
| MS Level for identification | $MS^1$, $MS^2$ | Limit of detection | No |
| Identification level | Molecular species level | RT verified by standard | Yes |
| Isotope correction at $MS^1$ | No | Separation of isobaric/isomeric interferece confirmed | Yes |
| Fragments for identification | | Model for separation prediction | Yes |

| Fragment name |
|---|
| Characteristic fragment (C10H13O4+) |

| | | | |
|---|---|---|---|
| Isotope correction at $MS^2$ | No | Lipid Identification Software | MS-DIAL |
| $MS^1$ verified by standard | No | Data manipulation | Smoothing, Centroiding |
| $MS^2$ verified by standard | No | Nomenclature for intact lipid molecule | Yes |
| Background check at $MS^1$ | Yes | Nomenclature for fragment ions | No |
| Background check at $MS^2$ | No | | |

#### 35) Coenzyme Q (CoQ) / Lipid quantification

| | | | |
|---|---|---|---|
| Quantitative | Yes | Type I isotope correction | No |
| MS Level for quantification | $MS^1$ | Limit of quantification | No |
| Internal lipid standard(s) $MS^1$ | | Normalization to reference | No |

| Internal standard | Endogenous subclass |
|---|---|
| LPC 18:1(d7) | CoQ subclass |

| | | | |
|---|---|---|---|
| Type of quantification | Internal standard amount | Lipid Quantification Software | MS-DIAL |
| Response correction | No | Batch correction | No |

#### 36) Dehydroergosterol ester (DEGSE) / Lipid identification

| | | | |
|---|---|---|---|
| Lipid class | Dehydroergosterol ester (DEGSE) | Did you presume assumptions for identification? | No |
| MS Level for identification | $MS^1$, $MS^2$ | Limit of detection | No |
| Identification level | Molecular species level | RT verified by standard | Yes |
| Isotope correction at $MS^1$ | No | Separation of isobaric/isomeric interferece confirmed | Yes |
| Fragments for identification | | Model for separation prediction | Yes |

| Fragment name |
|---|
| Neutral loss of fatty acyl |

| | | | |
|---|---|---|---|
| Isotope correction at $MS^2$ | No | Lipid Identification Software | MS-DIAL |
| $MS^1$ verified by standard | No | Data manipulation | Smoothing, Centroiding |
| $MS^2$ verified by standard | No | Nomenclature for intact lipid molecule | Yes |
| Background check at $MS^1$ | Yes | Nomenclature for fragment ions | No |
| Background check at $MS^2$ | No | | |

#### 36) Dehydroergosterol ester (DEGSE) / Lipid quantification

| | | | |
|---|---|---|---|
| Quantitative | Yes | Type I isotope correction | No |
| MS Level for quantification | $MS^1$ | Limit of quantification | No |
| Internal lipid standard(s) $MS^1$ | | Normalization to reference | No |

| Internal standard | Endogenous subclass |
|---|---|
| CE 18:1(d7) | DEGSE subclass |

| | | | |
|---|---|---|---|
| Type of quantification | Internal standard amount | Lipid Quantification Software | MS-DIAL |
| Response correction | No | Batch correction | No |

#### 37) Desmosterol ester (DSMSE) / Lipid identification

| | | | |
|---|---|---|---|
| Lipid class | Desmosterol ester (DSMSE) | Did you presume assumptions for identification? | No |
| MS Level for identification | $MS^1$, $MS^2$ | Limit of detection | No |
| Identification level | Molecular species level | RT verified by standard | Yes |
| Isotope correction at $MS^1$ | No | Separation of isobaric/isomeric interferece confirmed | Yes |
| Fragments for identification | Fragment name: Neutral loss of fatty acyl | Model for separation prediction | Yes |
| Isotope correction at $MS^2$ | No | Lipid Identification Software | MS-DIAL |
| $MS^1$ verified by standard | No | Data manipulation | Smoothing, Centroiding |
| $MS^2$ verified by standard | No | Nomenclature for intact lipid molecule | Yes |
| Background check at $MS^1$ | Yes | Nomenclature for fragment ions | No |
| Background check at $MS^2$ | No | | |

#### 37) Desmosterol ester (DSMSE) / Lipid quantification

| | | | |
|---|---|---|---|
| Quantitative | Yes | Type I isotope correction | No |
| MS Level for quantification | $MS^1$ | Limit of quantification | No |
| Internal lipid standard(s) $MS^1$ | Internal standard: CE 18:1(d7); Endogenous subclass: DSMSE subclass | Normalization to reference | No |
| Type of quantification | Internal standard amount | Lipid Quantification Software | MS-DIAL |
| Response correction | No | Batch correction | No |

#### 38) Diacylglyceryl glucuronide (DGGA) / Lipid identification

| | | | |
|---|---|---|---|
| Lipid class | Diacylglyceryl glucuronide (DGGA) | Did you presume assumptions for identification? | No |
| MS Level for identification | $MS^1$, $MS^2$ | Limit of detection | No |
| Identification level | Molecular species level | RT verified by standard | Yes |
| Isotope correction at $MS^1$ | No | Separation of isobaric/isomeric interferece confirmed | Yes |
| Fragments for identification | Fragment name: Fatty acid fragment | Model for separation prediction | Yes |
| Isotope correction at $MS^2$ | No | Lipid Identification Software | MS-DIAL |
| $MS^1$ verified by standard | No | Data manipulation | Smoothing, Centroiding |
| $MS^2$ verified by standard | No | Nomenclature for intact lipid molecule | Yes |
| Background check at $MS^1$ | Yes | Nomenclature for fragment ions | No |
| Background check at $MS^2$ | No | | |

#### 38) Diacylglyceryl glucuronide (DGGA) / Lipid quantification

| | | | |
|---|---|---|---|
| Quantitative | Yes | Type I isotope correction | No |
| MS Level for quantification | $MS^1$ | Limit of quantification | No |
| Internal lipid standard(s) $MS^1$ | | Normalization to reference | No |

| Internal standard | Endogenous subclass |
|---|---|
| LPC 18:1(d7) | DGGA subclass |

| | | | |
|---|---|---|---|
| Type of quantification | Internal standard amount | Lipid Quantification Software | MS-DIAL |
| Response correction | No | Batch correction | No |

#### 39) Diacylglyceryl trimethylhomoserine (DGTS) / Lipid identification

| | | | |
|---|---|---|---|
| Lipid class | Diacylglyceryl trimethylhomoserine (DGTS) | Did you presume assumptions for identification? | No |
| MS Level for identification | $MS^1$, $MS^2$ | Limit of detection | No |
| Identification level | Molecular species level | RT verified by standard | Yes |
| Isotope correction at $MS^1$ | No | Separation of isobaric/isomeric interferece confirmed | Yes |
| Fragments for identification | | Model for separation prediction | Yes |

| Fragment name |
|---|
| Characteristic fragment (C10H22NO5+) |
| Characteristic fragment (C7H14NO2+) |
| Neutral loss of fatty acyl |
| Neutral loss of fatty acyl and H2O |

| | | | |
|---|---|---|---|
| Isotope correction at $MS^2$ | No | Lipid Identification Software | MS-DIAL |
| $MS^1$ verified by standard | No | Data manipulation | Smoothing, Centroiding |
| $MS^2$ verified by standard | No | Nomenclature for intact lipid molecule | Yes |
| Background check at $MS^1$ | Yes | Nomenclature for fragment ions | No |
| Background check at $MS^2$ | No | | |

#### 39) Diacylglyceryl trimethylhomoserine (DGTS) / Lipid quantification

| | | | |
|---|---|---|---|
| Quantitative | Yes | Type I isotope correction | No |
| MS Level for quantification | $MS^1$ | Limit of quantification | No |
| Internal lipid standard(s) $MS^1$ | | Normalization to reference | No |

| Internal standard | Endogenous subclass |
|---|---|
| LPC 18:1(d7) | DGTS subclass |

| | | | |
|---|---|---|---|
| Type of quantification | Internal standard amount | Lipid Quantification Software | MS-DIAL |
| Response correction | No | Batch correction | No |

#### 40) Diacylglyceryl-3-O-carboxyhydroxymethylcholine (DGCC) / Lipid identification

| | | | |
|---|---|---|---|
| Lipid class | Diacylglyceryl-3-O-carboxyhydroxymethylcholine (DGCC) | Did you presume assumptions for identification? | No |
| MS Level for identification | $MS^1$, $MS^2$ | Limit of detection | No |
| Identification level | Molecular species level | RT verified by standard | Yes |
| Isotope correction at $MS^1$ | No | Separation of isobaric/isomeric interferece confirmed | Yes |
| Fragments for identification | | Model for separation prediction | Yes |

| Fragment name |
|---|
| Characteristic fragment (C6H14NO2+) |
| Neutral loss of fatty acyl |
| Neutral loss of fatty acyl and H2O |

| | | | |
|---|---|---|---|
| Isotope correction at $MS^2$ | No | Lipid Identification Software | MS-DIAL |
| $MS^1$ verified by standard | No | Data manipulation | Smoothing, Centroiding |
| $MS^2$ verified by standard | No | Nomenclature for intact lipid molecule | Yes |
| Background check at $MS^1$ | Yes | Nomenclature for fragment ions | No |
| Background check at $MS^2$ | No | | |

#### 40) Diacylglyceryl-3-O-carboxyhydroxymethylcholine (DGCC) / Lipid quantification

| | | | |
|---|---|---|---|
| Quantitative | Yes | Type I isotope correction | No |
| MS Level for quantification | $MS^1$ | Limit of quantification | No |
| Internal lipid standard(s) $MS^1$ | | Normalization to reference | No |

| Internal standard | Endogenous subclass |
|---|---|
| LPC 18:1(d7) | DGCC subclass |

| | | | |
|---|---|---|---|
| Type of quantification | Internal standard amount | Lipid Quantification Software | MS-DIAL |
| Response correction | No | Batch correction | No |

#### 41) Digalactosylmonoacylglycerol (DGMG) / Lipid identification

| | | | |
|---|---|---|---|
| Lipid class | Digalactosylmonoacylglycerol (DGMG) | Did you presume assumptions for identification? | No |
| MS Level for identification | $MS^1$, $MS^2$ | Limit of detection | No |
| Identification level | Molecular species level | RT verified by standard | Yes |
| Isotope correction at $MS^1$ | No | Separation of isobaric/isomeric interferece confirmed | Yes |
| Fragments for identification | | Model for separation prediction | Yes |

| Fragment name |
|---|
| Fatty acid fragment |

| | | | |
|---|---|---|---|
| Isotope correction at $MS^2$ | No | Lipid Identification Software | MS-DIAL |
| $MS^1$ verified by standard | No | Data manipulation | Smoothing, Centroiding |
| $MS^2$ verified by standard | No | Nomenclature for intact lipid molecule | Yes |
| Background check at $MS^1$ | Yes | Nomenclature for fragment ions | No |
| Background check at $MS^2$ | No | | |

#### 41) Digalactosylmonoacylglycerol (DGMG) / Lipid quantification

| | | | |
|---|---|---|---|
| Quantitative | Yes | Type I isotope correction | No |
| MS Level for quantification | $MS^1$ | Limit of quantification | No |
| Internal lipid standard(s) $MS^1$ | | Normalization to reference | No |

| Internal standard | Endogenous subclass |
|---|---|
| LPC 18:1(d7) | DGMG subclass |

| | | | |
|---|---|---|---|
| Type of quantification | Internal standard amount | Lipid Quantification Software | MS-DIAL |
| Response correction | No | Batch correction | No |

#### 42) Hex2Cer / Lipid identification

| | | | |
|---|---|---|---|
| Lipid class | Hex2Cer | Did you presume assumptions for identification? | No |
| MS Level for identification | $MS^1$, $MS^2$ | Limit of detection | No |
| Identification level | Molecular species level | RT verified by standard | Yes |
| Isotope correction at $MS^1$ | No | Separation of isobaric/isomeric interferece confirmed | Yes |
| Fragments for identification | | Model for separation prediction | Yes |

| Fragment name |
|---|
| Neutral loss of hexose |
| Neutral loss of 2hexose |
| Sphingosine -H2O fragment |
| Sphingosine -2H2O fragment |
| Sphingosine -CH4O2 fragment |

| | | | |
|---|---|---|---|
| Isotope correction at $MS^2$ | No | Lipid Identification Software | MS-DIAL |
| $MS^1$ verified by standard | No | Data manipulation | Smoothing, Centroiding |
| $MS^2$ verified by standard | No | Nomenclature for intact lipid molecule | Yes |
| Background check at $MS^1$ | Yes | Nomenclature for fragment ions | No |
| Background check at $MS^2$ | No | | |

#### 42) Hex2Cer / Lipid quantification

| | | | |
|---|---|---|---|
| Quantitative | Yes | Type I isotope correction | No |
| MS Level for quantification | $MS^1$ | Limit of quantification | No |
| Internal lipid standard(s) $MS^1$ | | Normalization to reference | No |

| Internal standard | Endogenous subclass |
|---|---|
| Cer 18:1;2O/15:0(d7) | Hex2Cer subclass |

| | | | |
|---|---|---|---|
| Type of quantification | Internal standard amount | Lipid Quantification Software | MS-DIAL |
| Response correction | No | Batch correction | No |

#### 43) Dilysocardiolipin (DLCL) / Lipid identification

| | | | |
|---|---|---|---|
| Lipid class | Dilysocardiolipin (DLCL) | Did you presume assumptions for identification? | No |
| MS Level for identification | $MS^1$, $MS^2$ | Limit of detection | No |
| Identification level | Molecular species level | RT verified by standard | Yes |
| Isotope correction at $MS^1$ | No | Separation of isobaric/isomeric interferece confirmed | Yes |
| Fragments for identification | | Model for separation prediction | Yes |

| Fragment name |
|---|
| Phosphoglycerol -H2O fragment |
| lysophosphatidic acid |

| | | | |
|---|---|---|---|
| Isotope correction at $MS^2$ | No | Lipid Identification Software | MS-DIAL |
| $MS^1$ verified by standard | No | Data manipulation | Smoothing, Centroiding |
| $MS^2$ verified by standard | No | Nomenclature for intact lipid molecule | Yes |
| Background check at $MS^1$ | Yes | Nomenclature for fragment ions | No |
| Background check at $MS^2$ | No | | |

#### 43) Dilysocardiolipin (DLCL) / Lipid quantification

| | | | |
|---|---|---|---|
| Quantitative | Yes | Type I isotope correction | No |
| MS Level for quantification | $MS^1$ | Limit of quantification | No |
| Internal lipid standard(s) $MS^1$ | | Normalization to reference | No |

| Internal standard | Endogenous subclass |
|---|---|
| PG 15:0_18:1(d7) | DLCL subclass |

| | | | |
|---|---|---|---|
| Type of quantification | Internal standard amount | Lipid Quantification Software | MS-DIAL |
| Response correction | No | Batch correction | No |

#### 44) Ergosterol ester (EGSE) / Lipid identification

| | | | |
|---|---|---|---|
| Lipid class | Ergosterol ester (EGSE) | Did you presume assumptions for identification? | No |
| MS Level for identification | $MS^1$, $MS^2$ | Limit of detection | No |
| Identification level | Molecular species level | RT verified by standard | Yes |
| Isotope correction at $MS^1$ | No | Separation of isobaric/isomeric interferece confirmed | Yes |
| Fragments for identification | | Model for separation prediction | Yes |
| Isotope correction at $MS^2$ | No | Lipid Identification Software | MS-DIAL |
| $MS^1$ verified by standard | No | Data manipulation | Smoothing, Centroiding |
| $MS^2$ verified by standard | No | Nomenclature for intact lipid molecule | Yes |
| Background check at $MS^1$ | Yes | Nomenclature for fragment ions | No |
| Background check at $MS^2$ | No | | |

Fragments for identification:

| Fragment name |
|---|
| Neutral loss of fatty acyl |

#### 44) Ergosterol ester (EGSE) / Lipid quantification

| | | | |
|---|---|---|---|
| Quantitative | Yes | Type I isotope correction | No |
| MS Level for quantification | $MS^1$ | Limit of quantification | No |
| Internal lipid standard(s) $MS^1$ | | Normalization to reference | No |
| Type of quantification | Internal standard amount | Lipid Quantification Software | MS-DIAL |
| Response correction | No | Batch correction | No |

Internal lipid standard(s) $MS^1$:

| Internal standard | Endogenous subclass |
|---|---|
| CE 18:1(d7) | EGSE subclass |

#### 45) Esterified ketodeoxycholic acid (KDCAE) / Lipid identification

| | | | |
|---|---|---|---|
| Lipid class | Esterified ketodeoxycholic acid (KDCAE) | Did you presume assumptions for identification? | No |
| MS Level for identification | $MS^1$, $MS^2$ | Limit of detection | No |
| Identification level | Molecular species level | RT verified by standard | Yes |
| Isotope correction at $MS^1$ | No | Separation of isobaric/isomeric interferece confirmed | Yes |
| Fragments for identification | | Model for separation prediction | Yes |
| Isotope correction at $MS^2$ | No | Lipid Identification Software | MS-DIAL |
| $MS^1$ verified by standard | No | Data manipulation | Smoothing, Centroiding |
| $MS^2$ verified by standard | No | Nomenclature for intact lipid molecule | Yes |
| Background check at $MS^1$ | Yes | Nomenclature for fragment ions | No |
| Background check at $MS^2$ | No | | |

Fragments for identification:

| Fragment name |
|---|
| Neutral loss of fatty acyl |

#### 45) Esterified ketodeoxycholic acid (KDCAE) / Lipid quantification

| | | | |
|---|---|---|---|
| Quantitative | Yes | Type I isotope correction | No |
| MS Level for quantification | $MS^1$ | Limit of quantification | No |
| Internal lipid standard(s) $MS^1$ | | Normalization to reference | No |

| Internal standard | Endogenous subclass |
|---|---|
| CE 18:1(d7) | KDCAE subclass |

| | | | |
|---|---|---|---|
| Type of quantification | Internal standard amount | Lipid Quantification Software | MS-DIAL |
| Response correction | No | Batch correction | No |

#### 46) Esterified taurodeoxycholic Acid (TDCAE) / Lipid identification

| | | | |
|---|---|---|---|
| Lipid class | Esterified taurodeoxycholic Acid (TDCAE) | Did you presume assumptions for identification? | No |
| MS Level for identification | $MS^1$, $MS^2$ | Limit of detection | No |
| Identification level | Molecular species level | RT verified by standard | Yes |
| Isotope correction at $MS^1$ | No | Separation of isobaric/isomeric interferece confirmed | Yes |
| Fragments for identification | | Model for separation prediction | Yes |

| Fragment name |
|---|
| Neutral loss of fatty acyl |

| | | | |
|---|---|---|---|
| Isotope correction at $MS^2$ | No | Lipid Identification Software | MS-DIAL |
| $MS^1$ verified by standard | No | Data manipulation | Smoothing, Centroiding |
| $MS^2$ verified by standard | No | Nomenclature for intact lipid molecule | Yes |
| Background check at $MS^1$ | Yes | Nomenclature for fragment ions | No |
| Background check at $MS^2$ | No | | |

#### 46) Esterified taurodeoxycholic Acid (TDCAE) / Lipid quantification

| | | | |
|---|---|---|---|
| Quantitative | Yes | Type I isotope correction | No |
| MS Level for quantification | $MS^1$ | Limit of quantification | No |
| Internal lipid standard(s) $MS^1$ | | Normalization to reference | No |

| Internal standard | Endogenous subclass |
|---|---|
| CE 18:1(d7) | TDCAE subclass |

| | | | |
|---|---|---|---|
| Type of quantification | Internal standard amount | Lipid Quantification Software | MS-DIAL |
| Response correction | No | Batch correction | No |

#### 47) Esterified deoxycholic acid (DCAE) / Lipid identification

| | | | |
|---|---|---|---|
| Lipid class | Esterified deoxycholic acid (DCAE) | Did you presume assumptions for identification? | No |
| MS Level for identification | $MS^1$, $MS^2$ | Limit of detection | No |
| Identification level | Molecular species level | RT verified by standard | Yes |
| Isotope correction at $MS^1$ | No | Separation of isobaric/isomeric interferece confirmed | Yes |
| Fragments for identification | | Model for separation prediction | Yes |
| Isotope correction at $MS^2$ | No | Lipid Identification Software | MS-DIAL |
| $MS^1$ verified by standard | No | Data manipulation | Smoothing, Centroiding |
| $MS^2$ verified by standard | No | Nomenclature for intact lipid molecule | Yes |
| Background check at $MS^1$ | Yes | Nomenclature for fragment ions | No |
| Background check at $MS^2$ | No | | |

| Fragment name |
|---|
| Neutral loss of fatty acyl |

#### 47) Esterified deoxycholic acid (DCAE) / Lipid quantification

| | | | |
|---|---|---|---|
| Quantitative | Yes | Type I isotope correction | No |
| MS Level for quantification | $MS^1$ | Limit of quantification | No |
| Internal lipid standard(s) $MS^1$ | | Normalization to reference | No |
| Type of quantification | Internal standard amount | Lipid Quantification Software | MS-DIAL |
| Response correction | No | Batch correction | No |

| Internal standard | Endogenous subclass |
|---|---|
| CE 18:1(d7) | DCAE subclass |

#### 48) Esterified ketolithocholic acid (KLCAE) / Lipid identification

| | | | |
|---|---|---|---|
| Lipid class | Esterified ketolithocholic acid (KLCAE) | Did you presume assumptions for identification? | No |
| MS Level for identification | $MS^1$, $MS^2$ | Limit of detection | No |
| Identification level | Molecular species level | RT verified by standard | Yes |
| Isotope correction at $MS^1$ | No | Separation of isobaric/isomeric interferece confirmed | Yes |
| Fragments for identification | | Model for separation prediction | Yes |
| Isotope correction at $MS^2$ | No | Lipid Identification Software | MS-DIAL |
| $MS^1$ verified by standard | No | Data manipulation | Smoothing, Centroiding |
| $MS^2$ verified by standard | No | Nomenclature for intact lipid molecule | Yes |
| Background check at $MS^1$ | Yes | Nomenclature for fragment ions | No |
| Background check at $MS^2$ | No | | |

| Fragment name |
|---|
| Neutral loss of fatty acyl |

#### 48) Esterified ketolithocholic acid (KLCAE) / Lipid quantification

| | | | |
|---|---|---|---|
| Quantitative | Yes | Type I isotope correction | No |
| MS Level for quantification | $MS^1$ | Limit of quantification | No |
| Internal lipid standard(s) $MS^1$ | | Normalization to reference | No |

| Internal standard | Endogenous subclass |
|---|---|
| CE 18:1(d7) | KLCAE subclass |

| | | | |
|---|---|---|---|
| Type of quantification | Internal standard amount | Lipid Quantification Software | MS-DIAL |
| Response correction | No | Batch correction | No |

#### 49) Esterified lithocholic acid (LCAE) / Lipid identification

| | | | |
|---|---|---|---|
| Lipid class | Esterified lithocholic acid (LCAE) | Did you presume assumptions for identification? | No |
| MS Level for identification | $MS^1$, $MS^2$ | Limit of detection | No |
| Identification level | Molecular species level | RT verified by standard | Yes |
| Isotope correction at $MS^1$ | No | Separation of isobaric/isomeric interferece confirmed | Yes |
| Fragments for identification | | Model for separation prediction | Yes |

| Fragment name |
|---|
| Neutral loss of fatty acyl |

| | | | |
|---|---|---|---|
| Isotope correction at $MS^2$ | No | Lipid Identification Software | MS-DIAL |
| $MS^1$ verified by standard | No | Data manipulation | Smoothing, Centroiding |
| $MS^2$ verified by standard | No | Nomenclature for intact lipid molecule | Yes |
| Background check at $MS^1$ | Yes | Nomenclature for fragment ions | No |
| Background check at $MS^2$ | No | | |

#### 49) Esterified lithocholic acid (LCAE) / Lipid quantification

| | | | |
|---|---|---|---|
| Quantitative | Yes | Type I isotope correction | No |
| MS Level for quantification | $MS^1$ | Limit of quantification | No |
| Internal lipid standard(s) $MS^1$ | | Normalization to reference | No |

| Internal standard | Endogenous subclass |
|---|---|
| CE 18:1(d7) | LCAE subclass |

| | | | |
|---|---|---|---|
| Type of quantification | Internal standard amount | Lipid Quantification Software | MS-DIAL |
| Response correction | No | Batch correction | No |

#### 50) Ether-linked digalactosyldiacylglycerol (EtherDGDG) / Lipid identification

| | | | |
|---|---|---|---|
| Lipid class | Ether-linked digalactosyldiacylglycerol (EtherDGDG) | Did you presume assumptions for identification? | No |
| MS Level for identification | $MS^1$, $MS^2$ | Limit of detection | No |
| Identification level | Molecular species level | RT verified by standard | Yes |
| Isotope correction at $MS^1$ | No | Separation of isobaric/isomeric interferece confirmed | Yes |
| Fragments for identification | | Model for separation prediction | Yes |
| Isotope correction at $MS^2$ | No | Lipid Identification Software | MS-DIAL |
| $MS^1$ verified by standard | No | Data manipulation | Smoothing, Centroiding |
| $MS^2$ verified by standard | No | Nomenclature for intact lipid molecule | Yes |
| Background check at $MS^1$ | Yes | Nomenclature for fragment ions | No |
| Background check at $MS^2$ | No | | |

| Fragment name |
|---|
| Neutral loss of fatty acyl |
| Fatty acid fragment |

#### 50) Ether-linked digalactosyldiacylglycerol (EtherDGDG) / Lipid quantification

| | | | |
|---|---|---|---|
| Quantitative | Yes | Type I isotope correction | No |
| MS Level for quantification | $MS^1$ | Limit of quantification | No |
| Internal lipid standard(s) $MS^1$ | | Normalization to reference | No |
| Type of quantification | Internal standard amount | Lipid Quantification Software | MS-DIAL |
| Response correction | No | Batch correction | No |

| Internal standard | Endogenous subclass |
|---|---|
| LPC 18:1(d7) | EtherDGDG subclass |

#### 51) LPC O / Lipid identification

| | | | |
|---|---|---|---|
| Lipid class | LPC O | Did you presume assumptions for identification? | No |
| MS Level for identification | $MS^1$, $MS^2$ | Limit of detection | No |
| Identification level | Molecular species level | RT verified by standard | Yes |
| Isotope correction at $MS^1$ | No | Separation of isobaric/isomeric interferece confirmed | Yes |
| Fragments for identification | | Model for separation prediction | Yes |

| Fragment name |
|---|
| Characteristic fragments (C5H14NO+) |
| Characteristic fragments (C2H6O4P+) |
| Characteristic fragments (C5H15NO4P+) |

| | | | |
|---|---|---|---|
| Isotope correction at $MS^2$ | No | Lipid Identification Software | MS-DIAL |
| $MS^1$ verified by standard | No | Data manipulation | Smoothing, Centroiding |
| $MS^2$ verified by standard | No | Nomenclature for intact lipid molecule | Yes |
| Background check at $MS^1$ | Yes | Nomenclature for fragment ions | No |
| Background check at $MS^2$ | No | | |

#### 51) LPC O / Lipid quantification

| | | | |
|---|---|---|---|
| Quantitative | Yes | Type I isotope correction | No |
| MS Level for quantification | $MS^1$ | Limit of quantification | No |
| Internal lipid standard(s) $MS^1$ | | Normalization to reference | No |

| Internal standard | Endogenous subclass |
|---|---|
| LPC 18:1(d7) | EtherLPC subclass |

| | | | |
|---|---|---|---|
| Type of quantification | Internal standard amount | Lipid Quantification Software | MS-DIAL |
| Response correction | No | Batch correction | No |

#### 52) LPE O / Lipid identification

| | | | |
|---|---|---|---|
| Lipid class | LPE O | Did you presume assumptions for identification? | No |
| MS Level for identification | $MS^1$, $MS^2$ | Limit of detection | No |
| Identification level | Molecular species level | RT verified by standard | Yes |
| Isotope correction at $MS^1$ | No | Separation of isobaric/isomeric interferece confirmed | Yes |
| Fragments for identification | | Model for separation prediction | Yes |

| Fragment name |
|---|
| Neutral loss of C3H8NO4P |
| Neutral loss of C3H10NO5P |

| | | | |
|---|---|---|---|
| Isotope correction at $MS^2$ | No | Lipid Identification Software | MS-DIAL |
| $MS^1$ verified by standard | No | Data manipulation | Smoothing, Centroiding |
| $MS^2$ verified by standard | No | Nomenclature for intact lipid molecule | Yes |
| Background check at $MS^1$ | Yes | Nomenclature for fragment ions | No |
| Background check at $MS^2$ | No | | |

#### 52) LPE O / Lipid quantification

| | | | |
|---|---|---|---|
| Quantitative | Yes | Type I isotope correction | No |
| MS Level for quantification | $MS^1$ | Limit of quantification | No |
| Internal lipid standard(s) $MS^1$ | | Normalization to reference | No |

| Internal standard | Endogenous subclass |
|---|---|
| LPE 18:1(d7) | EtherLPE subclass |

| | | | |
|---|---|---|---|
| Type of quantification | Internal standard amount | Lipid Quantification Software | MS-DIAL |
| Response correction | No | Batch correction | No |

#### 53) Ether-linked lysophosphatidylglycerol (EtherLPG) / Lipid identification

| | | | |
|---|---|---|---|
| Lipid class | Ether-linked lysophosphatidylglycerol (EtherLPG) | Did you presume assumptions for identification? | No |
| MS Level for identification | $MS^1$, $MS^2$ | Limit of detection | No |
| Identification level | Molecular species level | RT verified by standard | Yes |
| Isotope correction at $MS^1$ | No | Separation of isobaric/isomeric interferece confirmed | Yes |
| Fragments for identification | | Model for separation prediction | Yes |

| Fragment name |
|---|
| Phosphoglycerol -H2O fragment |
| Alkyl Ether fragment |

| | | | |
|---|---|---|---|
| Isotope correction at $MS^2$ | No | Lipid Identification Software | MS-DIAL |
| $MS^1$ verified by standard | No | Data manipulation | Smoothing, Centroiding |
| $MS^2$ verified by standard | No | Nomenclature for intact lipid molecule | Yes |
| Background check at $MS^1$ | Yes | Nomenclature for fragment ions | No |
| Background check at $MS^2$ | No | | |

#### 53) Ether-linked lysophosphatidylglycerol (EtherLPG) / Lipid quantification

| | | | |
|---|---|---|---|
| Quantitative | Yes | Type I isotope correction | No |
| MS Level for quantification | $MS^1$ | Limit of quantification | No |
| Internal lipid standard(s) $MS^1$ | | Normalization to reference | No |

| Internal standard | Endogenous subclass |
|---|---|
| PG 15:0_18:1(d7) | EtherLPG subclass |

| | | | |
|---|---|---|---|
| Type of quantification | Internal standard amount | Lipid Quantification Software | MS-DIAL |
| Response correction | No | Batch correction | No |

#### 54) Ether-linked monogalactosyldiacylglycerol (EtherMGDG) / Lipid identification

| | | | |
|---|---|---|---|
| Lipid class | Ether-linked monogalactosyldiacylglycerol (EtherMGDG) | Did you presume assumptions for identification? | No |
| MS Level for identification | $MS^1$, $MS^2$ | Limit of detection | No |
| Identification level | Molecular species level | RT verified by standard | Yes |
| Isotope correction at $MS^1$ | No | Separation of isobaric/isomeric interferece confirmed | Yes |
| Fragments for identification | | Model for separation prediction | Yes |
| Isotope correction at $MS^2$ | No | Lipid Identification Software | MS-DIAL |
| $MS^1$ verified by standard | No | Data manipulation | Smoothing, Centroiding |
| $MS^2$ verified by standard | No | Nomenclature for intact lipid molecule | Yes |
| Background check at $MS^1$ | Yes | Nomenclature for fragment ions | No |
| Background check at $MS^2$ | No | | |

| Fragment name |
|---|
| Neutral loss of fatty acyl |
| Fatty acid fragment |

#### 54) Ether-linked monogalactosyldiacylglycerol (EtherMGDG) / Lipid quantification

| | | | |
|---|---|---|---|
| Quantitative | Yes | Type I isotope correction | No |
| MS Level for quantification | $MS^1$ | Limit of quantification | No |
| Internal lipid standard(s) $MS^1$ | | Normalization to reference | No |
| Type of quantification | Internal standard amount | Lipid Quantification Software | MS-DIAL |
| Response correction | No | Batch correction | No |

| Internal standard | Endogenous subclass |
|---|---|
| LPC 18:1(d7) | EtherMGDG subclass |

#### 55) Ether-linked oxidized phosphatidylcholine (EtherOxPC) / Lipid identification

| | | | |
|---|---|---|---|
| Lipid class | Ether-linked oxidized phosphatidylcholine (EtherOxPC) | Did you presume assumptions for identification? | No |
| MS Level for identification | $MS^1$, $MS^2$ | Limit of detection | No |
| Identification level | Molecular species level | RT verified by standard | Yes |
| Isotope correction at $MS^1$ | No | Separation of isobaric/isomeric interferece confirmed | Yes |
| Fragments for identification | | Model for separation prediction | Yes |

| Fragment name |
|---|
| Oxidized fatty acid fragment |
| Oxidized fatty acid -H2O fragment |
| Neutral loss of methyl moiety |

| | | | |
|---|---|---|---|
| Isotope correction at $MS^2$ | No | Lipid Identification Software | MS-DIAL |
| $MS^1$ verified by standard | No | Data manipulation | Smoothing, Centroiding |
| $MS^2$ verified by standard | No | Nomenclature for intact lipid molecule | Yes |
| Background check at $MS^1$ | Yes | Nomenclature for fragment ions | No |
| Background check at $MS^2$ | No | | |

#### 55) Ether-linked oxidized phosphatidylcholine (EtherOxPC) / Lipid quantification

| | | | |
|---|---|---|---|
| Quantitative | Yes | Type I isotope correction | No |
| MS Level for quantification | $MS^1$ | Limit of quantification | No |
| Internal lipid standard(s) $MS^1$ | | Normalization to reference | No |

| Internal standard | Endogenous subclass |
|---|---|
| PC 15:0_18:1(d7) | EtherOxPC subclass |

| | | | |
|---|---|---|---|
| Type of quantification | Internal standard amount | Lipid Quantification Software | MS-DIAL |
| Response correction | No | Batch correction | No |

#### 56) Ether-linked oxidized phosphatidylethanolamine (EtherOxPE) / Lipid identification

| | | | |
|---|---|---|---|
| Lipid class | Ether-linked oxidized phosphatidylethanolamine (EtherOxPE) | Did you presume assumptions for identification? | No |
| MS Level for identification | $MS^1$, $MS^2$ | Limit of detection | No |
| Identification level | Molecular species level | RT verified by standard | Yes |
| Isotope correction at $MS^1$ | No | Separation of isobaric/isomeric interferece confirmed | Yes |
| Fragments for identification | | Model for separation prediction | Yes |
| Isotope correction at $MS^2$ | No | Lipid Identification Software | MS-DIAL |
| $MS^1$ verified by standard | No | Data manipulation | Smoothing, Centroiding |
| $MS^2$ verified by standard | No | Nomenclature for intact lipid molecule | Yes |
| Background check at $MS^1$ | Yes | Nomenclature for fragment ions | No |
| Background check at $MS^2$ | No | | |

Fragments for identification:

| Fragment name |
|---|
| Oxidized fatty acid fragment |
| Oxidized fatty acid -H2O fragment |

#### 56) Ether-linked oxidized phosphatidylethanolamine (EtherOxPE) / Lipid quantification

| | | | |
|---|---|---|---|
| Quantitative | Yes | Type I isotope correction | No |
| MS Level for quantification | $MS^1$ | Limit of quantification | No |
| Internal lipid standard(s) $MS^1$ | | Normalization to reference | No |
| Type of quantification | Internal standard amount | Lipid Quantification Software | MS-DIAL |
| Response correction | No | Batch correction | No |

Internal lipid standard(s) $MS^1$:

| Internal standard | Endogenous subclass |
|---|---|
| PE 15:0_18:1(d7) | EtherOxPE subclass |

#### 57) PC O / Lipid identification

| | | | |
|---|---|---|---|
| Lipid class | PC O | Did you presume assumptions for identification? | No |
| MS Level for identification | $MS^1$, $MS^2$ | Limit of detection | No |
| Identification level | Molecular species level | RT verified by standard | Yes |
| Isotope correction at $MS^1$ | No | Separation of isobaric/isomeric interferece confirmed | Yes |
| Fragments for identification | | Model for separation prediction | Yes |

| Fragment name |
|---|
| Neutral loss of methyl moiety |
| Fatty acid fragment |

| | | | |
|---|---|---|---|
| Isotope correction at $MS^2$ | No | Lipid Identification Software | MS-DIAL |
| $MS^1$ verified by standard | No | Data manipulation | Smoothing, Centroiding |
| $MS^2$ verified by standard | No | Nomenclature for intact lipid molecule | Yes |
| Background check at $MS^1$ | Yes | Nomenclature for fragment ions | No |
| Background check at $MS^2$ | No | | |

#### 57) PC O / Lipid quantification

| | | | |
|---|---|---|---|
| Quantitative | Yes | Type I isotope correction | No |
| MS Level for quantification | $MS^1$ | Limit of quantification | No |
| Internal lipid standard(s) $MS^1$ | | Normalization to reference | No |

| Internal standard | Endogenous subclass |
|---|---|
| PC 15:0_18:1(d7) | EtherPC subclass |

| | | | |
|---|---|---|---|
| Type of quantification | Internal standard amount | Lipid Quantification Software | MS-DIAL |
| Response correction | No | Batch correction | No |

#### 58) PE O / Lipid identification

| | | | |
|---|---|---|---|
| Lipid class | PE O | Did you presume assumptions for identification? | No |
| MS Level for identification | $MS^1$, $MS^2$ | Limit of detection | No |
| Identification level | Molecular species level | RT verified by standard | Yes |
| Isotope correction at $MS^1$ | No | Separation of isobaric/isomeric interferece confirmed | Yes |
| Fragments for identification | | Model for separation prediction | Yes |

| Fragment name |
|---|
| Neutral loss of fatty acyl |
| Fatty acid fragment |

| | | | |
|---|---|---|---|
| Isotope correction at $MS^2$ | No | Lipid Identification Software | MS-DIAL |
| $MS^1$ verified by standard | No | Data manipulation | Smoothing, Centroiding |
| $MS^2$ verified by standard | No | Nomenclature for intact lipid molecule | Yes |
| Background check at $MS^1$ | Yes | Nomenclature for fragment ions | No |
| Background check at $MS^2$ | No | | |

#### 58) PE O / Lipid quantification

| | | | |
|---|---|---|---|
| Quantitative | Yes | Type I isotope correction | No |
| MS Level for quantification | $MS^1$ | Limit of quantification | No |
| Internal lipid standard(s) $MS^1$ | | Normalization to reference | No |

| Internal standard | Endogenous subclass |
|---|---|
| PE 15:0_18:1(d7) | EtherPE subclass |

| | | | |
|---|---|---|---|
| Type of quantification | Internal standard amount | Lipid Quantification Software | MS-DIAL |
| Response correction | No | Batch correction | No |

#### 59) Ether-linked phosphatidylglycerol (EtherPG) / Lipid identification

| | | | |
|---|---|---|---|
| Lipid class | Ether-linked phosphatidylglycerol (EtherPG) | Did you presume assumptions for identification? | No |
| MS Level for identification | $MS^1$, $MS^2$ | Limit of detection | No |
| Identification level | Molecular species level | RT verified by standard | Yes |
| Isotope correction at $MS^1$ | No | Separation of isobaric/isomeric interferece confirmed | Yes |
| Fragments for identification | | Model for separation prediction | Yes |

| Fragment name |
|---|
| Phosphoglycerol -H2O |
| Fatty acid fragment |
| Alkyl ether +O fragment |

| | | | |
|---|---|---|---|
| Isotope correction at $MS^2$ | No | Lipid Identification Software | MS-DIAL |
| $MS^1$ verified by standard | No | Data manipulation | Smoothing, Centroiding |
| $MS^2$ verified by standard | No | Nomenclature for intact lipid molecule | Yes |
| Background check at $MS^1$ | Yes | Nomenclature for fragment ions | No |
| Background check at $MS^2$ | No | | |

#### 59) Ether-linked phosphatidylglycerol (EtherPG) / Lipid quantification

| | | | |
|---|---|---|---|
| Quantitative | Yes | Type I isotope correction | No |
| MS Level for quantification | $MS^1$ | Limit of quantification | No |
| Internal lipid standard(s) $MS^1$ | | Normalization to reference | No |

| Internal standard | Endogenous subclass |
|---|---|
| PG 15:0_18:1(d7) | EtherPG subclass |

| | | | |
|---|---|---|---|
| Type of quantification | Internal standard amount | Lipid Quantification Software | MS-DIAL |
| Response correction | No | Batch correction | No |

#### 60) Ether-linked phosphatidylinositol (EtherPI) / Lipid identification

| | | | |
|---|---|---|---|
| Lipid class | Ether-linked phosphatidylinositol (EtherPI) | Did you presume assumptions for identification? | No |
| MS Level for identification | $MS^1$, $MS^2$ | Limit of detection | No |
| Identification level | Molecular species level | RT verified by standard | Yes |
| Isotope correction at $MS^1$ | No | Separation of isobaric/isomeric interferece confirmed | Yes |
| Fragments for identification | | Model for separation prediction | Yes |

| Fragment name |
|---|
| Phosphoinositol -H2O |
| Fatty acid fragment |
| Alkyl ether + C3H5O4P fragment |

| | | | |
|---|---|---|---|
| Isotope correction at $MS^2$ | No | Lipid Identification Software | MS-DIAL |
| $MS^1$ verified by standard | No | Data manipulation | Smoothing, Centroiding |
| $MS^2$ verified by standard | No | Nomenclature for intact lipid molecule | Yes |
| Background check at $MS^1$ | Yes | Nomenclature for fragment ions | No |
| Background check at $MS^2$ | No | | |

#### 60) Ether-linked phosphatidylinositol (EtherPI) / Lipid quantification

| | | | |
|---|---|---|---|
| Quantitative | Yes | Type I isotope correction | No |
| MS Level for quantification | $MS^1$ | Limit of quantification | No |
| Internal lipid standard(s) $MS^1$ | | Normalization to reference | No |

| Internal standard | Endogenous subclass |
|---|---|
| PI 15:0_18:1(d7) | EtherPI subclass |

| | | | |
|---|---|---|---|
| Type of quantification | Internal standard amount | Lipid Quantification Software | MS-DIAL |
| Response correction | No | Batch correction | No |

#### 61) Ether-linked phosphatidylserine (EtherPS) / Lipid identification

| | | | |
|---|---|---|---|
| Lipid class | Ether-linked phosphatidylserine (EtherPS) | Did you presume assumptions for identification? | No |
| MS Level for identification | $MS^1$, $MS^2$ | Limit of detection | No |
| Identification level | Molecular species level | RT verified by standard | Yes |
| Isotope correction at $MS^1$ | No | Separation of isobaric/isomeric interferece confirmed | Yes |
| Fragments for identification | | Model for separation prediction | Yes |

| Fragment name |
|---|
| Neutral loss of C3H5NO2 |
| Neutral loss of 22:6 Acyl and H2O and C3H5NO2 |
| Fatty acid fragment |

| | | | |
|---|---|---|---|
| Isotope correction at $MS^2$ | No | Lipid Identification Software | MS-DIAL |
| $MS^1$ verified by standard | No | Data manipulation | Smoothing, Centroiding |
| $MS^2$ verified by standard | No | Nomenclature for intact lipid molecule | Yes |
| Background check at $MS^1$ | Yes | Nomenclature for fragment ions | No |
| Background check at $MS^2$ | No | | |

#### 61) Ether-linked phosphatidylserine (EtherPS) / Lipid quantification

| | | | |
|---|---|---|---|
| Quantitative | Yes | Type I isotope correction | No |
| MS Level for quantification | $MS^1$ | Limit of quantification | No |
| Internal lipid standard(s) $MS^1$ | | Normalization to reference | No |

| Internal standard | Endogenous subclass |
|---|---|
| PS 15:0_18:1(d7) | EtherPS subclass |

| | | | |
|---|---|---|---|
| Type of quantification | Internal standard amount | Lipid Quantification Software | MS-DIAL |
| Response correction | No | Batch correction | No |

#### 62) Ether-linked triacylglycerol (EtherTG) / Lipid identification

| | | | |
|---|---|---|---|
| Lipid class | Ether-linked triacylglycerol (EtherTG) | Did you presume assumptions for identification? | No |
| MS Level for identification | $MS^1$, $MS^2$ | Limit of detection | No |
| Identification level | Molecular species level | RT verified by standard | Yes |
| Isotope correction at $MS^1$ | No | Separation of isobaric/isomeric interferece confirmed | Yes |
| Fragments for identification | | Model for separation prediction | Yes |

| Fragment name |
|---|
| Neutral loss of fatty acyl and H2O |
| Neutral loss of alkyl ether |

| | | | |
|---|---|---|---|
| Isotope correction at $MS^2$ | No | Lipid Identification Software | MS-DIAL |
| $MS^1$ verified by standard | No | Data manipulation | Smoothing, Centroiding |
| $MS^2$ verified by standard | No | Nomenclature for intact lipid molecule | Yes |
| Background check at $MS^1$ | Yes | Nomenclature for fragment ions | No |
| Background check at $MS^2$ | No | | |

#### 62) Ether-linked triacylglycerol (EtherTG) / Lipid quantification

| | | | |
|---|---|---|---|
| Quantitative | Yes | Type I isotope correction | No |
| MS Level for quantification | $MS^1$ | Limit of quantification | No |
| Internal lipid standard(s) $MS^1$ | | Normalization to reference | No |

| Internal standard | Endogenous subclass |
|---|---|
| TG 15:0_18:1(d7)_15:0 | EtherTG subclass |

| | | | |
|---|---|---|---|
| Type of quantification | Internal standard amount | Lipid Quantification Software | MS-DIAL |
| Response correction | No | Batch correction | No |

#### 63) Fatty acid ester of hydroxyl fatty acid (FAHFA) / Lipid identification

| | | | |
|---|---|---|---|
| Lipid class | Fatty acid ester of hydroxyl fatty acid (FAHFA) | Did you presume assumptions for identification? | No |
| MS Level for identification | $MS^1$, $MS^2$ | Limit of detection | No |
| Identification level | Molecular species level | RT verified by standard | Yes |
| Isotope correction at $MS^1$ | No | Separation of isobaric/isomeric interferece confirmed | Yes |
| Fragments for identification | | Model for separation prediction | Yes |

| Fragment name |
|---|
| Neutral loss of fatty acyl and H2O |
| Fatty acid fragment |

| | | | |
|---|---|---|---|
| Isotope correction at $MS^2$ | No | Lipid Identification Software | MS-DIAL |
| $MS^1$ verified by standard | No | Data manipulation | Smoothing, Centroiding |
| $MS^2$ verified by standard | No | Nomenclature for intact lipid molecule | Yes |
| Background check at $MS^1$ | Yes | Nomenclature for fragment ions | No |
| Background check at $MS^2$ | No | | |

#### 63) Fatty acid ester of hydroxyl fatty acid (FAHFA) / Lipid quantification

| | | | |
|---|---|---|---|
| Quantitative | Yes | Type I isotope correction | No |
| MS Level for quantification | $MS^1$ | Limit of quantification | No |
| Internal lipid standard(s) $MS^1$ | | Normalization to reference | No |

| Internal standard | Endogenous subclass |
|---|---|
| FA 18:0(d3) | FAHFA subclass |

| | | | |
|---|---|---|---|
| Type of quantification | Internal standard amount | Lipid Quantification Software | MS-DIAL |
| Response correction | No | Batch correction | No |

#### 64) Ganglioside GD1a (GD1a) / Lipid identification

| | | | |
|---|---|---|---|
| Lipid class | Ganglioside GD1a (GD1a) | Did you presume assumptions for identification? | No |
| MS Level for identification | $MS^1$, $MS^2$ | Limit of detection | No |
| Identification level | Species level | RT verified by standard | Yes |
| Isotope correction at $MS^1$ | No | Separation of isobaric/isomeric interferece confirmed | Yes |
| Fragments for identification | | Model for separation prediction | Yes |

| Fragment name |
|---|
| Characteristic fragment (C11H16NO8-) |

| | | | |
|---|---|---|---|
| Isotope correction at $MS^2$ | No | Lipid Identification Software | MS-DIAL |
| $MS^1$ verified by standard | No | Data manipulation | Smoothing, Centroiding |
| $MS^2$ verified by standard | No | Nomenclature for intact lipid molecule | Yes |
| Background check at $MS^1$ | Yes | Nomenclature for fragment ions | No |
| Background check at $MS^2$ | No | | |

#### 64) Ganglioside GD1a (GD1a) / Lipid quantification

| | | | |
|---|---|---|---|
| Quantitative | Yes | Type I isotope correction | No |
| MS Level for quantification | $MS^1$ | Limit of quantification | No |
| Internal lipid standard(s) $MS^1$ | | Normalization to reference | No |
| Type of quantification | Internal standard amount | Lipid Quantification Software | MS-DIAL |
| Response correction | No | Batch correction | No |

| Internal standard | Endogenous subclass |
|---|---|
| LPC 18:1(d7) | GD1a subclass |

#### 65) Ganglioside GD1b (GD1b) / Lipid identification

| | | | |
|---|---|---|---|
| Lipid class | Ganglioside GD1b (GD1b) | Did you presume assumptions for identification? | No |
| MS Level for identification | $MS^1$, $MS^2$ | Limit of detection | No |
| Identification level | Species level | RT verified by standard | Yes |
| Isotope correction at $MS^1$ | No | Separation of isobaric/isomeric interferece confirmed | Yes |
| Fragments for identification | | Model for separation prediction | Yes |
| Isotope correction at $MS^2$ | No | Lipid Identification Software | MS-DIAL |
| $MS^1$ verified by standard | No | Data manipulation | Smoothing, Centroiding |
| $MS^2$ verified by standard | No | Nomenclature for intact lipid molecule | Yes |
| Background check at $MS^1$ | Yes | Nomenclature for fragment ions | No |
| Background check at $MS^2$ | No | | |

| Fragment name |
|---|
| Characteristic fragment (C11H16NO8-) |

#### 65) Ganglioside GD1b (GD1b) / Lipid quantification

| | | | |
|---|---|---|---|
| Quantitative | Yes | Type I isotope correction | No |
| MS Level for quantification | $MS^1$ | Limit of quantification | No |
| Internal lipid standard(s) $MS^1$ | | Normalization to reference | No |
| Type of quantification | Internal standard amount | Lipid Quantification Software | MS-DIAL |
| Response correction | No | Batch correction | No |

| Internal standard | Endogenous subclass |
|---|---|
| LPC 18:1(d7) | GD1b subclass |

#### 66) Ganglioside GD2 (GD2) / Lipid identification

| | | | |
|---|---|---|---|
| Lipid class | Ganglioside GD2 (GD2) | Did you presume assumptions for identification? | No |
| MS Level for identification | $MS^1$, $MS^2$ | Limit of detection | No |
| Identification level | Species level | RT verified by standard | Yes |
| Isotope correction at $MS^1$ | No | Separation of isobaric/isomeric interferece confirmed | Yes |
| Fragments for identification | | Model for separation prediction | Yes |

| Fragment name |
|---|
| Characteristic fragment (C11H16NO8-) |

| | | | |
|---|---|---|---|
| Isotope correction at $MS^2$ | No | Lipid Identification Software | MS-DIAL |
| $MS^1$ verified by standard | No | Data manipulation | Smoothing, Centroiding |
| $MS^2$ verified by standard | No | Nomenclature for intact lipid molecule | Yes |
| Background check at $MS^1$ | Yes | Nomenclature for fragment ions | No |
| Background check at $MS^2$ | No | | |

#### 66) Ganglioside GD2 (GD2) / Lipid quantification

| | | | |
|---|---|---|---|
| Quantitative | Yes | Type I isotope correction | No |
| MS Level for quantification | $MS^1$ | Limit of quantification | No |
| Internal lipid standard(s) $MS^1$ | | Normalization to reference | No |

| Internal standard | Endogenous subclass |
|---|---|
| LPC 18:1(d7) | GD2 subclass |

| | | | |
|---|---|---|---|
| Type of quantification | Internal standard amount | Lipid Quantification Software | MS-DIAL |
| Response correction | No | Batch correction | No |

#### 67) GD3 / Lipid identification

| | | | |
|---|---|---|---|
| Lipid class | GD3 | Did you presume assumptions for identification? | No |
| MS Level for identification | $MS^1$, $MS^2$ | Limit of detection | No |
| Identification level | Species level | RT verified by standard | Yes |
| Isotope correction at $MS^1$ | No | Separation of isobaric/isomeric interferece confirmed | Yes |
| Fragments for identification | | Model for separation prediction | Yes |

| Fragment name |
|---|
| Characteristic fragment (C11H16NO8-) |

| | | | |
|---|---|---|---|
| Isotope correction at $MS^2$ | No | Lipid Identification Software | MS-DIAL |
| $MS^1$ verified by standard | No | Data manipulation | Smoothing, Centroiding |
| $MS^2$ verified by standard | No | Nomenclature for intact lipid molecule | Yes |
| Background check at $MS^1$ | Yes | Nomenclature for fragment ions | No |
| Background check at $MS^2$ | No | | |

#### 67) GD3 / Lipid quantification

| | | | |
|---|---|---|---|
| Quantitative | Yes | Type I isotope correction | No |
| MS Level for quantification | $MS^1$ | Limit of quantification | No |
| Internal lipid standard(s) $MS^1$ | | Normalization to reference | No |

| Internal standard | Endogenous subclass |
|---|---|
| LPC 18:1(d7) | GD3 subclass |

| | | | |
|---|---|---|---|
| Type of quantification | Internal standard amount | Lipid Quantification Software | MS-DIAL |
| Response correction | No | Batch correction | No |

#### 68) GM1 / Lipid identification

| | | | |
|---|---|---|---|
| Lipid class | GM1 | Did you presume assumptions for identification? | No |
| MS Level for identification | $MS^1$, $MS^2$ | Limit of detection | No |
| Identification level | Species level | RT verified by standard | Yes |
| Isotope correction at $MS^1$ | No | Separation of isobaric/isomeric interferece confirmed | Yes |
| Fragments for identification | | Model for separation prediction | Yes |

| Fragment name |
|---|
| Characteristic fragment (C11H16NO8-) |

| | | | |
|---|---|---|---|
| Isotope correction at $MS^2$ | No | Lipid Identification Software | MS-DIAL |
| $MS^1$ verified by standard | No | Data manipulation | Smoothing, Centroiding |
| $MS^2$ verified by standard | No | Nomenclature for intact lipid molecule | Yes |
| Background check at $MS^1$ | Yes | Nomenclature for fragment ions | No |
| Background check at $MS^2$ | No | | |

#### 68) GM1 / Lipid quantification

| | | | |
|---|---|---|---|
| Quantitative | Yes | Type I isotope correction | No |
| MS Level for quantification | $MS^1$ | Limit of quantification | No |
| Internal lipid standard(s) $MS^1$ | | Normalization to reference | No |

| Internal standard | Endogenous subclass |
|---|---|
| LPC 18:1(d7) | GM1 subclass |

| | | | |
|---|---|---|---|
| Type of quantification | Internal standard amount | Lipid Quantification Software | MS-DIAL |
| Response correction | No | Batch correction | No |

#### 69) GM3 / Lipid identification

| | | | |
|---|---|---|---|
| Lipid class | GM3 | Did you presume assumptions for identification? | No |
| MS Level for identification | $MS^1$, $MS^2$ | Limit of detection | No |
| Identification level | Species level | RT verified by standard | Yes |
| Isotope correction at $MS^1$ | No | Separation of isobaric/isomeric interferece confirmed | Yes |
| Fragments for identification | | Model for separation prediction | Yes |

| Fragment name |
|---|
| Neutral loss of H2O |
| Neutral loss of H2O and C23H37NO18 |
| Sphingosine -H2O fragment |
| Sphingosine -2H2O fragment |

| | | | |
|---|---|---|---|
| Isotope correction at $MS^2$ | No | Lipid Identification Software | MS-DIAL |
| $MS^1$ verified by standard | No | Data manipulation | Smoothing, Centroiding |
| $MS^2$ verified by standard | No | Nomenclature for intact lipid molecule | Yes |
| Background check at $MS^1$ | Yes | Nomenclature for fragment ions | No |
| Background check at $MS^2$ | No | | |

#### 69) GM3 / Lipid quantification

| | | | |
|---|---|---|---|
| Quantitative | Yes | Type I isotope correction | No |
| MS Level for quantification | $MS^1$ | Limit of quantification | No |
| Internal lipid standard(s) $MS^1$ | | Normalization to reference | No |

| Internal standard | Endogenous subclass |
|---|---|
| LPC 18:1(d7) | GM3 subclass |

| | | | |
|---|---|---|---|
| Type of quantification | Internal standard amount | Lipid Quantification Software | MS-DIAL |
| Response correction | No | Batch correction | No |

#### 70) Ganglioside GQ1b (GQ1b) / Lipid identification

| | | | |
|---|---|---|---|
| Lipid class | Ganglioside GQ1b (GQ1b) | Did you presume assumptions for identification? | No |
| MS Level for identification | $MS^1$, $MS^2$ | Limit of detection | No |
| Identification level | Species level | RT verified by standard | Yes |
| Isotope correction at $MS^1$ | No | Separation of isobaric/isomeric interferece confirmed | Yes |
| Fragments for identification | | Model for separation prediction | Yes |

| Fragment name |
|---|
| Characteristic fragment (C11H16NO8-) |
| Characteristic fragment (C22H33N2O16-) |

| | | | |
|---|---|---|---|
| Isotope correction at $MS^2$ | No | Lipid Identification Software | MS-DIAL |
| $MS^1$ verified by standard | No | Data manipulation | Smoothing, Centroiding |
| $MS^2$ verified by standard | No | Nomenclature for intact lipid molecule | Yes |
| Background check at $MS^1$ | Yes | Nomenclature for fragment ions | No |
| Background check at $MS^2$ | No | | |

#### 70) Ganglioside GQ1b (GQ1b) / Lipid quantification

| | | | |
|---|---|---|---|
| Quantitative | Yes | Type I isotope correction | No |
| MS Level for quantification | $MS^1$ | Limit of quantification | No |
| Internal lipid standard(s) $MS^1$ | | Normalization to reference | No |

| Internal standard | Endogenous subclass |
|---|---|
| LPC 18:1(d7) | GQ1b subclass |

| | | | |
|---|---|---|---|
| Type of quantification | Internal standard amount | Lipid Quantification Software | MS-DIAL |
| Response correction | No | Batch correction | No |

#### 71) Ganglioside GT1b (GT1b) / Lipid identification

| | | | |
|---|---|---|---|
| Lipid class | Ganglioside GT1b (GT1b) | Did you presume assumptions for identification? | No |
| MS Level for identification | $MS^1$, $MS^2$ | Limit of detection | No |
| Identification level | Species level | RT verified by standard | Yes |
| Isotope correction at $MS^1$ | No | Separation of isobaric/isomeric interferece confirmed | Yes |
| Fragments for identification | | Model for separation prediction | Yes |

| Fragment name |
|---|
| Characteristic fragment (C11H16NO8-) |
| Characteristic fragment (C22H33N2O16-) |

| | | | |
|---|---|---|---|
| Isotope correction at $MS^2$ | No | Lipid Identification Software | MS-DIAL |
| $MS^1$ verified by standard | No | Data manipulation | Smoothing, Centroiding |
| $MS^2$ verified by standard | No | Nomenclature for intact lipid molecule | Yes |
| Background check at $MS^1$ | Yes | Nomenclature for fragment ions | No |
| Background check at $MS^2$ | No | | |

#### 71) Ganglioside GT1b (GT1b) / Lipid quantification

| | | | |
|---|---|---|---|
| Quantitative | Yes | Type I isotope correction | No |
| MS Level for quantification | $MS^1$ | Limit of quantification | No |
| Internal lipid standard(s) $MS^1$ | | Normalization to reference | No |

| Internal standard | Endogenous subclass |
|---|---|
| LPC 18:1(d7) | GT1b subclass |

| | | | |
|---|---|---|---|
| Type of quantification | Internal standard amount | Lipid Quantification Software | MS-DIAL |
| Response correction | No | Batch correction | No |

#### 72) Glycerophospho N-acyl ethanolamine (GPNAE) / Lipid identification

| | | | |
|---|---|---|---|
| Lipid class | Glycerophospho N-acyl ethanolamine (GPNAE) | Did you presume assumptions for identification? | No |
| MS Level for identification | $MS^1$, $MS^2$ | Limit of detection | No |
| Identification level | Molecular species level | RT verified by standard | Yes |
| Isotope correction at $MS^1$ | No | Separation of isobaric/isomeric interferece confirmed | Yes |
| Fragments for identification | | Model for separation prediction | Yes |

| Fragment name |
|---|
| Characteristic fragment (C3H8PO6-) |
| Phosphite |

| | | | |
|---|---|---|---|
| Isotope correction at $MS^2$ | No | Lipid Identification Software | MS-DIAL |
| $MS^1$ verified by standard | No | Data manipulation | Smoothing, Centroiding |
| $MS^2$ verified by standard | No | Nomenclature for intact lipid molecule | Yes |
| Background check at $MS^1$ | Yes | Nomenclature for fragment ions | No |
| Background check at $MS^2$ | No | | |

#### 72) Glycerophospho N-acyl ethanolamine (GPNAE) / Lipid quantification

| | | | |
|---|---|---|---|
| Quantitative | Yes | Type I isotope correction | No |
| MS Level for quantification | $MS^1$ | Limit of quantification | No |
| Internal lipid standard(s) $MS^1$ | | Normalization to reference | No |

| Internal standard | Endogenous subclass |
|---|---|
| LPC 18:1(d7) | GPNAE subclass |

| | | | |
|---|---|---|---|
| Type of quantification | Internal standard amount | Lipid Quantification Software | MS-DIAL |
| Response correction | No | Batch correction | No |

#### 73) Hemibismonoacylglycerophosphate (HBMP) / Lipid identification

| | | | |
|---|---|---|---|
| Lipid class | Hemibismonoacylglycerophosphate (HBMP) | Did you presume assumptions for identification? | No |
| MS Level for identification | $MS^1$, $MS^2$ | Limit of detection | No |
| Identification level | Molecular species level | RT verified by standard | Yes |
| Isotope correction at $MS^1$ | No | Separation of isobaric/isomeric interferece confirmed | Yes |
| Fragments for identification | | Model for separation prediction | Yes |

| Fragment name |
|---|
| Dehydro-monoacyl glycerols |
| Dehydro-diacyl glycerol |

| | | | |
|---|---|---|---|
| Isotope correction at $MS^2$ | No | Lipid Identification Software | MS-DIAL |
| $MS^1$ verified by standard | No | Data manipulation | Smoothing, Centroiding |
| $MS^2$ verified by standard | No | Nomenclature for intact lipid molecule | Yes |
| Background check at $MS^1$ | Yes | Nomenclature for fragment ions | No |
| Background check at $MS^2$ | No | | |

#### 73) Hemibismonoacylglycerophosphate (HBMP) / Lipid quantification

| | | | |
|---|---|---|---|
| Quantitative | Yes | Type I isotope correction | No |
| MS Level for quantification | $MS^1$ | Limit of quantification | No |
| Internal lipid standard(s) $MS^1$ | | Normalization to reference | No |

| Internal standard | Endogenous subclass |
|---|---|
| PG 15:0_18:1(d7) | HBMP subclass |

| | | | |
|---|---|---|---|
| Type of quantification | Internal standard amount | Lipid Quantification Software | MS-DIAL |
| Response correction | No | Batch correction | No |

#### 74) Hexosylceramide alpha-hydroxy fatty acid-phytospingosine (HexCer_AP) / Lipid identification

| | | | |
|---|---|---|---|
| Lipid class | Hexosylceramide alpha-hydroxy fatty acid-phytospingosine (HexCer_AP) | Did you presume assumptions for identification? | No |
| MS Level for identification | $MS^1$, $MS^2$ | Limit of detection | No |
| Identification level | Molecular species level | RT verified by standard | Yes |
| Isotope correction at $MS^1$ | No | Separation of isobaric/isomeric interferece confirmed | Yes |
| Fragments for identification | | Model for separation prediction | Yes |

| Fragment name |
|---|
| Neutral loss of hexose |
| Neutral loss of hexose and H2O |
| Phytosphingosine |
| Phytosphingosine -H2O fragment |
| Phytosphingosine -3H2O fragment |

| | | | |
|---|---|---|---|
| Isotope correction at $MS^2$ | No | Lipid Identification Software | MS-DIAL |
| $MS^1$ verified by standard | No | Data manipulation | Smoothing, Centroiding |
| $MS^2$ verified by standard | No | Nomenclature for intact lipid molecule | Yes |
| Background check at $MS^1$ | Yes | Nomenclature for fragment ions | No |
| Background check at $MS^2$ | No | | |

#### 74) Hexosylceramide alpha-hydroxy fatty acid-phytospingosine (HexCer_AP) / Lipid quantification

| | | | |
|---|---|---|---|
| Quantitative | Yes | Type I isotope correction | No |
| MS Level for quantification | $MS^1$ | Limit of quantification | No |
| Internal lipid standard(s) $MS^1$ | | Normalization to reference | No |

| Internal standard | Endogenous subclass |
|---|---|
| Cer 18:1;2O/15:0(d7) | HexCer_AP subclass |

| | | | |
|---|---|---|---|
| Type of quantification | Internal standard amount | Lipid Quantification Software | MS-DIAL |
| Response correction | No | Batch correction | No |

#### 75) Hexosylceramide Esterified omega-hydroxy fatty acid-sphingosine (HexCer_EOS) / Lipid identification

| | | | |
|---|---|---|---|
| Lipid class | Hexosylceramide Esterified omega-hydroxy fatty acid-sphingosine (HexCer_EOS) | Did you presume assumptions for identification? | No |
| MS Level for identification | $MS^1$, $MS^2$ | Limit of detection | No |
| Identification level | Molecular species level | RT verified by standard | Yes |
| Isotope correction at $MS^1$ | No | Separation of isobaric/isomeric interferece confirmed | Yes |
| Fragments for identification | | Model for separation prediction | Yes |

| Fragment name |
|---|
| Neutral loss of hexose |
| Neutral loss of hexose and H2O |
| Sphingosine -H2O fragment |
| Sphingosine -2H2O fragment |
| Sphingosine -CH4O2 fragment |

| | | | |
|---|---|---|---|
| Isotope correction at $MS^2$ | No | Lipid Identification Software | MS-DIAL |
| $MS^1$ verified by standard | No | Data manipulation | Smoothing, Centroiding |
| $MS^2$ verified by standard | No | Nomenclature for intact lipid molecule | Yes |
| Background check at $MS^1$ | Yes | Nomenclature for fragment ions | No |
| Background check at $MS^2$ | No | | |

#### 75) Hexosylceramide Esterified omega-hydroxy fatty acid-sphingosine (HexCer_EOS) / Lipid quantification

| | | | |
|---|---|---|---|
| Quantitative | Yes | Type I isotope correction | No |
| MS Level for quantification | $MS^1$ | Limit of quantification | No |
| Internal lipid standard(s) $MS^1$ | | Normalization to reference | No |

| Internal standard | Endogenous subclass |
|---|---|
| Cer 18:1;2O/15:0(d7) | HexCer_EOS subclass |

| | | | |
|---|---|---|---|
| Type of quantification | Internal standard amount | Lipid Quantification Software | MS-DIAL |
| Response correction | No | Batch correction | No |

#### 76) Hexosylceramide hydroxyfatty acid-dihydrosphingosine (HexCer_HDS) / Lipid identification

| | | | |
|---|---|---|---|
| Lipid class | Hexosylceramide hydroxyfatty acid-dihydrosphingosine (HexCer_HDS) | Did you presume assumptions for identification? | No |
| MS Level for identification | $MS^1$, $MS^2$ | Limit of detection | No |
| Identification level | Molecular species level | RT verified by standard | Yes |
| Isotope correction at $MS^1$ | No | Separation of isobaric/isomeric interferece confirmed | Yes |
| Fragments for identification | | Model for separation prediction | Yes |

| Fragment name |
|---|
| Neutral loss of hexose |
| Neutral loss of hexose and H2O |
| Sphinganine -H2O fragment |
| Sphinganine -2H2O fragment |
| Sphinganine -CH4O2 fragment |

| | | | |
|---|---|---|---|
| Isotope correction at $MS^2$ | No | Lipid Identification Software | MS-DIAL |
| $MS^1$ verified by standard | No | Data manipulation | Smoothing, Centroiding |
| $MS^2$ verified by standard | No | Nomenclature for intact lipid molecule | Yes |
| Background check at $MS^1$ | Yes | Nomenclature for fragment ions | No |
| Background check at $MS^2$ | No | | |

#### 76) Hexosylceramide hydroxyfatty acid-dihydrosphingosine (HexCer_HDS) / Lipid quantification

| | | | |
|---|---|---|---|
| Quantitative | Yes | Type I isotope correction | No |
| MS Level for quantification | $MS^1$ | Limit of quantification | No |
| Internal lipid standard(s) $MS^1$ | | Normalization to reference | No |

| Internal standard | Endogenous subclass |
|---|---|
| Cer 18:1;2O/15:0(d7) | HexCer_HDS subclass |

| | | | |
|---|---|---|---|
| Type of quantification | Internal standard amount | Lipid Quantification Software | MS-DIAL |
| Response correction | No | Batch correction | No |

#### 77) Hexosylceramide hydroxyfatty acid-sphingosine (HexCer_HS) / Lipid identification

| | | | |
|---|---|---|---|
| Lipid class | Hexosylceramide hydroxyfatty acid-sphingosine (HexCer_HS) | Did you presume assumptions for identification? | No |
| MS Level for identification | $MS^1$, $MS^2$ | Limit of detection | No |
| Identification level | Molecular species level | RT verified by standard | Yes |
| Isotope correction at $MS^1$ | No | Separation of isobaric/isomeric interferece confirmed | Yes |
| Fragments for identification | | Model for separation prediction | Yes |

| Fragment name |
|---|
| Neutral loss of hexose |
| Neutral loss of hexose and H2O |
| Sphingosine -H2O fragment |
| Sphingosine -2H2O fragment |
| Sphingosine -CH4O2 fragment |

| | | | |
|---|---|---|---|
| Isotope correction at $MS^2$ | No | Lipid Identification Software | MS-DIAL |
| $MS^1$ verified by standard | No | Data manipulation | Smoothing, Centroiding |
| $MS^2$ verified by standard | No | Nomenclature for intact lipid molecule | Yes |
| Background check at $MS^1$ | Yes | Nomenclature for fragment ions | No |
| Background check at $MS^2$ | No | | |

#### 77) Hexosylceramide hydroxyfatty acid-sphingosine (HexCer_HS) / Lipid quantification

| | | | |
|---|---|---|---|
| Quantitative | Yes | Type I isotope correction | No |
| MS Level for quantification | $MS^1$ | Limit of quantification | No |
| Internal lipid standard(s) $MS^1$ | | Normalization to reference | No |

| Internal standard | Endogenous subclass |
|---|---|
| Cer 18:1;2O/15:0(d7) | HexCer_HS subclass |

| | | | |
|---|---|---|---|
| Type of quantification | Internal standard amount | Lipid Quantification Software | MS-DIAL |
| Response correction | No | Batch correction | No |

#### 78) Hexosylceramide non-hydroxyfatty acid-dihydrosphingosine (HexCer_NDS) / Lipid identification

| | | | |
|---|---|---|---|
| Lipid class | Hexosylceramide non-hydroxyfatty acid-dihydrosphingosine (HexCer_NDS) | Did you presume assumptions for identification? | No |
| MS Level for identification | $MS^1$, $MS^2$ | Limit of detection | No |
| Identification level | Molecular species level | RT verified by standard | Yes |
| Isotope correction at $MS^1$ | No | Separation of isobaric/isomeric interferece confirmed | Yes |
| Fragments for identification | | Model for separation prediction | Yes |
| Isotope correction at $MS^2$ | No | Lipid Identification Software | MS-DIAL |
| $MS^1$ verified by standard | No | Data manipulation | Smoothing, Centroiding |
| $MS^2$ verified by standard | No | Nomenclature for intact lipid molecule | Yes |
| Background check at $MS^1$ | Yes | Nomenclature for fragment ions | No |
| Background check at $MS^2$ | No | | |

| Fragment name |
|---|
| Neutral loss of hexose and H2O |
| Sphinganine -H2O fragment |
| Sphinganine -2H2O fragment |

#### 78) Hexosylceramide non-hydroxyfatty acid-dihydrosphingosine (HexCer_NDS) / Lipid quantification

| | | | |
|---|---|---|---|
| Quantitative | Yes | Type I isotope correction | No |
| MS Level for quantification | $MS^1$ | Limit of quantification | No |
| Internal lipid standard(s) $MS^1$ | | Normalization to reference | No |
| Type of quantification | Internal standard amount | Lipid Quantification Software | MS-DIAL |
| Response correction | No | Batch correction | No |

| Internal standard | Endogenous subclass |
|---|---|
| Cer 18:1;2O/15:0(d7) | HexCer_NDS subclass |

#### 79) Hexosylceramide non-hydroxyfatty acid-sphingosine (HexCer_NS) / Lipid identification

| | | | |
|---|---|---|---|
| Lipid class | Hexosylceramide non-hydroxyfatty acid-sphingosine (HexCer_NS) | Did you presume assumptions for identification? | No |
| MS Level for identification | $MS^1$, $MS^2$ | Limit of detection | No |
| Identification level | Molecular species level | RT verified by standard | Yes |
| Isotope correction at $MS^1$ | No | Separation of isobaric/isomeric interferece confirmed | Yes |
| Fragments for identification | | Model for separation prediction | Yes |

| Fragment name |
|---|
| Neutral loss of H2O |
| Neutral loss of hexose and H2O |
| Sphingosine -H2O fragment |
| Sphingosine -2H2O fragment |
| Sphingosine -CH4O2 fragment |

| | | | |
|---|---|---|---|
| Isotope correction at $MS^2$ | No | Lipid Identification Software | MS-DIAL |
| $MS^1$ verified by standard | No | Data manipulation | Smoothing, Centroiding |
| $MS^2$ verified by standard | No | Nomenclature for intact lipid molecule | Yes |
| Background check at $MS^1$ | Yes | Nomenclature for fragment ions | No |
| Background check at $MS^2$ | No | | |

#### 79) Hexosylceramide non-hydroxyfatty acid-sphingosine (HexCer_NS) / Lipid quantification

| | | | |
|---|---|---|---|
| Quantitative | Yes | Type I isotope correction | No |
| MS Level for quantification | $MS^1$ | Limit of quantification | No |
| Internal lipid standard(s) $MS^1$ | | Normalization to reference | No |

| Internal standard | Endogenous subclass |
|---|---|
| Cer 18:1;2O/15:0(d7) | HexCer_NS subclass |

| | | | |
|---|---|---|---|
| Type of quantification | Internal standard amount | Lipid Quantification Software | MS-DIAL |
| Response correction | No | Batch correction | No |

#### 80) Lysocardiolipin (MLCL) / Lipid identification

| | | | |
|---|---|---|---|
| Lipid class | Lysocardiolipin (MLCL) | Did you presume assumptions for identification? | No |
| MS Level for identification | $MS^1$, $MS^2$ | Limit of detection | No |
| Identification level | Molecular species level | RT verified by standard | Yes |
| Isotope correction at $MS^1$ | No | Separation of isobaric/isomeric interferece confirmed | Yes |
| Fragments for identification | | Model for separation prediction | Yes |

| Fragment name |
|---|
| Phosphoglycerol -H2O fragment |
| Phosphatidic acid |
| Lysophosphatidic acid |
| Fatty acid fragment |

| | | | |
|---|---|---|---|
| Isotope correction at $MS^2$ | No | Lipid Identification Software | MS-DIAL |
| $MS^1$ verified by standard | No | Data manipulation | Smoothing, Centroiding |
| $MS^2$ verified by standard | No | Nomenclature for intact lipid molecule | Yes |
| Background check at $MS^1$ | Yes | Nomenclature for fragment ions | No |
| Background check at $MS^2$ | No | | |

#### 80) Lysocardiolipin (MLCL) / Lipid quantification

| | | | |
|---|---|---|---|
| Quantitative | Yes | Type I isotope correction | No |
| MS Level for quantification | $MS^1$ | Limit of quantification | No |
| Internal lipid standard(s) $MS^1$ | | Normalization to reference | No |

| Internal standard | Endogenous subclass |
|---|---|
| PG 15:0_18:1(d7) | MLCL subclass |

| | | | |
|---|---|---|---|
| Type of quantification | Internal standard amount | Lipid Quantification Software | MS-DIAL |
| Response correction | No | Batch correction | No |

#### 81) Lysodiacylglyceryl-3-O-carboxyhydroxymethylcholine (LDGCC) / Lipid identification

| | | | |
|---|---|---|---|
| Lipid class | Lysodiacylglyceryl-3-O-carboxyhydroxymethylcholine (LDGCC) | Did you presume assumptions for identification? | No |
| MS Level for identification | $MS^1$, $MS^2$ | Limit of detection | No |
| Identification level | Molecular species level | RT verified by standard | Yes |
| Isotope correction at $MS^1$ | No | Separation of isobaric/isomeric interferece confirmed | Yes |
| Fragments for identification | | Model for separation prediction | Yes |

| Fragment name |
|---|
| Characteristic fragment (C6H14NO2+) |

| | | | |
|---|---|---|---|
| Isotope correction at $MS^2$ | No | Lipid Identification Software | MS-DIAL |
| $MS^1$ verified by standard | No | Data manipulation | Smoothing, Centroiding |
| $MS^2$ verified by standard | No | Nomenclature for intact lipid molecule | Yes |
| Background check at $MS^1$ | Yes | Nomenclature for fragment ions | No |
| Background check at $MS^2$ | No | | |

#### 81) Lysodiacylglyceryl-3-O-carboxyhydroxymethylcholine (LDGCC) / Lipid quantification

| | | | |
|---|---|---|---|
| Quantitative | Yes | Type I isotope correction | No |
| MS Level for quantification | $MS^1$ | Limit of quantification | No |
| Internal lipid standard(s) $MS^1$ | | Normalization to reference | No |

| Internal standard | Endogenous subclass |
|---|---|
| LPC 18:1(d7) | LDGCC subclass |

| | | | |
|---|---|---|---|
| Type of quantification | Internal standard amount | Lipid Quantification Software | MS-DIAL |
| Response correction | No | Batch correction | No |

#### 82) Lysogiacylglyceryl trimethylhomoserine (LDGTS) / Lipid identification

| | | | |
|---|---|---|---|
| Lipid class | Lysogiacylglyceryl trimethylhomoserine (LDGTS) | Did you presume assumptions for identification? | No |
| MS Level for identification | $MS^1$, $MS^2$ | Limit of detection | No |
| Identification level | Molecular species level | RT verified by standard | Yes |
| Isotope correction at $MS^1$ | No | Separation of isobaric/isomeric interferece confirmed | Yes |
| Fragments for identification | | Model for separation prediction | Yes |

| Fragment name |
|---|
| Characteristic fragments (C7H14NO2+) |
| Characteristic fragments (C10H22NO5+) |

| | | | |
|---|---|---|---|
| Isotope correction at $MS^2$ | No | Lipid Identification Software | MS-DIAL |
| $MS^1$ verified by standard | No | Data manipulation | Smoothing, Centroiding |
| $MS^2$ verified by standard | No | Nomenclature for intact lipid molecule | Yes |
| Background check at $MS^1$ | Yes | Nomenclature for fragment ions | No |
| Background check at $MS^2$ | No | | |

#### 82) Lysogiacylglyceryl trimethylhomoserine (LDGTS) / Lipid quantification

| | | | |
|---|---|---|---|
| Quantitative | Yes | Type I isotope correction | No |
| MS Level for quantification | $MS^1$ | Limit of quantification | No |
| Internal lipid standard(s) $MS^1$ | | Normalization to reference | No |

| Internal standard | Endogenous subclass |
|---|---|
| LPC 18:1(d7) | LDGTS subclass |

| | | | |
|---|---|---|---|
| Type of quantification | Internal standard amount | Lipid Quantification Software | MS-DIAL |
| Response correction | No | Batch correction | No |

#### 83) LPC / Lipid identification

| | | | |
|---|---|---|---|
| Lipid class | LPC | Did you presume assumptions for identification? | No |
| MS Level for identification | $MS^1$, $MS^2$ | Limit of detection | No |
| Identification level | Molecular species level | RT verified by standard | Yes |
| Isotope correction at $MS^1$ | No | Separation of isobaric/isomeric interferece confirmed | Yes |
| Fragments for identification | | Model for separation prediction | Yes |

| Fragment name |
|---|
| Characteristic fragment (C5H15NO4P+) |

| | | | |
|---|---|---|---|
| Isotope correction at $MS^2$ | No | Lipid Identification Software | MS-DIAL |
| $MS^1$ verified by standard | Yes | Data manipulation | Smoothing, Centroiding |
| $MS^2$ verified by standard | Yes | Nomenclature for intact lipid molecule | Yes |
| Background check at $MS^1$ | Yes | Nomenclature for fragment ions | No |
| Background check at $MS^2$ | No | | |

#### 83) LPC / Lipid quantification

| | | | |
|---|---|---|---|
| Quantitative | Yes | Type I isotope correction | No |
| MS Level for quantification | $MS^1$ | Limit of quantification | No |
| Internal lipid standard(s) $MS^1$ | | Normalization to reference | No |

| Internal standard | Endogenous subclass |
|---|---|
| LPC 18:1(d7) | LPC subclass |

| | | | |
|---|---|---|---|
| Type of quantification | Internal standard amount | Lipid Quantification Software | MS-DIAL |
| Response correction | No | Batch correction | No |

#### 84) LPA / Lipid identification

| | | | |
|---|---|---|---|
| Lipid class | LPA | Did you presume assumptions for identification? | No |
| MS Level for identification | $MS^1$, $MS^2$ | Limit of detection | No |
| Identification level | Molecular species level | RT verified by standard | Yes |
| Isotope correction at $MS^1$ | No | Separation of isobaric/isomeric interferece confirmed | Yes |
| Fragments for identification | | Model for separation prediction | Yes |

| Fragment name |
|---|
| Phosphoglycerol -H2O fragment |

| | | | |
|---|---|---|---|
| Isotope correction at $MS^2$ | No | Lipid Identification Software | MS-DIAL |
| $MS^1$ verified by standard | No | Data manipulation | Smoothing, Centroiding |
| $MS^2$ verified by standard | No | Nomenclature for intact lipid molecule | Yes |
| Background check at $MS^1$ | Yes | Nomenclature for fragment ions | No |
| Background check at $MS^2$ | No | | |

#### 84) LPA / Lipid quantification

| | | | |
|---|---|---|---|
| Quantitative | Yes | Type I isotope correction | No |
| MS Level for quantification | $MS^1$ | Limit of quantification | No |
| Internal lipid standard(s) $MS^1$ | | Normalization to reference | No |

| Internal standard | Endogenous subclass |
|---|---|
| LPC 18:1(d7) | LPA subclass |

| | | | |
|---|---|---|---|
| Type of quantification | Internal standard amount | Lipid Quantification Software | MS-DIAL |
| Response correction | No | Batch correction | No |

#### 85) LPE / Lipid identification

| | | | |
|---|---|---|---|
| Lipid class | LPE | Did you presume assumptions for identification? | No |
| MS Level for identification | $MS^1$, $MS^2$ | Limit of detection | No |
| Identification level | Molecular species level | RT verified by standard | Yes |
| Isotope correction at $MS^1$ | No | Separation of isobaric/isomeric interferece confirmed | Yes |
| Fragments for identification | | Model for separation prediction | Yes |

| Fragment name |
|---|
| Neutral loss of C2H8NO4P |

| | | | |
|---|---|---|---|
| Isotope correction at $MS^2$ | No | Lipid Identification Software | MS-DIAL |
| $MS^1$ verified by standard | Yes | Data manipulation | Smoothing, Centroiding |
| $MS^2$ verified by standard | Yes | Nomenclature for intact lipid molecule | Yes |
| Background check at $MS^1$ | Yes | Nomenclature for fragment ions | No |
| Background check at $MS^2$ | No | | |

#### 85) LPE / Lipid quantification

| | | | |
|---|---|---|---|
| Quantitative | Yes | Type I isotope correction | No |
| MS Level for quantification | $MS^1$ | Limit of quantification | No |
| Internal lipid standard(s) $MS^1$ | | Normalization to reference | No |

| Internal standard | Endogenous subclass |
|---|---|
| LPE 18:1(d7) | LPE subclass |

| | | | |
|---|---|---|---|
| Type of quantification | Internal standard amount | Lipid Quantification Software | MS-DIAL |
| Response correction | No | Batch correction | No |

#### 86) LPG / Lipid identification

| | | | |
|---|---|---|---|
| Lipid class | LPG | Did you presume assumptions for identification? | No |
| MS Level for identification | $MS^1$, $MS^2$ | Limit of detection | No |
| Identification level | Molecular species level | RT verified by standard | Yes |
| Isotope correction at $MS^1$ | No | Separation of isobaric/isomeric interferece confirmed | Yes |
| Fragments for identification | | Model for separation prediction | Yes |

| Fragment name |
|---|
| Phosphoglycerol -H2O fragment |

| | | | |
|---|---|---|---|
| Isotope correction at $MS^2$ | No | Lipid Identification Software | MS-DIAL |
| $MS^1$ verified by standard | No | Data manipulation | Smoothing, Centroiding |
| $MS^2$ verified by standard | No | Nomenclature for intact lipid molecule | Yes |
| Background check at $MS^1$ | Yes | Nomenclature for fragment ions | No |
| Background check at $MS^2$ | No | | |

#### 86) LPG / Lipid quantification

| | | | |
|---|---|---|---|
| Quantitative | Yes | Type I isotope correction | No |
| MS Level for quantification | $MS^1$ | Limit of quantification | No |
| Internal lipid standard(s) $MS^1$ | | Normalization to reference | No |

| Internal standard | Endogenous subclass |
|---|---|
| PG 15:0_18:1(d7) | LPG subclass |

| | | | |
|---|---|---|---|
| Type of quantification | Internal standard amount | Lipid Quantification Software | MS-DIAL |
| Response correction | No | Batch correction | No |

#### 87) LPI / Lipid identification

| | | | |
|---|---|---|---|
| Lipid class | LPI | Did you presume assumptions for identification? | No |
| MS Level for identification | $MS^1$, $MS^2$ | Limit of detection | No |
| Identification level | Molecular species level | RT verified by standard | Yes |
| Isotope correction at $MS^1$ | No | Separation of isobaric/isomeric interferece confirmed | Yes |
| Fragments for identification | | Model for separation prediction | Yes |

| Fragment name |
|---|
| Phosphoinositol -H2O fragment |
| Characteristic fragment (C9H16O10P-) |
| Fatty acid fragment |

| | | | |
|---|---|---|---|
| Isotope correction at $MS^2$ | No | Lipid Identification Software | MS-DIAL |
| $MS^1$ verified by standard | No | Data manipulation | Smoothing, Centroiding |
| $MS^2$ verified by standard | No | Nomenclature for intact lipid molecule | Yes |
| Background check at $MS^1$ | Yes | Nomenclature for fragment ions | No |
| Background check at $MS^2$ | No | | |

#### 87) LPI / Lipid quantification

| | | | |
|---|---|---|---|
| Quantitative | Yes | Type I isotope correction | No |
| MS Level for quantification | $MS^1$ | Limit of quantification | No |
| Internal lipid standard(s) $MS^1$ | | Normalization to reference | No |

| Internal standard | Endogenous subclass |
|---|---|
| PI 15:0_18:1(d7) | LPI subclass |

| | | | |
|---|---|---|---|
| Type of quantification | Internal standard amount | Lipid Quantification Software | MS-DIAL |
| Response correction | No | Batch correction | No |

#### 88) LPS / Lipid identification

| | | | |
|---|---|---|---|
| Lipid class | LPS | Did you presume assumptions for identification? | No |
| MS Level for identification | $MS^1$, $MS^2$ | Limit of detection | No |
| Identification level | Molecular species level | RT verified by standard | Yes |
| Isotope correction at $MS^1$ | No | Separation of isobaric/isomeric interferece confirmed | Yes |
| Fragments for identification | | Model for separation prediction | Yes |

| Fragment name |
|---|
| Neutral loss of C3H6NO2 |
| Phosphoglycerol -H2O fragment |

| | | | |
|---|---|---|---|
| Isotope correction at $MS^2$ | No | Lipid Identification Software | MS-DIAL |
| $MS^1$ verified by standard | No | Data manipulation | Smoothing, Centroiding |
| $MS^2$ verified by standard | No | Nomenclature for intact lipid molecule | Yes |
| Background check at $MS^1$ | Yes | Nomenclature for fragment ions | No |
| Background check at $MS^2$ | No | | |

#### 88) LPS / Lipid quantification

| | | | |
|---|---|---|---|
| Quantitative | Yes | Type I isotope correction | No |
| MS Level for quantification | $MS^1$ | Limit of quantification | No |
| Internal lipid standard(s) $MS^1$ | | Normalization to reference | No |

| Internal standard | Endogenous subclass |
|---|---|
| PS 15:0_18:1(d7) | LPS subclass |

| | | | |
|---|---|---|---|
| Type of quantification | Internal standard amount | Lipid Quantification Software | MS-DIAL |
| Response correction | No | Batch correction | No |

#### 89) MIPC / Lipid identification

| | | | |
|---|---|---|---|
| Lipid class | MIPC | Did you presume assumptions for identification? | No |
| MS Level for identification | $MS^1$, $MS^2$ | Limit of detection | No |
| Identification level | Molecular species level | RT verified by standard | Yes |
| Isotope correction at $MS^1$ | No | Separation of isobaric/isomeric interferece confirmed | Yes |
| Fragments for identification | | Model for separation prediction | Yes |

| Fragment name |
|---|
| Characteristic fragment (C12H22O14P-) |
| Phytosphingosine -C2H7NO fragment |

| | | | |
|---|---|---|---|
| Isotope correction at $MS^2$ | No | Lipid Identification Software | MS-DIAL |
| $MS^1$ verified by standard | No | Data manipulation | Smoothing, Centroiding |
| $MS^2$ verified by standard | No | Nomenclature for intact lipid molecule | Yes |
| Background check at $MS^1$ | Yes | Nomenclature for fragment ions | No |
| Background check at $MS^2$ | No | | |

#### 89) MIPC / Lipid quantification

| | | | |
|---|---|---|---|
| Quantitative | Yes | Type I isotope correction | No |
| MS Level for quantification | $MS^1$ | Limit of quantification | No |
| Internal lipid standard(s) $MS^1$ | | Normalization to reference | No |

| Internal standard | Endogenous subclass |
|---|---|
| PI 15:0_18:1(d7) | MIPC subclass |

| | | | |
|---|---|---|---|
| Type of quantification | Internal standard amount | Lipid Quantification Software | MS-DIAL |
| Response correction | No | Batch correction | No |

#### 90) MG / Lipid identification

| | | | |
|---|---|---|---|
| Lipid class | MG | Did you presume assumptions for identification? | No |
| MS Level for identification | $MS^1$, $MS^2$ | Limit of detection | No |
| Identification level | Molecular species level | RT verified by standard | Yes |
| Isotope correction at $MS^1$ | No | Separation of isobaric/isomeric interferece confirmed | Yes |
| Fragments for identification | | Model for separation prediction | Yes |

| Fragment name |
|---|
| Neutral loss of H2O |

| | | | |
|---|---|---|---|
| Isotope correction at $MS^2$ | No | Lipid Identification Software | MS-DIAL |
| $MS^1$ verified by standard | Yes | Data manipulation | Smoothing, Centroiding |
| $MS^2$ verified by standard | Yes | Nomenclature for intact lipid molecule | Yes |
| Background check at $MS^1$ | Yes | Nomenclature for fragment ions | No |
| Background check at $MS^2$ | No | | |

#### 90) MG / Lipid quantification

| | | | |
|---|---|---|---|
| Quantitative | Yes | Type I isotope correction | No |
| MS Level for quantification | $MS^1$ | Limit of quantification | No |
| Internal lipid standard(s) $MS^1$ | | Normalization to reference | No |

| Internal standard | Endogenous subclass |
|---|---|
| MG 18:1(d7) | MG subclass |

| | | | |
|---|---|---|---|
| Type of quantification | Internal standard amount | Lipid Quantification Software | MS-DIAL |
| Response correction | No | Batch correction | No |

#### 91) MGDG / Lipid identification

| | | | |
|---|---|---|---|
| Lipid class | MGDG | Did you presume assumptions for identification? | No |
| MS Level for identification | $MS^1$, $MS^2$ | Limit of detection | No |
| Identification level | Molecular species level | RT verified by standard | Yes |
| Isotope correction at $MS^1$ | No | Separation of isobaric/isomeric interferece confirmed | Yes |
| Fragments for identification | | Model for separation prediction | Yes |

| Fragment name |
|---|
| Fatty acid fragment |

| | | | |
|---|---|---|---|
| Isotope correction at $MS^2$ | No | Lipid Identification Software | MS-DIAL |
| $MS^1$ verified by standard | No | Data manipulation | Smoothing, Centroiding |
| $MS^2$ verified by standard | No | Nomenclature for intact lipid molecule | Yes |
| Background check at $MS^1$ | Yes | Nomenclature for fragment ions | No |
| Background check at $MS^2$ | No | | |

#### 91) MGDG / Lipid quantification

| | | | |
|---|---|---|---|
| Quantitative | Yes | Type I isotope correction | No |
| MS Level for quantification | $MS^1$ | Limit of quantification | No |
| Internal lipid standard(s) $MS^1$ | | Normalization to reference | No |

| Internal standard | Endogenous subclass |
|---|---|
| LPC 18:1(d7) | MGDG subclass |

| | | | |
|---|---|---|---|
| Type of quantification | Internal standard amount | Lipid Quantification Software | MS-DIAL |
| Response correction | No | Batch correction | No |

#### 92) Monogalactosylmonoacylglycerol (MGMG) / Lipid identification

| | | | |
|---|---|---|---|
| Lipid class | Monogalactosylmonoacylglycerol (MGMG) | Did you presume assumptions for identification? | No |
| MS Level for identification | $MS^1$, $MS^2$ | Limit of detection | No |
| Identification level | Molecular species level | RT verified by standard | Yes |
| Isotope correction at $MS^1$ | No | Separation of isobaric/isomeric interferece confirmed | Yes |
| Fragments for identification | | Model for separation prediction | Yes |

| Fragment name |
|---|
| Fatty acid fragment |

| | | | |
|---|---|---|---|
| Isotope correction at $MS^2$ | No | Lipid Identification Software | MS-DIAL |
| $MS^1$ verified by standard | No | Data manipulation | Smoothing, Centroiding |
| $MS^2$ verified by standard | No | Nomenclature for intact lipid molecule | Yes |
| Background check at $MS^1$ | Yes | Nomenclature for fragment ions | No |
| Background check at $MS^2$ | No | | |

#### 92) Monogalactosylmonoacylglycerol (MGMG) / Lipid quantification

| | | | |
|---|---|---|---|
| Quantitative | Yes | Type I isotope correction | No |
| MS Level for quantification | MS$^1$ | Limit of quantification | No |
| Internal lipid standard(s) MS$^1$ | | Normalization to reference | No |
| Type of quantification | Internal standard amount | Lipid Quantification Software | MS-DIAL |
| Response correction | No | Batch correction | No |

| Internal standard | Endogenous subclass |
|---|---|
| LPC 18:1(d7) | MGMG subclass |

#### 93) N-acyl ethanolamines (NAE) / Lipid identification

| | | | |
|---|---|---|---|
| Lipid class | N-acyl ethanolamines (NAE) | Did you presume assumptions for identification? | No |
| MS Level for identification | MS$^1$, MS$^2$ | Limit of detection | No |
| Identification level | Molecular species level | RT verified by standard | Yes |
| Isotope correction at MS$^1$ | No | Separation of isobaric/isomeric interferece confirmed | Yes |
| Fragments for identification | | Model for separation prediction | Yes |
| Isotope correction at MS$^2$ | No | Lipid Identification Software | MS-DIAL |
| MS$^1$ verified by standard | No | Data manipulation | Smoothing, Centroiding |
| MS$^2$ verified by standard | No | Nomenclature for intact lipid molecule | Yes |
| Background check at MS$^1$ | Yes | Nomenclature for fragment ions | No |
| Background check at MS$^2$ | No | | |

| Fragment name |
|---|
| Neutral loss of 2H |

#### 93) N-acyl ethanolamines (NAE) / Lipid quantification

| | | | |
|---|---|---|---|
| Quantitative | Yes | Type I isotope correction | No |
| MS Level for quantification | MS$^1$ | Limit of quantification | No |
| Internal lipid standard(s) MS$^1$ | | Normalization to reference | No |
| Type of quantification | Internal standard amount | Lipid Quantification Software | MS-DIAL |
| Response correction | No | Batch correction | No |

| Internal standard | Endogenous subclass |
|---|---|
| LPC 18:1(d7) | NAE subclass |

#### 94) N-acyl glycine (NAGly) / Lipid identification

| | | | |
|---|---|---|---|
| Lipid class | N-acyl glycine (NAGly) | Did you presume assumptions for identification? | No |
| MS Level for identification | $MS^1$, $MS^2$ | Limit of detection | No |
| Identification level | Molecular species level | RT verified by standard | Yes |
| Isotope correction at $MS^1$ | No | Separation of isobaric/isomeric interferece confirmed | Yes |
| Fragments for identification | | Model for separation prediction | Yes |
| Isotope correction at $MS^2$ | No | Lipid Identification Software | MS-DIAL |
| $MS^1$ verified by standard | No | Data manipulation | Smoothing, Centroiding |
| $MS^2$ verified by standard | No | Nomenclature for intact lipid molecule | Yes |
| Background check at $MS^1$ | Yes | Nomenclature for fragment ions | No |
| Background check at $MS^2$ | No | | |

| Fragment name |
|---|
| Glycine |
| Fatty acyl fragment |
| Fatty acyl -H2O fragment |
| Neutral loss of Acyl and H2O |

#### 94) N-acyl glycine (NAGly) / Lipid quantification

| | | | |
|---|---|---|---|
| Quantitative | Yes | Type I isotope correction | No |
| MS Level for quantification | $MS^1$ | Limit of quantification | No |
| Internal lipid standard(s) $MS^1$ | | Normalization to reference | No |
| Type of quantification | Internal standard amount | Lipid Quantification Software | MS-DIAL |
| Response correction | No | Batch correction | No |

| Internal standard | Endogenous subclass |
|---|---|
| LPC 18:1(d7) | NAGly subclass |

#### 95) N-acyl glycyl serine (NAGlySer) / Lipid identification

| | | | |
|---|---|---|---|
| Lipid class | N-acyl glycyl serine (NAGlySer) | Did you presume assumptions for identification? | No |
| MS Level for identification | $MS^1$, $MS^2$ | Limit of detection | No |
| Identification level | Molecular species level | RT verified by standard | Yes |
| Isotope correction at $MS^1$ | No | Separation of isobaric/isomeric interferece confirmed | Yes |
| Fragments for identification | | Model for separation prediction | Yes |

| Fragment name |
|---|
| Glycylserine |
| Fatty acyl fragment |
| Serine |
| Neutral loss of Acyl and H2O |
| Acyl Glycine fragment |

| | | | |
|---|---|---|---|
| Isotope correction at $MS^2$ | No | Lipid Identification Software | MS-DIAL |
| $MS^1$ verified by standard | No | Data manipulation | Smoothing, Centroiding |
| $MS^2$ verified by standard | No | Nomenclature for intact lipid molecule | Yes |
| Background check at $MS^1$ | Yes | Nomenclature for fragment ions | No |
| Background check at $MS^2$ | No | | |

#### 95) N-acyl glycyl serine (NAGlySer) / Lipid quantification

| | | | |
|---|---|---|---|
| Quantitative | Yes | Type I isotope correction | No |
| MS Level for quantification | $MS^1$ | Limit of quantification | No |
| Internal lipid standard(s) $MS^1$ | | Normalization to reference | No |

| Internal standard | Endogenous subclass |
|---|---|
| LPC 18:1(d7) | NAGlySer subclass |

| | | | |
|---|---|---|---|
| Type of quantification | Internal standard amount | Lipid Quantification Software | MS-DIAL |
| Response correction | No | Batch correction | No |

#### 96) N-acyl ornithine (NAOrn) / Lipid identification

| | | | |
|---|---|---|---|
| Lipid class | N-acyl ornithine (NAOrn) | Did you presume assumptions for identification? | No |
| MS Level for identification | $MS^1$, $MS^2$ | Limit of detection | No |
| Identification level | Molecular species level | RT verified by standard | Yes |
| Isotope correction at $MS^1$ | No | Separation of isobaric/isomeric interferece confirmed | Yes |
| Fragments for identification | | Model for separation prediction | Yes |

| Fragment name |
|---|
| Ornithine -H2O |
| Characteristic fragment (C4H8N+) |
| Neutral loss of Acyl and H2O |
| Neutral loss of Acyl and 2H2O |
| Fatty acyl -H2O fragment |

| | | | |
|---|---|---|---|
| Isotope correction at $MS^2$ | No | Lipid Identification Software | MS-DIAL |
| $MS^1$ verified by standard | No | Data manipulation | Smoothing, Centroiding |
| $MS^2$ verified by standard | No | Nomenclature for intact lipid molecule | Yes |
| Background check at $MS^1$ | Yes | Nomenclature for fragment ions | No |
| Background check at $MS^2$ | No | | |

#### 96) N-acyl ornithine (NAOrn) / Lipid quantification

| | | | |
|---|---|---|---|
| Quantitative | Yes | Type I isotope correction | No |
| MS Level for quantification | $MS^1$ | Limit of quantification | No |
| Internal lipid standard(s) $MS^1$ | | Normalization to reference | No |

| Internal standard | Endogenous subclass |
|---|---|
| LPC 18:1(d7) | NAOrn subclass |

| | | | |
|---|---|---|---|
| Type of quantification | Internal standard amount | Lipid Quantification Software | MS-DIAL |
| Response correction | No | Batch correction | No |

#### 97) N-acyl-lysophosphatidylethanolamine (LNAPE) / Lipid identification

| | | | |
|---|---|---|---|
| Lipid class | N-acyl-lysophosphatidylethanolamine (LNAPE) | Did you presume assumptions for identification? | No |
| MS Level for identification | $MS^1$, $MS^2$ | Limit of detection | No |
| Identification level | Molecular species level | RT verified by standard | Yes |
| Isotope correction at $MS^1$ | No | Separation of isobaric/isomeric interferece confirmed | Yes |
| Fragments for identification | | Model for separation prediction | Yes |

| Fragment name |
|---|
| Phosphoglycerol -H2O fragment |
| Fatty acid fragment |
| Neutral loss of Acyl |
| Neutral loss of Acyl and H2O |

| | | | |
|---|---|---|---|
| Isotope correction at $MS^2$ | No | Lipid Identification Software | MS-DIAL |
| $MS^1$ verified by standard | No | Data manipulation | Smoothing, Centroiding |
| $MS^2$ verified by standard | No | Nomenclature for intact lipid molecule | Yes |
| Background check at $MS^1$ | Yes | Nomenclature for fragment ions | No |
| Background check at $MS^2$ | No | | |

#### 97) N-acyl-lysophosphatidylethanolamine (LNAPE) / Lipid quantification

| | | | |
|---|---|---|---|
| Quantitative | Yes | Type I isotope correction | No |
| MS Level for quantification | $MS^1$ | Limit of quantification | No |
| Internal lipid standard(s) $MS^1$ | | Normalization to reference | No |

| Internal standard | Endogenous subclass |
|---|---|
| PE 15:0_18:1(d7) | LNAPE subclass |

| | | | |
|---|---|---|---|
| Type of quantification | Internal standard amount | Lipid Quantification Software | MS-DIAL |
| Response correction | No | Batch correction | No |

#### 98) N-acyl-lysophosphatidylserine (LNAPS) / Lipid identification

| | | | |
|---|---|---|---|
| Lipid class | N-acyl-lysophosphatidylserine (LNAPS) | Did you presume assumptions for identification? | No |
| MS Level for identification | $MS^1$, $MS^2$ | Limit of detection | No |
| Identification level | Molecular species level | RT verified by standard | Yes |
| Isotope correction at $MS^1$ | No | Separation of isobaric/isomeric interferece confirmed | Yes |
| Fragments for identification | | Model for separation prediction | Yes |

| Fragment name |
|---|
| Phosphoglycerol -H2O fragment |
| Neutral loss of Acyl and C3H5NO2 |

| | | | |
|---|---|---|---|
| Isotope correction at $MS^2$ | No | Lipid Identification Software | MS-DIAL |
| $MS^1$ verified by standard | No | Data manipulation | Smoothing, Centroiding |
| $MS^2$ verified by standard | No | Nomenclature for intact lipid molecule | Yes |
| Background check at $MS^1$ | Yes | Nomenclature for fragment ions | No |
| Background check at $MS^2$ | No | | |

#### 98) N-acyl-lysophosphatidylserine (LNAPS) / Lipid quantification

| | | | |
|---|---|---|---|
| Quantitative | Yes | Type I isotope correction | No |
| MS Level for quantification | $MS^1$ | Limit of quantification | No |
| Internal lipid standard(s) $MS^1$ | | Normalization to reference | No |

| Internal standard | Endogenous subclass |
|---|---|
| PS 15:0_18:1(d7) | LNAPS subclass |

| | | | |
|---|---|---|---|
| Type of quantification | Internal standard amount | Lipid Quantification Software | MS-DIAL |
| Response correction | No | Batch correction | No |

#### 99) DMPE / Lipid identification

| | | | |
|---|---|---|---|
| Lipid class | DMPE | Did you presume assumptions for identification? | No |
| MS Level for identification | $MS^1$, $MS^2$ | Limit of detection | No |
| Identification level | Molecular species level | RT verified by standard | Yes |
| Isotope correction at $MS^1$ | No | Separation of isobaric/isomeric interferece confirmed | Yes |
| Fragments for identification | | Model for separation prediction | Yes |

| Fragment name |
|---|
| Fatty acid fragment |

| | | | |
|---|---|---|---|
| Isotope correction at $MS^2$ | No | Lipid Identification Software | MS-DIAL |
| $MS^1$ verified by standard | No | Data manipulation | Smoothing, Centroiding |
| $MS^2$ verified by standard | No | Nomenclature for intact lipid molecule | Yes |
| Background check at $MS^1$ | Yes | Nomenclature for fragment ions | No |
| Background check at $MS^2$ | No | | |

#### 99) DMPE / Lipid quantification

| | | | |
|---|---|---|---|
| Quantitative | Yes | Type I isotope correction | No |
| MS Level for quantification | $MS^1$ | Limit of quantification | No |
| Internal lipid standard(s) $MS^1$ | | Normalization to reference | No |

| Internal standard | Endogenous subclass |
|---|---|
| PE 15:0_18:1(d7) | DMPE subclass |

| | | | |
|---|---|---|---|
| Type of quantification | Internal standard amount | Lipid Quantification Software | MS-DIAL |
| Response correction | No | Batch correction | No |

#### 100) NGcGM3 (NGcGM3) / Lipid identification

| | | | |
|---|---|---|---|
| Lipid class | NGcGM3 (NGcGM3) | Did you presume assumptions for identification? | No |
| MS Level for identification | $MS^1$, $MS^2$ | Limit of detection | No |
| Identification level | Molecular species level | RT verified by standard | Yes |
| Isotope correction at $MS^1$ | No | Separation of isobaric/isomeric interferece confirmed | Yes |
| Fragments for identification | | Model for separation prediction | Yes |

| Fragment name |
|---|
| Characteristic fragment (C11H16NO9-) |

| | | | |
|---|---|---|---|
| Isotope correction at $MS^2$ | No | Lipid Identification Software | MS-DIAL |
| $MS^1$ verified by standard | No | Data manipulation | Smoothing, Centroiding |
| $MS^2$ verified by standard | No | Nomenclature for intact lipid molecule | Yes |
| Background check at $MS^1$ | Yes | Nomenclature for fragment ions | No |
| Background check at $MS^2$ | No | | |

#### 100) NGcGM3 (NGcGM3) / Lipid quantification

| | | | |
|---|---|---|---|
| Quantitative | Yes | Type I isotope correction | No |
| MS Level for quantification | $MS^1$ | Limit of quantification | No |
| Internal lipid standard(s) $MS^1$ | | Normalization to reference | No |

| Internal standard | Endogenous subclass |
|---|---|
| LPC 18:1(d7) | NGcGM3 subclass |

| | | | |
|---|---|---|---|
| Type of quantification | Internal standard amount | Lipid Quantification Software | MS-DIAL |
| Response correction | No | Batch correction | No |

#### 101) MMPE / Lipid identification

| | | | |
|---|---|---|---|
| Lipid class | MMPE | Did you presume assumptions for identification? | No |
| MS Level for identification | $MS^1$, $MS^2$ | Limit of detection | No |
| Identification level | Molecular species level | RT verified by standard | Yes |
| Isotope correction at $MS^1$ | No | Separation of isobaric/isomeric interferece confirmed | Yes |
| Fragments for identification | | Model for separation prediction | Yes |
| Isotope correction at $MS^2$ | No | Lipid Identification Software | MS-DIAL |
| $MS^1$ verified by standard | No | Data manipulation | Smoothing, Centroiding |
| $MS^2$ verified by standard | No | Nomenclature for intact lipid molecule | Yes |
| Background check at $MS^1$ | Yes | Nomenclature for fragment ions | No |
| Background check at $MS^2$ | No | | |

| Fragment name |
|---|
| Fatty acid fragment |

#### 101) MMPE / Lipid quantification

| | | | |
|---|---|---|---|
| Quantitative | Yes | Type I isotope correction | No |
| MS Level for quantification | $MS^1$ | Limit of quantification | No |
| Internal lipid standard(s) $MS^1$ | | Normalization to reference | No |
| Type of quantification | Internal standard amount | Lipid Quantification Software | MS-DIAL |
| Response correction | No | Batch correction | No |

| Internal standard | Endogenous subclass |
|---|---|
| PE 15:0_18:1(d7) | MMPE subclass |

#### 102) Oxidized fatty acid (OxFA) / Lipid identification

| | | | |
|---|---|---|---|
| Lipid class | Oxidized fatty acid (OxFA) | Did you presume assumptions for identification? | No |
| MS Level for identification | $MS^1$, $MS^2$ | Limit of detection | No |
| Identification level | Species level | RT verified by standard | Yes |
| Isotope correction at $MS^1$ | No | Separation of isobaric/isomeric interferece confirmed | Yes |
| Fragments for identification | | Model for separation prediction | Yes |
| Isotope correction at $MS^2$ | No | Lipid Identification Software | MS-DIAL |
| $MS^1$ verified by standard | No | Data manipulation | Smoothing, Centroiding |
| $MS^2$ verified by standard | No | Nomenclature for intact lipid molecule | Yes |
| Background check at $MS^1$ | Yes | Nomenclature for fragment ions | No |
| Background check at $MS^2$ | No | | |

| Fragment name |
|---|
| Neutral loss of H2O |

#### 102) Oxidized fatty acid (OxFA) / Lipid quantification

| | | | |
|---|---|---|---|
| Quantitative | Yes | Type I isotope correction | No |
| MS Level for quantification | $MS^1$ | Limit of quantification | No |
| Internal lipid standard(s) $MS^1$ | | Normalization to reference | No |

| Internal standard | Endogenous subclass |
|---|---|
| FA 18:0(d3) | OxFA subclass |

| | | | |
|---|---|---|---|
| Type of quantification | Internal standard amount | Lipid Quantification Software | MS-DIAL |
| Response correction | No | Batch correction | No |

#### 103) Oxidized phosphatidylcholine (OxPC) / Lipid identification

| | | | |
|---|---|---|---|
| Lipid class | Oxidized phosphatidylcholine (OxPC) | Did you presume assumptions for identification? | No |
| MS Level for identification | $MS^1$, $MS^2$ | Limit of detection | No |
| Identification level | Molecular species level | RT verified by standard | Yes |
| Isotope correction at $MS^1$ | No | Separation of isobaric/isomeric interferece confirmed | Yes |
| Fragments for identification | | Model for separation prediction | Yes |

| Fragment name |
|---|
| Neutral loss of methyl moiety |
| Fatty acid fragment |
| Oxidized fatty acid fragment |
| Oxidized fatty acid -H2O fragment |

| | | | |
|---|---|---|---|
| Isotope correction at $MS^2$ | No | Lipid Identification Software | MS-DIAL |
| $MS^1$ verified by standard | No | Data manipulation | Smoothing, Centroiding |
| $MS^2$ verified by standard | No | Nomenclature for intact lipid molecule | Yes |
| Background check at $MS^1$ | Yes | Nomenclature for fragment ions | No |
| Background check at $MS^2$ | No | | |

#### 103) Oxidized phosphatidylcholine (OxPC) / Lipid quantification

| | | | |
|---|---|---|---|
| Quantitative | Yes | Type I isotope correction | No |
| MS Level for quantification | $MS^1$ | Limit of quantification | No |
| Internal lipid standard(s) $MS^1$ | | Normalization to reference | No |

| Internal standard | Endogenous subclass |
|---|---|
| PC 15:0_18:1(d7) | OxPC subclass |

| | | | |
|---|---|---|---|
| Type of quantification | Internal standard amount | Lipid Quantification Software | MS-DIAL |
| Response correction | No | Batch correction | No |

#### 104) Oxidized phosphatidylethanolamine (OxPE) / Lipid identification

| | | | |
|---|---|---|---|
| Lipid class | Oxidized phosphatidylethanolamine (OxPE) | Did you presume assumptions for identification? | No |
| MS Level for identification | $MS^1$, $MS^2$ | Limit of detection | No |
| Identification level | Molecular species level | RT verified by standard | Yes |
| Isotope correction at $MS^1$ | No | Separation of isobaric/isomeric interferece confirmed | Yes |
| Fragments for identification | | Model for separation prediction | Yes |

| Fragment name |
|---|
| Neutral loss of H2O |
| Fatty acid fragment |
| Oxidized fatty acid fragment |
| Oxidized fatty acid -H2O fragment |

| | | | |
|---|---|---|---|
| Isotope correction at $MS^2$ | No | Lipid Identification Software | MS-DIAL |
| $MS^1$ verified by standard | No | Data manipulation | Smoothing, Centroiding |
| $MS^2$ verified by standard | No | Nomenclature for intact lipid molecule | Yes |
| Background check at $MS^1$ | Yes | Nomenclature for fragment ions | No |
| Background check at $MS^2$ | No | | |

#### 104) Oxidized phosphatidylethanolamine (OxPE) / Lipid quantification

| | | | |
|---|---|---|---|
| Quantitative | Yes | Type I isotope correction | No |
| MS Level for quantification | $MS^1$ | Limit of quantification | No |
| Internal lipid standard(s) $MS^1$ | | Normalization to reference | No |

| Internal standard | Endogenous subclass |
|---|---|
| PE 15:0_18:1(d7) | OxPE subclass |

| | | | |
|---|---|---|---|
| Type of quantification | Internal standard amount | Lipid Quantification Software | MS-DIAL |
| Response correction | No | Batch correction | No |

#### 105) Oxidized phosphatidylglycerol (OxPG) / Lipid identification

| | | | |
|---|---|---|---|
| Lipid class | Oxidized phosphatidylglycerol (OxPG) | Did you presume assumptions for identification? | No |
| MS Level for identification | $MS^1$, $MS^2$ | Limit of detection | No |
| Identification level | Molecular species level | RT verified by standard | Yes |
| Isotope correction at $MS^1$ | No | Separation of isobaric/isomeric interferece confirmed | Yes |
| Fragments for identification | | Model for separation prediction | Yes |

| Fragment name |
|---|
| Fatty acid fragment |
| Oxidized fatty acid fragment |
| Oxidized fatty acid -H2O fragment |

| | | | |
|---|---|---|---|
| Isotope correction at $MS^2$ | No | Lipid Identification Software | MS-DIAL |
| $MS^1$ verified by standard | No | Data manipulation | Smoothing, Centroiding |
| $MS^2$ verified by standard | No | Nomenclature for intact lipid molecule | Yes |
| Background check at $MS^1$ | Yes | Nomenclature for fragment ions | No |
| Background check at $MS^2$ | No | | |

#### 105) Oxidized phosphatidylglycerol (OxPG) / Lipid quantification

| | | | |
|---|---|---|---|
| Quantitative | Yes | Type I isotope correction | No |
| MS Level for quantification | $MS^1$ | Limit of quantification | No |
| Internal lipid standard(s) $MS^1$ | | Normalization to reference | No |

| Internal standard | Endogenous subclass |
|---|---|
| PG 15:0_18:1(d7) | OxPG subclass |

| | | | |
|---|---|---|---|
| Type of quantification | Internal standard amount | Lipid Quantification Software | MS-DIAL |
| Response correction | No | Batch correction | No |

#### 106) Oxidized phosphatidylinositol (OxPI) / Lipid identification

| | | | |
|---|---|---|---|
| Lipid class | Oxidized phosphatidylinositol (OxPI) | Did you presume assumptions for identification? | No |
| MS Level for identification | $MS^1$, $MS^2$ | Limit of detection | No |
| Identification level | Molecular species level | RT verified by standard | Yes |
| Isotope correction at $MS^1$ | No | Separation of isobaric/isomeric interferece confirmed | Yes |
| Fragments for identification | | Model for separation prediction | Yes |

| Fragment name |
|---|
| Phosphoinositol -H2O fragment |
| Characteristic fragment (C9H14O9P-) |
| Neutral loss of H2O |
| Fatty acid fragment |
| Oxidized fatty acid fragment |
| Oxidized fatty acid -H2O fragment |

| | | | |
|---|---|---|---|
| Isotope correction at $MS^2$ | No | Lipid Identification Software | MS-DIAL |
| $MS^1$ verified by standard | No | Data manipulation | Smoothing, Centroiding |
| $MS^2$ verified by standard | No | Nomenclature for intact lipid molecule | Yes |
| Background check at $MS^1$ | Yes | Nomenclature for fragment ions | No |
| Background check at $MS^2$ | No | | |

#### 106) Oxidized phosphatidylinositol (OxPI) / Lipid quantification

| | | | |
|---|---|---|---|
| Quantitative | Yes | Type I isotope correction | No |
| MS Level for quantification | $MS^1$ | Limit of quantification | No |
| Internal lipid standard(s) $MS^1$ | | Normalization to reference | No |

| Internal standard | Endogenous subclass |
|---|---|
| PI 15:0_18:1(d7) | OxPI subclass |

| | | | |
|---|---|---|---|
| Type of quantification | Internal standard amount | Lipid Quantification Software | MS-DIAL |
| Response correction | No | Batch correction | No |

#### 107) Oxidized phosphatidylserine (OxPS) / Lipid identification

| | | | |
|---|---|---|---|
| Lipid class | Oxidized phosphatidylserine (OxPS) | Did you presume assumptions for identification? | No |
| MS Level for identification | $MS^1$, $MS^2$ | Limit of detection | No |
| Identification level | Molecular species level | RT verified by standard | Yes |
| Isotope correction at $MS^1$ | No | Separation of isobaric/isomeric interferece confirmed | Yes |
| Fragments for identification | | Model for separation prediction | Yes |

| Fragment name |
|---|
| Neutral loss of C3H6NO2 |
| Neutral loss of H2O |
| Fatty acid fragment |
| Oxidized fatty acid fragment |
| Oxidized fatty acid -H2O fragment |

| | | | |
|---|---|---|---|
| Isotope correction at $MS^2$ | No | Lipid Identification Software | MS-DIAL |
| $MS^1$ verified by standard | No | Data manipulation | Smoothing, Centroiding |
| $MS^2$ verified by standard | No | Nomenclature for intact lipid molecule | Yes |
| Background check at $MS^1$ | Yes | Nomenclature for fragment ions | No |
| Background check at $MS^2$ | No | | |

#### 107) Oxidized phosphatidylserine (OxPS) / Lipid quantification

| | | | |
|---|---|---|---|
| Quantitative | Yes | Type I isotope correction | No |
| MS Level for quantification | $MS^1$ | Limit of quantification | No |
| Internal lipid standard(s) $MS^1$ | | Normalization to reference | No |

| Internal standard | Endogenous subclass |
|---|---|
| PS 15:0_18:1(d7) | OxPS subclass |

| | | | |
|---|---|---|---|
| Type of quantification | Internal standard amount | Lipid Quantification Software | MS-DIAL |
| Response correction | No | Batch correction | No |

#### 108) Oxidized triglyceride (OxTG) / Lipid identification

| | | | |
|---|---|---|---|
| Lipid class | Oxidized triglyceride (OxTG) | Did you presume assumptions for identification? | No |
| MS Level for identification | $MS^1$, $MS^2$ | Limit of detection | No |
| Identification level | Molecular species level | RT verified by standard | Yes |
| Isotope correction at $MS^1$ | No | Separation of isobaric/isomeric interferece confirmed | Yes |
| Fragments for identification | | Model for separation prediction | Yes |

| Fragment name |
|---|
| Neutral loss of acyl and H2O |
| Neutral loss of acyl and 2H2O |
| Neutral loss of acyl and H2O and O |

| | | | |
|---|---|---|---|
| Isotope correction at $MS^2$ | No | Lipid Identification Software | MS-DIAL |
| $MS^1$ verified by standard | No | Data manipulation | Smoothing, Centroiding |
| $MS^2$ verified by standard | No | Nomenclature for intact lipid molecule | Yes |
| Background check at $MS^1$ | Yes | Nomenclature for fragment ions | No |
| Background check at $MS^2$ | No | | |

#### 108) Oxidized triglyceride (OxTG) / Lipid quantification

| | | | |
|---|---|---|---|
| Quantitative | Yes | Type I isotope correction | No |
| MS Level for quantification | $MS^1$ | Limit of quantification | No |
| Internal lipid standard(s) $MS^1$ | | Normalization to reference | No |

| Internal standard | Endogenous subclass |
|---|---|
| TG 15:0_18:1(d7)_15:0 | OxTG subclass |

| | | | |
|---|---|---|---|
| Type of quantification | Internal standard amount | Lipid Quantification Software | MS-DIAL |
| Response correction | No | Batch correction | No |

#### 109) PA / Lipid identification

| | | | |
|---|---|---|---|
| Lipid class | PA | Did you presume assumptions for identification? | No |
| MS Level for identification | $MS^1$, $MS^2$ | Limit of detection | No |
| Identification level | Molecular species level | RT verified by standard | Yes |
| Isotope correction at $MS^1$ | No | Separation of isobaric/isomeric interferece confirmed | Yes |
| Fragments for identification | | Model for separation prediction | Yes |

| Fragment name |
|---|
| Phosphoglycerol -H2O fragment |
| Fatty acid fragment |

| | | | |
|---|---|---|---|
| Isotope correction at $MS^2$ | No | Lipid Identification Software | MS-DIAL |
| $MS^1$ verified by standard | No | Data manipulation | Smoothing, Centroiding |
| $MS^2$ verified by standard | No | Nomenclature for intact lipid molecule | Yes |
| Background check at $MS^1$ | Yes | Nomenclature for fragment ions | No |
| Background check at $MS^2$ | No | | |

#### 109) PA / Lipid quantification

| | | | |
|---|---|---|---|
| Quantitative | Yes | Type I isotope correction | No |
| MS Level for quantification | $MS^1$ | Limit of quantification | No |
| Internal lipid standard(s) $MS^1$ | | Normalization to reference | No |

| Internal standard | Endogenous subclass |
|---|---|
| LPC 18:1(d7) | PA subclass |

| | | | |
|---|---|---|---|
| Type of quantification | Internal standard amount | Lipid Quantification Software | MS-DIAL |
| Response correction | No | Batch correction | No |

#### 110) PC / Lipid identification

| | | | |
|---|---|---|---|
| Lipid class | PC | Did you presume assumptions for identification? | No |
| MS Level for identification | $MS^1$, $MS^2$ | Limit of detection | No |
| Identification level | Molecular species level | RT verified by standard | Yes |
| Isotope correction at $MS^1$ | No | Separation of isobaric/isomeric interferece confirmed | Yes |
| Fragments for identification | | Model for separation prediction | Yes |

| Fragment name |
|---|
| Neutral loss of methyl moiety |
| Fatty acid fragment |

| | | | |
|---|---|---|---|
| Isotope correction at $MS^2$ | No | Lipid Identification Software | MS-DIAL |
| $MS^1$ verified by standard | Yes | Data manipulation | Smoothing, Centroiding |
| $MS^2$ verified by standard | Yes | Nomenclature for intact lipid molecule | Yes |
| Background check at $MS^1$ | Yes | Nomenclature for fragment ions | No |
| Background check at $MS^2$ | No | | |

#### 110) PC / Lipid quantification

| | | | |
|---|---|---|---|
| Quantitative | Yes | Type I isotope correction | No |
| MS Level for quantification | $MS^1$ | Limit of quantification | No |
| Internal lipid standard(s) $MS^1$ | | Normalization to reference | No |

| Internal standard | Endogenous subclass |
|---|---|
| PC 15:0_18:1(d7) | PC subclass |

| | | | |
|---|---|---|---|
| Type of quantification | Internal standard amount | Lipid Quantification Software | MS-DIAL |
| Response correction | No | Batch correction | No |

#### 111) Phosphatidylethanol (PEtOH) / Lipid identification

| | | | |
|---|---|---|---|
| Lipid class | Phosphatidylethanol (PEtOH) | Did you presume assumptions for identification? | No |
| MS Level for identification | $MS^1$, $MS^2$ | Limit of detection | No |
| Identification level | Molecular species level | RT verified by standard | Yes |
| Isotope correction at $MS^1$ | No | Separation of isobaric/isomeric interferece confirmed | Yes |
| Fragments for identification | | Model for separation prediction | Yes |

| Fragment name |
|---|
| Phosphoethanol |
| Fatty acid fragment |

| | | | |
|---|---|---|---|
| Isotope correction at $MS^2$ | No | Lipid Identification Software | MS-DIAL |
| $MS^1$ verified by standard | No | Data manipulation | Smoothing, Centroiding |
| $MS^2$ verified by standard | No | Nomenclature for intact lipid molecule | Yes |
| Background check at $MS^1$ | Yes | Nomenclature for fragment ions | No |
| Background check at $MS^2$ | No | | |

#### 111) Phosphatidylethanol (PEtOH) / Lipid quantification

| | | | |
|---|---|---|---|
| Quantitative | Yes | Type I isotope correction | No |
| MS Level for quantification | $MS^1$ | Limit of quantification | No |
| Internal lipid standard(s) $MS^1$ | | Normalization to reference | No |

| Internal standard | Endogenous subclass |
|---|---|
| LPC 18:1(d7) | PEtOH subclass |

| | | | |
|---|---|---|---|
| Type of quantification | Internal standard amount | Lipid Quantification Software | MS-DIAL |
| Response correction | No | Batch correction | No |

#### 112) PE / Lipid identification

| | | | |
|---|---|---|---|
| Lipid class | PE | Did you presume assumptions for identification? | No |
| MS Level for identification | $MS^1$, $MS^2$ | Limit of detection | No |
| Identification level | Molecular species level | RT verified by standard | Yes |
| Isotope correction at $MS^1$ | No | Separation of isobaric/isomeric interferece confirmed | Yes |
| Fragments for identification | | Model for separation prediction | Yes |

| Fragment name |
|---|
| Characteristic fragment (C5H11NO5P-) |
| Fatty acid fragment |

| | | | |
|---|---|---|---|
| Isotope correction at $MS^2$ | No | Lipid Identification Software | MS-DIAL |
| $MS^1$ verified by standard | Yes | Data manipulation | Smoothing, Centroiding |
| $MS^2$ verified by standard | Yes | Nomenclature for intact lipid molecule | Yes |
| Background check at $MS^1$ | Yes | Nomenclature for fragment ions | No |
| Background check at $MS^2$ | No | | |

#### 112) PE / Lipid quantification

| | | | |
|---|---|---|---|
| Quantitative | Yes | Type I isotope correction | No |
| MS Level for quantification | $MS^1$ | Limit of quantification | No |
| Internal lipid standard(s) $MS^1$ | | Normalization to reference | No |

| Internal standard | Endogenous subclass |
|---|---|
| PE 15:0_18:1(d7) | PE subclass |

| | | | |
|---|---|---|---|
| Type of quantification | Internal standard amount | Lipid Quantification Software | MS-DIAL |
| Response correction | No | Batch correction | No |

#### 113) PG / Lipid identification

| | | | |
|---|---|---|---|
| Lipid class | PG | Did you presume assumptions for identification? | No |
| MS Level for identification | $MS^1$, $MS^2$ | Limit of detection | No |
| Identification level | Molecular species level | RT verified by standard | Yes |
| Isotope correction at $MS^1$ | No | Separation of isobaric/isomeric interferece confirmed | Yes |
| Fragments for identification | | Model for separation prediction | Yes |

| Fragment name |
|---|
| Phosphoglycerol -H2O |
| Fatty acid fragment |

| | | | |
|---|---|---|---|
| Isotope correction at $MS^2$ | No | Lipid Identification Software | MS-DIAL |
| $MS^1$ verified by standard | Yes | Data manipulation | Smoothing, Centroiding |
| $MS^2$ verified by standard | Yes | Nomenclature for intact lipid molecule | Yes |
| Background check at $MS^1$ | Yes | Nomenclature for fragment ions | No |
| Background check at $MS^2$ | No | | |

#### 113) PG / Lipid quantification

| | | | |
|---|---|---|---|
| Quantitative | Yes | Type I isotope correction | No |
| MS Level for quantification | $MS^1$ | Limit of quantification | No |
| Internal lipid standard(s) $MS^1$ | | Normalization to reference | No |

| Internal standard | Endogenous subclass |
|---|---|
| PG 15:0_18:1(d7) | PG subclass |

| | | | |
|---|---|---|---|
| Type of quantification | Internal standard amount | Lipid Quantification Software | MS-DIAL |
| Response correction | No | Batch correction | No |

#### 114) PI / Lipid identification

| | | | |
|---|---|---|---|
| Lipid class | PI | Did you presume assumptions for identification? | No |
| MS Level for identification | $MS^1$, $MS^2$ | Limit of detection | No |
| Identification level | Molecular species level | RT verified by standard | Yes |
| Isotope correction at $MS^1$ | No | Separation of isobaric/isomeric interferece confirmed | Yes |
| Fragments for identification | | Model for separation prediction | Yes |

| Fragment name |
|---|
| Phosphoinositol -H2O |
| Characteristic fragment (C9H14O9P-) |
| Fatty acid fragment |

| | | | |
|---|---|---|---|
| Isotope correction at $MS^2$ | No | Lipid Identification Software | MS-DIAL |
| $MS^1$ verified by standard | Yes | Data manipulation | Smoothing, Centroiding |
| $MS^2$ verified by standard | Yes | Nomenclature for intact lipid molecule | Yes |
| Background check at $MS^1$ | Yes | Nomenclature for fragment ions | No |
| Background check at $MS^2$ | No | | |

#### 114) PI / Lipid quantification

| | | | |
|---|---|---|---|
| Quantitative | Yes | Type I isotope correction | No |
| MS Level for quantification | $MS^1$ | Limit of quantification | No |
| Internal lipid standard(s) $MS^1$ | | Normalization to reference | No |

| Internal standard | Endogenous subclass |
|---|---|
| PI 15:0_18:1(d7) | PI subclass |

| | | | |
|---|---|---|---|
| Type of quantification | Internal standard amount | Lipid Quantification Software | MS-DIAL |
| Response correction | No | Batch correction | No |

#### 115) Phosphatidylmethanol (PMeOH) / Lipid identification

| | | | |
|---|---|---|---|
| Lipid class | Phosphatidylmethanol (PMeOH) | Did you presume assumptions for identification? | No |
| MS Level for identification | $MS^1$, $MS^2$ | Limit of detection | No |
| Identification level | Molecular species level | RT verified by standard | Yes |
| Isotope correction at $MS^1$ | No | Separation of isobaric/isomeric interferece confirmed | Yes |
| Fragments for identification | | Model for separation prediction | Yes |

| Fragment name |
|---|
| Phosphomethanol |
| Fatty acid fragment |

| | | | |
|---|---|---|---|
| Isotope correction at $MS^2$ | No | Lipid Identification Software | MS-DIAL |
| $MS^1$ verified by standard | No | Data manipulation | Smoothing, Centroiding |
| $MS^2$ verified by standard | No | Nomenclature for intact lipid molecule | Yes |
| Background check at $MS^1$ | Yes | Nomenclature for fragment ions | No |
| Background check at $MS^2$ | No | | |

#### 115) Phosphatidylmethanol (PMeOH) / Lipid quantification

| | | | |
|---|---|---|---|
| Quantitative | Yes | Type I isotope correction | No |
| MS Level for quantification | $MS^1$ | Limit of quantification | No |
| Internal lipid standard(s) $MS^1$ | | Normalization to reference | No |

| Internal standard | Endogenous subclass |
|---|---|
| LPC 18:1(d7) | PMeOH subclass |

| | | | |
|---|---|---|---|
| Type of quantification | Internal standard amount | Lipid Quantification Software | MS-DIAL |
| Response correction | No | Batch correction | No |

#### 116) PS / Lipid identification

| | | | |
|---|---|---|---|
| Lipid class | PS | Did you presume assumptions for identification? | No |
| MS Level for identification | $MS^1$, $MS^2$ | Limit of detection | No |
| Identification level | Molecular species level | RT verified by standard | Yes |
| Isotope correction at $MS^1$ | No | Separation of isobaric/isomeric interferece confirmed | Yes |
| Fragments for identification | | Model for separation prediction | Yes |

| Fragment name |
|---|
| Neutral loss of C3H6NO2 |
| Fatty acid fragment |

| | | | |
|---|---|---|---|
| Isotope correction at $MS^2$ | No | Lipid Identification Software | MS-DIAL |
| $MS^1$ verified by standard | Yes | Data manipulation | Smoothing, Centroiding |
| $MS^2$ verified by standard | Yes | Nomenclature for intact lipid molecule | Yes |
| Background check at $MS^1$ | Yes | Nomenclature for fragment ions | No |
| Background check at $MS^2$ | No | | |

#### 116) PS / Lipid quantification

| | | | |
|---|---|---|---|
| Quantitative | Yes | Type I isotope correction | No |
| MS Level for quantification | $MS^1$ | Limit of quantification | No |
| Internal lipid standard(s) $MS^1$ | | Normalization to reference | No |

| Internal standard | Endogenous subclass |
|---|---|
| PS 15:0_18:1(d7) | PS subclass |

| | | | |
|---|---|---|---|
| Type of quantification | Internal standard amount | Lipid Quantification Software | MS-DIAL |
| Response correction | No | Batch correction | No |

#### 117) Phytosphingosine (PhytoSph) / Lipid identification

| | | | |
|---|---|---|---|
| Lipid class | Phytosphingosine (PhytoSph) | Did you presume assumptions for identification? | No |
| MS Level for identification | $MS^1$, $MS^2$ | Limit of detection | No |
| Identification level | Molecular species level | RT verified by standard | Yes |
| Isotope correction at $MS^1$ | No | Separation of isobaric/isomeric interferece confirmed | Yes |
| Fragments for identification | | Model for separation prediction | Yes |

| Fragment name |
|---|
| Neutral loss of H2O |
| Neutral loss of 2H2O |
| Neutral loss of 3H2O |
| Neutral loss of CH4O2 |

| | | | |
|---|---|---|---|
| Isotope correction at $MS^2$ | No | Lipid Identification Software | MS-DIAL |
| $MS^1$ verified by standard | No | Data manipulation | Smoothing, Centroiding |
| $MS^2$ verified by standard | No | Nomenclature for intact lipid molecule | Yes |
| Background check at $MS^1$ | Yes | Nomenclature for fragment ions | No |
| Background check at $MS^2$ | No | | |

#### 117) Phytosphingosine (PhytoSph) / Lipid quantification

| | | | |
|---|---|---|---|
| Quantitative | Yes | Type I isotope correction | No |
| MS Level for quantification | $MS^1$ | Limit of quantification | No |
| Internal lipid standard(s) $MS^1$ | | Normalization to reference | No |

| Internal standard | Endogenous subclass |
|---|---|
| Cer 18:1;2O/15:0(d7) | PhytoSph subclass |

| | | | |
|---|---|---|---|
| Type of quantification | Internal standard amount | Lipid Quantification Software | MS-DIAL |
| Response correction | No | Batch correction | No |

#### 118) Semino lipid (EtherSMGDG) / Lipid identification

| | | | |
|---|---|---|---|
| Lipid class | Semino lipid (EtherSMGDG) | Did you presume assumptions for identification? | No |
| MS Level for identification | $MS^1$, $MS^2$ | Limit of detection | No |
| Identification level | Molecular species level | RT verified by standard | Yes |
| Isotope correction at $MS^1$ | No | Separation of isobaric/isomeric interferece confirmed | Yes |
| Fragments for identification | | Model for separation prediction | Yes |

| Fragment name |
|---|
| Sulfate |
| Hexosylsulfate |
| Neutral loss of acyl and H2O |

| | | | |
|---|---|---|---|
| Isotope correction at $MS^2$ | No | Lipid Identification Software | MS-DIAL |
| $MS^1$ verified by standard | No | Data manipulation | Smoothing, Centroiding |
| $MS^2$ verified by standard | No | Nomenclature for intact lipid molecule | Yes |
| Background check at $MS^1$ | Yes | Nomenclature for fragment ions | No |
| Background check at $MS^2$ | No | | |

#### 118) Semino lipid (EtherSMGDG) / Lipid quantification

| | | | |
|---|---|---|---|
| Quantitative | Yes | Type I isotope correction | No |
| MS Level for quantification | $MS^1$ | Limit of quantification | No |
| Internal lipid standard(s) $MS^1$ | | Normalization to reference | No |

| Internal standard | Endogenous subclass |
|---|---|
| Cer 18:1;2O/15:0(d7) | EtherSMGDG subclass |

| | | | |
|---|---|---|---|
| Type of quantification | Internal standard amount | Lipid Quantification Software | MS-DIAL |
| Response correction | No | Batch correction | No |

#### 119) Semino lipid (SMGDG) / Lipid identification

| | | | |
|---|---|---|---|
| Lipid class | Semino lipid (SMGDG) | Did you presume assumptions for identification? | No |
| MS Level for identification | $MS^1$, $MS^2$ | Limit of detection | No |
| Identification level | Molecular species level | RT verified by standard | Yes |
| Isotope correction at $MS^1$ | No | Separation of isobaric/isomeric interferece confirmed | Yes |
| Fragments for identification | | Model for separation prediction | Yes |

| Fragment name |
|---|
| Sulfate |
| Hexosylsulfate |
| Neutral loss of acyl and H2O |
| Fatty acid fragment |

| | | | |
|---|---|---|---|
| Isotope correction at $MS^2$ | No | Lipid Identification Software | MS-DIAL |
| $MS^1$ verified by standard | No | Data manipulation | Smoothing, Centroiding |
| $MS^2$ verified by standard | No | Nomenclature for intact lipid molecule | Yes |
| Background check at $MS^1$ | Yes | Nomenclature for fragment ions | No |
| Background check at $MS^2$ | No | | |

#### 119) Semino lipid (SMGDG) / Lipid quantification

| | | | |
|---|---|---|---|
| Quantitative | Yes | Type I isotope correction | No |
| MS Level for quantification | $MS^1$ | Limit of quantification | No |
| Internal lipid standard(s) $MS^1$ | | Normalization to reference | No |

| Internal standard | Endogenous subclass |
|---|---|
| Cer 18:1;2O/15:0(d7) | SMGDG subclass |

| | | | |
|---|---|---|---|
| Type of quantification | Internal standard amount | Lipid Quantification Software | MS-DIAL |
| Response correction | No | Batch correction | No |

#### 120) Sitosterol ester (SISE) / Lipid identification

| | | | |
|---|---|---|---|
| Lipid class | Sitosterol ester (SISE) | Did you presume assumptions for identification? | No |
| MS Level for identification | $MS^1$, $MS^2$ | Limit of detection | No |
| Identification level | Molecular species level | RT verified by standard | Yes |
| Isotope correction at $MS^1$ | No | Separation of isobaric/isomeric interferece confirmed | Yes |
| Fragments for identification | | Model for separation prediction | Yes |
| Isotope correction at $MS^2$ | No | Lipid Identification Software | MS-DIAL |
| $MS^1$ verified by standard | No | Data manipulation | Smoothing, Centroiding |
| $MS^2$ verified by standard | No | Nomenclature for intact lipid molecule | Yes |
| Background check at $MS^1$ | Yes | Nomenclature for fragment ions | No |
| Background check at $MS^2$ | No | | |

| Fragment name |
|---|
| Neutral loss of fatty acid |

#### 120) Sitosterol ester (SISE) / Lipid quantification

| | | | |
|---|---|---|---|
| Quantitative | Yes | Type I isotope correction | No |
| MS Level for quantification | $MS^1$ | Limit of quantification | No |
| Internal lipid standard(s) $MS^1$ | | Normalization to reference | No |
| Type of quantification | Internal standard amount | Lipid Quantification Software | MS-DIAL |
| Response correction | No | Batch correction | No |

| Internal standard | Endogenous subclass |
|---|---|
| CE 18:1(d7) | SISE subclass |

#### 121) Sphinganine (DHSph) / Lipid identification

| | | | |
|---|---|---|---|
| Lipid class | Sphinganine (DHSph) | Did you presume assumptions for identification? | No |
| MS Level for identification | $MS^1$, $MS^2$ | Limit of detection | No |
| Identification level | Molecular species level | RT verified by standard | Yes |
| Isotope correction at $MS^1$ | No | Separation of isobaric/isomeric interferece confirmed | Yes |
| Fragments for identification | | Model for separation prediction | Yes |
| Isotope correction at $MS^2$ | No | Lipid Identification Software | MS-DIAL |
| $MS^1$ verified by standard | No | Data manipulation | Smoothing, Centroiding |
| $MS^2$ verified by standard | No | Nomenclature for intact lipid molecule | Yes |
| Background check at $MS^1$ | Yes | Nomenclature for fragment ions | No |
| Background check at $MS^2$ | No | | |

| Fragment name |
|---|
| Neutral loss of H2O |
| Neutral loss of 2H2O |
| Neutral loss of CH4O2 |

#### 121) Sphinganine (DHSph) / Lipid quantification

| | | | |
|---|---|---|---|
| Quantitative | Yes | Type I isotope correction | No |
| MS Level for quantification | $MS^1$ | Limit of quantification | No |
| Internal lipid standard(s) $MS^1$ | | Normalization to reference | No |

| Internal standard | Endogenous subclass |
|---|---|
| Cer 18:1;2O/15:0(d7) | DHSph subclass |

| | | | |
|---|---|---|---|
| Type of quantification | Internal standard amount | Lipid Quantification Software | MS-DIAL |
| Response correction | No | Batch correction | No |

#### 122) Sphingomyelin (SM) / Lipid identification

| | | | |
|---|---|---|---|
| Lipid class | Sphingomyelin (SM) | Did you presume assumptions for identification? | No |
| MS Level for identification | $MS^1$, $MS^2$ | Limit of detection | No |
| Identification level | Molecular species level | RT verified by standard | Yes |
| Isotope correction at $MS^1$ | No | Separation of isobaric/isomeric interferece confirmed | Yes |
| Fragments for identification | | Model for separation prediction | Yes |

| Fragment name |
|---|
| Neutral loss of methyl moiety |
| Characteristic fragment (C4H11NO4P-) |
| Neutral loss of methyl moiety and acyl |

| | | | |
|---|---|---|---|
| Isotope correction at $MS^2$ | No | Lipid Identification Software | MS-DIAL |
| $MS^1$ verified by standard | Yes | Data manipulation | Smoothing, Centroiding |
| $MS^2$ verified by standard | Yes | Nomenclature for intact lipid molecule | Yes |
| Background check at $MS^1$ | Yes | Nomenclature for fragment ions | No |
| Background check at $MS^2$ | No | | |

#### 122) Sphingomyelin (SM) / Lipid quantification

| | | | |
|---|---|---|---|
| Quantitative | Yes | Type I isotope correction | No |
| MS Level for quantification | $MS^1$ | Limit of quantification | No |
| Internal lipid standard(s) $MS^1$ | | Normalization to reference | No |

| Internal standard | Endogenous subclass |
|---|---|
| SM 18:1;2O/18:1(d9) | SM subclass |

| | | | |
|---|---|---|---|
| Type of quantification | Internal standard amount | Lipid Quantification Software | MS-DIAL |
| Response correction | No | Batch correction | No |

#### 123) SPB / Lipid identification

| | | | |
|---|---|---|---|
| Lipid class | SPB | Did you presume assumptions for identification? | No |
| MS Level for identification | $MS^1$, $MS^2$ | Limit of detection | No |
| Identification level | Molecular species level | RT verified by standard | Yes |
| Isotope correction at $MS^1$ | No | Separation of isobaric/isomeric interferece confirmed | Yes |
| Fragments for identification | | Model for separation prediction | Yes |

| Fragment name |
|---|
| Neutral loss of H2O |
| Neutral loss of 2H2O |
| Neutral loss of CH4O2 |

| | | | |
|---|---|---|---|
| Isotope correction at $MS^2$ | No | Lipid Identification Software | MS-DIAL |
| $MS^1$ verified by standard | No | Data manipulation | Smoothing, Centroiding |
| $MS^2$ verified by standard | No | Nomenclature for intact lipid molecule | Yes |
| Background check at $MS^1$ | Yes | Nomenclature for fragment ions | No |
| Background check at $MS^2$ | No | | |

#### 123) SPB / Lipid quantification

| | | | |
|---|---|---|---|
| Quantitative | Yes | Type I isotope correction | No |
| MS Level for quantification | $MS^1$ | Limit of quantification | No |
| Internal lipid standard(s) $MS^1$ | | Normalization to reference | No |

| Internal standard | Endogenous subclass |
|---|---|
| Cer 18:1;2O/15:0(d7) | Sph subclass |

| | | | |
|---|---|---|---|
| Type of quantification | Internal standard amount | Lipid Quantification Software | MS-DIAL |
| Response correction | No | Batch correction | No |

#### 124) Sterol sulfate (SSulfate) / Lipid identification

| | | | |
|---|---|---|---|
| Lipid class | Sterol sulfate (SSulfate) | Did you presume assumptions for identification? | No |
| MS Level for identification | $MS^1$, $MS^2$ | Limit of detection | No |
| Identification level | Molecular species level | RT verified by standard | Yes |
| Isotope correction at $MS^1$ | No | Separation of isobaric/isomeric interferece confirmed | Yes |
| Fragments for identification | | Model for separation prediction | Yes |

| Fragment name |
|---|
| Cholesterol sulfate |
| Sulfate |

| | | | |
|---|---|---|---|
| Isotope correction at $MS^2$ | No | Lipid Identification Software | MS-DIAL |
| $MS^1$ verified by standard | No | Data manipulation | Smoothing, Centroiding |
| $MS^2$ verified by standard | No | Nomenclature for intact lipid molecule | Yes |
| Background check at $MS^1$ | Yes | Nomenclature for fragment ions | No |
| Background check at $MS^2$ | No | | |

#### 124) Sterol sulfate (SSulfate) / Lipid quantification

| | | | |
|---|---|---|---|
| Quantitative | Yes | Type I isotope correction | No |
| MS Level for quantification | $MS^1$ | Limit of quantification | No |
| Internal lipid standard(s) $MS^1$ | | Normalization to reference | No |

| Internal standard | Endogenous subclass |
|---|---|
| LPC 18:1(d7) | SSulfate subclass |

| | | | |
|---|---|---|---|
| Type of quantification | Internal standard amount | Lipid Quantification Software | MS-DIAL |
| Response correction | No | Batch correction | No |

#### 125) Stigmasterol ester (STSE) / Lipid identification

| | | | |
|---|---|---|---|
| Lipid class | Stigmasterol ester (STSE) | Did you presume assumptions for identification? | No |
| MS Level for identification | $MS^1$, $MS^2$ | Limit of detection | No |
| Identification level | Molecular species level | RT verified by standard | Yes |
| Isotope correction at $MS^1$ | No | Separation of isobaric/isomeric interferece confirmed | Yes |
| Fragments for identification | | Model for separation prediction | Yes |

| Fragment name |
|---|
| Neutral loss of fatty acid |

| | | | |
|---|---|---|---|
| Isotope correction at $MS^2$ | No | Lipid Identification Software | MS-DIAL |
| $MS^1$ verified by standard | No | Data manipulation | Smoothing, Centroiding |
| $MS^2$ verified by standard | No | Nomenclature for intact lipid molecule | Yes |
| Background check at $MS^1$ | Yes | Nomenclature for fragment ions | No |
| Background check at $MS^2$ | No | | |

#### 125) Stigmasterol ester (STSE) / Lipid quantification

| | | | |
|---|---|---|---|
| Quantitative | Yes | Type I isotope correction | No |
| MS Level for quantification | $MS^1$ | Limit of quantification | No |
| Internal lipid standard(s) $MS^1$ | | Normalization to reference | No |

| Internal standard | Endogenous subclass |
|---|---|
| CE 18:1(d7) | STSE subclass |

| | | | |
|---|---|---|---|
| Type of quantification | Internal standard amount | Lipid Quantification Software | MS-DIAL |
| Response correction | No | Batch correction | No |

#### 126) Stigmasterol hexoside (SHex) / Lipid identification

| | | | |
|---|---|---|---|
| Lipid class | Stigmasterol hexoside (SHex) | Did you presume assumptions for identification? | No |
| MS Level for identification | $MS^1$, $MS^2$ | Limit of detection | No |
| Identification level | Molecular species level | RT verified by standard | Yes |
| Isotope correction at $MS^1$ | No | Separation of isobaric/isomeric interferece confirmed | Yes |
| Fragments for identification | | Model for separation prediction | Yes |

| Fragment name |
|---|
| Hexose |

| | | | |
|---|---|---|---|
| Isotope correction at $MS^2$ | No | Lipid Identification Software | MS-DIAL |
| $MS^1$ verified by standard | No | Data manipulation | Smoothing, Centroiding |
| $MS^2$ verified by standard | No | Nomenclature for intact lipid molecule | Yes |
| Background check at $MS^1$ | Yes | Nomenclature for fragment ions | No |
| Background check at $MS^2$ | No | | |

#### 126) Stigmasterol hexoside (SHex) / Lipid quantification

| | | | |
|---|---|---|---|
| Quantitative | Yes | Type I isotope correction | No |
| MS Level for quantification | $MS^1$ | Limit of quantification | No |
| Internal lipid standard(s) $MS^1$ | | Normalization to reference | No |

| Internal standard | Endogenous subclass |
|---|---|
| LPC 18:1(d7) | SHex subclass |

| | | | |
|---|---|---|---|
| Type of quantification | Internal standard amount | Lipid Quantification Software | MS-DIAL |
| Response correction | No | Batch correction | No |

#### 127) SHexCer / Lipid identification

| | | | |
|---|---|---|---|
| Lipid class | SHexCer | Did you presume assumptions for identification? | No |
| MS Level for identification | $MS^1$, $MS^2$ | Limit of detection | No |
| Identification level | Molecular species level | RT verified by standard | Yes |
| Isotope correction at $MS^1$ | No | Separation of isobaric/isomeric interferece confirmed | Yes |
| Fragments for identification | | Model for separation prediction | Yes |

| Fragment name |
|---|
| Neutral loss of sulfate |
| Neutral loss of sulfate and hexose |
| Sphingosine -H2O fragment |
| Sphingosine -2H2O fragment |
| Sphingosine -CH4O2 fragment |

| | | | |
|---|---|---|---|
| Isotope correction at $MS^2$ | No | Lipid Identification Software | MS-DIAL |
| $MS^1$ verified by standard | No | Data manipulation | Smoothing, Centroiding |
| $MS^2$ verified by standard | No | Nomenclature for intact lipid molecule | Yes |
| Background check at $MS^1$ | Yes | Nomenclature for fragment ions | No |
| Background check at $MS^2$ | No | | |

#### 127) SHexCer / Lipid quantification

| | | | |
|---|---|---|---|
| Quantitative | Yes | Type I isotope correction | No |
| MS Level for quantification | $MS^1$ | Limit of quantification | No |
| Internal lipid standard(s) $MS^1$ | | Normalization to reference | No |

| Internal standard | Endogenous subclass |
|---|---|
| Cer 18:1;2O/15:0(d7) | SHexCer subclass |

| | | | |
|---|---|---|---|
| Type of quantification | Internal standard amount | Lipid Quantification Software | MS-DIAL |
| Response correction | No | Batch correction | No |

#### 128) SQDG / Lipid identification

| | | | |
|---|---|---|---|
| Lipid class | SQDG | Did you presume assumptions for identification? | No |
| MS Level for identification | $MS^1$, $MS^2$ | Limit of detection | No |
| Identification level | Molecular species level | RT verified by standard | Yes |
| Isotope correction at $MS^1$ | No | Separation of isobaric/isomeric interferece confirmed | Yes |
| Fragments for identification | | Model for separation prediction | Yes |

| Fragment name |
|---|
| Neutral loss of C6H10O7S |
| Dehydro-monoacyl glycerols |

| | | | |
|---|---|---|---|
| Isotope correction at $MS^2$ | No | Lipid Identification Software | MS-DIAL |
| $MS^1$ verified by standard | No | Data manipulation | Smoothing, Centroiding |
| $MS^2$ verified by standard | No | Nomenclature for intact lipid molecule | Yes |
| Background check at $MS^1$ | Yes | Nomenclature for fragment ions | No |
| Background check at $MS^2$ | No | | |

#### 128) SQDG / Lipid quantification

| | | | |
|---|---|---|---|
| Quantitative | Yes | Type I isotope correction | No |
| MS Level for quantification | $MS^1$ | Limit of quantification | No |
| Internal lipid standard(s) $MS^1$ | | Normalization to reference | No |

| Internal standard | Endogenous subclass |
|---|---|
| LPC 18:1(d7) | SQDG subclass |

| | | | |
|---|---|---|---|
| Type of quantification | Internal standard amount | Lipid Quantification Software | MS-DIAL |
| Response correction | No | Batch correction | No |

#### 129) TG / Lipid identification

| | | | |
|---|---|---|---|
| Lipid class | TG | Did you presume assumptions for identification? | No |
| MS Level for identification | $MS^1$, $MS^2$ | Limit of detection | No |
| Identification level | Molecular species level | RT verified by standard | Yes |
| Isotope correction at $MS^1$ | No | Separation of isobaric/isomeric interferece confirmed | Yes |
| Fragments for identification | | Model for separation prediction | Yes |

| Fragment name |
|---|
| Neutral loss of acyl and H2O |

| | | | |
|---|---|---|---|
| Isotope correction at $MS^2$ | No | Lipid Identification Software | MS-DIAL |
| $MS^1$ verified by standard | Yes | Data manipulation | Smoothing, Centroiding |
| $MS^2$ verified by standard | Yes | Nomenclature for intact lipid molecule | Yes |
| Background check at $MS^1$ | Yes | Nomenclature for fragment ions | No |
| Background check at $MS^2$ | No | | |

#### 129) TG / Lipid quantification

| | | | |
|---|---|---|---|
| Quantitative | Yes | Type I isotope correction | No |
| MS Level for quantification | $MS^1$ | Limit of quantification | No |
| Internal lipid standard(s) $MS^1$ | | Normalization to reference | No |

| Internal standard | Endogenous subclass |
|---|---|
| TG 15:0_18:1(d7)_15:0 | TG subclass |

| | | | |
|---|---|---|---|
| Type of quantification | Internal standard amount | Lipid Quantification Software | MS-DIAL |
| Response correction | No | Batch correction | No |

#### 130) Triacylglycerol estolides (TG_EST) / Lipid identification

| | | | |
|---|---|---|---|
| Lipid class | Triacylglycerol estolides (TG_EST) | Did you presume assumptions for identification? | No |
| MS Level for identification | $MS^1$, $MS^2$ | Limit of detection | No |
| Identification level | Molecular species level | RT verified by standard | Yes |
| Isotope correction at $MS^1$ | No | Separation of isobaric/isomeric interferece confirmed | Yes |
| Fragments for identification | | Model for separation prediction | Yes |

| Fragment name |
|---|
| Neutral loss of acyl and H2O |
| Neutral loss of oxidized acyl and H2O |
| Neutral loss of FAHFA |

| | | | |
|---|---|---|---|
| Isotope correction at $MS^2$ | No | Lipid Identification Software | MS-DIAL |
| $MS^1$ verified by standard | No | Data manipulation | Smoothing, Centroiding |
| $MS^2$ verified by standard | No | Nomenclature for intact lipid molecule | Yes |
| Background check at $MS^1$ | Yes | Nomenclature for fragment ions | No |
| Background check at $MS^2$ | No | | |

#### 130) Triacylglycerol estolides (TG_EST) / Lipid quantification

| | | | |
|---|---|---|---|
| Quantitative | Yes | Type I isotope correction | No |
| MS Level for quantification | $MS^1$ | Limit of quantification | No |
| Internal lipid standard(s) $MS^1$ | | Normalization to reference | No |

| Internal standard | Endogenous subclass |
|---|---|
| TG 15:0_18:1(d7)_15:0 | TG_EST subclass |

| | | | |
|---|---|---|---|
| Type of quantification | Internal standard amount | Lipid Quantification Software | MS-DIAL |
| Response correction | No | Batch correction | No |

#### 131) Hex3Cer / Lipid identification

| | | | |
|---|---|---|---|
| Lipid class | Hex3Cer | Did you presume assumptions for identification? | No |
| MS Level for identification | $MS^1$, $MS^2$ | Limit of detection | No |
| Identification level | Molecular species level | RT verified by standard | Yes |
| Isotope correction at $MS^1$ | No | Separation of isobaric/isomeric interferece confirmed | Yes |
| Fragments for identification | | Model for separation prediction | Yes |

| Fragment name |
|---|
| Neutral loss of hexose |
| Neutral loss of 2hexose |
| Neutral loss of 3hexose |
| Neutral loss of 3hexose and H2O |
| Sphingosine -H2O fragment |
| Sphingosine -2H2O fragment |
| Sphingosine -CH4O2 fragment |

| | | | |
|---|---|---|---|
| Isotope correction at $MS^2$ | No | Lipid Identification Software | MS-DIAL |
| $MS^1$ verified by standard | No | Data manipulation | Smoothing, Centroiding |
| $MS^2$ verified by standard | No | Nomenclature for intact lipid molecule | Yes |
| Background check at $MS^1$ | Yes | Nomenclature for fragment ions | No |
| Background check at $MS^2$ | No | | |

#### 131) Hex3Cer / Lipid quantification

| | | | |
|---|---|---|---|
| Quantitative | Yes | Type I isotope correction | No |
| MS Level for quantification | $MS^1$ | Limit of quantification | No |
| Internal lipid standard(s) $MS^1$ | | Normalization to reference | No |

| Internal standard | Endogenous subclass |
|---|---|
| Cer 18:1;2O/15:0(d7) | Hex3Cer subclass |

| | | | |
|---|---|---|---|
| Type of quantification | Internal standard amount | Lipid Quantification Software | MS-DIAL |
| Response correction | No | Batch correction | No |

#### 132) Vitamin A fatty acid ester (VAE) / Lipid identification

| | | | |
|---|---|---|---|
| Lipid class | Vitamin A fatty acid ester (VAE) | Did you presume assumptions for identification? | No |
| MS Level for identification | $MS^1$, $MS^2$ | Limit of detection | No |
| Identification level | Molecular species level | RT verified by standard | Yes |
| Isotope correction at $MS^1$ | No | Separation of isobaric/isomeric interferece confirmed | Yes |
| Fragments for identification | | Model for separation prediction | Yes |

| Fragment name |
|---|
| Characteristic fragment (C20H29+) |
| Characteristic fragment (C9H11+) |

| | | | |
|---|---|---|---|
| Isotope correction at $MS^2$ | No | Lipid Identification Software | MS-DIAL |
| $MS^1$ verified by standard | No | Data manipulation | Smoothing, Centroiding |
| $MS^2$ verified by standard | No | Nomenclature for intact lipid molecule | Yes |
| Background check at $MS^1$ | Yes | Nomenclature for fragment ions | No |
| Background check at $MS^2$ | No | | |

#### 132) Vitamin A fatty acid ester (VAE) / Lipid quantification

| | | | |
|---|---|---|---|
| Quantitative | Yes | Type I isotope correction | No |
| MS Level for quantification | $MS^1$ | Limit of quantification | No |
| Internal lipid standard(s) $MS^1$ | | Normalization to reference | No |

| Internal standard | Endogenous subclass |
|---|---|
| LPC 18:1(d7) | VAE subclass |

| | | | |
|---|---|---|---|
| Type of quantification | Internal standard amount | Lipid Quantification Software | MS-DIAL |
| Response correction | No | Batch correction | No |

#### 133) Vitamin D / Lipid identification

| | | | |
|---|---|---|---|
| Lipid class | Vitamin D | Did you presume assumptions for identification? | No |
| MS Level for identification | $MS^1$, $MS^2$ | Limit of detection | No |
| Identification level | Species level | RT verified by standard | Yes |
| Isotope correction at $MS^1$ | No | Separation of isobaric/isomeric interferece confirmed | Yes |
| Fragments for identification | | Model for separation prediction | Yes |

| Fragment name |
|---|
| Neutral loss of H2O |
| Neutral loss of 2H2O |

| | | | |
|---|---|---|---|
| Isotope correction at $MS^2$ | No | Lipid Identification Software | MS-DIAL |
| $MS^1$ verified by standard | No | Data manipulation | Smoothing, Centroiding |
| $MS^2$ verified by standard | No | Nomenclature for intact lipid molecule | Yes |
| Background check at $MS^1$ | Yes | Nomenclature for fragment ions | No |
| Background check at $MS^2$ | No | | |

#### 133) Vitamin D / Lipid quantification

| | | | |
|---|---|---|---|
| Quantitative | Yes | Type I isotope correction | No |
| MS Level for quantification | $MS^1$ | Limit of quantification | No |
| Internal lipid standard(s) $MS^1$ | | Normalization to reference | No |

| Internal standard | Endogenous subclass |
|---|---|
| LPC 18:1(d7) | Vitamin D subclass |

| | | | |
|---|---|---|---|
| Type of quantification | Internal standard amount | Lipid Quantification Software | MS-DIAL |
| Response correction | No | Batch correction | No |

#### 134) Vitamin E / Lipid identification

| | | | |
|---|---|---|---|
| Lipid class | Vitamin E | Did you presume assumptions for identification? | No |
| MS Level for identification | $MS^1$, $MS^2$ | Limit of detection | No |
| Identification level | Species level | RT verified by standard | Yes |
| Isotope correction at $MS^1$ | No | Separation of isobaric/isomeric interferece confirmed | Yes |
| Fragments for identification | | Model for separation prediction | Yes |

| Fragment name |
|---|
| Characteristic fragment (C10H11O2-) |

| | | | |
|---|---|---|---|
| Isotope correction at $MS^2$ | No | Lipid Identification Software | MS-DIAL |
| $MS^1$ verified by standard | No | Data manipulation | Smoothing, Centroiding |
| $MS^2$ verified by standard | No | Nomenclature for intact lipid molecule | Yes |
| Background check at $MS^1$ | Yes | Nomenclature for fragment ions | No |
| Background check at $MS^2$ | No | | |

#### 134) Vitamin E / Lipid quantification

| | | | |
|---|---|---|---|
| Quantitative | Yes | Type I isotope correction | No |
| MS Level for quantification | $MS^1$ | Limit of quantification | No |
| Internal lipid standard(s) $MS^1$ | | Normalization to reference | No |

| Internal standard | Endogenous subclass |
|---|---|
| LPC 18:1(d7) | Vitamin E subclass |

| | | | |
|---|---|---|---|
| Type of quantification | Internal standard amount | Lipid Quantification Software | MS-DIAL |
| Response correction | No | Batch correction | No |

#### 135) PE P / Lipid identification

| | | | |
|---|---|---|---|
| Lipid class | PE P | Did you presume assumptions for identification? | No |
| MS Level for identification | $MS^1$, $MS^2$ | Limit of detection | No |
| Identification level | Molecular species level | RT verified by standard | Yes |
| Isotope correction at $MS^1$ | No | Separation of isobaric/isomeric interferece confirmed | Yes |
| Fragments for identification | | Model for separation prediction | Yes |

| Fragment name |
|---|
| Neutral loss of C2H8NO4P |
| Alkyl ether +C2H8NO3P fragment |
| Dehydro-monoacyl glycerols |

| | | | |
|---|---|---|---|
| Isotope correction at $MS^2$ | No | Lipid Identification Software | MS-DIAL |
| $MS^1$ verified by standard | No | Data manipulation | Smoothing, Centroiding |
| $MS^2$ verified by standard | No | Nomenclature for intact lipid molecule | Yes |
| Background check at $MS^1$ | Yes | Nomenclature for fragment ions | No |
| Background check at $MS^2$ | No | | |

#### 135) PE P / Lipid quantification

| | | | |
|---|---|---|---|
| Quantitative | Yes | Type I isotope correction | No |
| MS Level for quantification | $MS^1$ | Limit of quantification | No |
| Internal lipid standard(s) $MS^1$ | | Normalization to reference | No |

| Internal standard | Endogenous subclass |
|---|---|
| PE 15:0_18:1(d7) | EtherPE subclass |

| | | | |
|---|---|---|---|
| Type of quantification | Internal standard amount | Lipid Quantification Software | MS-DIAL |
| Response correction | No | Batch correction | No |
